# Pervasive selective sweeps across human gut microbiomes

**DOI:** 10.1101/2023.12.22.573162

**Authors:** Richard Wolff, Nandita R. Garud

## Abstract

The human gut microbiome is composed of a highly diverse consortia of species which are continually evolving within and across hosts. The ability to identify adaptations common to many human gut microbiomes would not only reveal shared selection pressures across hosts, but also key drivers of functional differentiation of the microbiome that may affect community structure and host traits. However, to date there has not been a systematic scan for adaptations that have spread across human gut microbiomes. Here, we develop a novel selection scan statistic named the integrated Linkage Disequilibrium Score (iLDS) that can detect the spread of adaptive haplotypes across host microbiomes via migration and horizontal gene transfer. Specifically, iLDS leverages signals of hitchhiking of deleterious variants with the beneficial variant. Application of the statistic to *∼*30 of the most prevalent commensal gut species from 24 populations around the world revealed more than 300 selective sweeps across species. We find an enrichment for selective sweeps at loci involved in carbohydrate metabolism—potentially indicative of adaptation to features of host diet—and we find that the targets of selection significantly differ between Westernized and non-Westernized populations. Underscoring the potential role of diet in driving selection, we find a selective sweep absent from non-Westernized populations but ubiquitous in Westernized populations at a locus known to be involved in the metabolism of maltodextrin, a synthetic starch that has recently become a widespread component of Western diets. In summary, we demonstrate that selective sweeps across host microbiomes are a common feature of the evolution of the human gut microbiome, and that targets of selection may be strongly impacted by host diet.

## Introduction

The diverse species that compose the human gut microbiome continually evolve throughout a host’s lifetime. Recent work has shown that rapid adaptation is a hallmark of evolution in the human microbiome, as novel mutations often arise and sweep to high frequency within healthy hosts on timescales of days to months [1, 2, 3, 4, 5, 6, 7]. These evolutionary dynamics can have functional consequences for the host, as microbial genetic variants are associated with numerous traits including metabolic capacity, disease susceptibility, and digestion of food [8, 9, 10, 11].

A novel adaptation which appears initially in one host microbiome may spread across host microbiomes through strain or phage transmission and subsequent horizontal gene transfer (HGT). The human gut microbiome is known to be a hotspot for HGT [12, 13, 14], allowing adaptive alleles to be easily recombined onto new genetic backgrounds. While it has been shown that HGT plays a crucial role in transmission of some genes, such as antibiotic resistance genes, especially across species boundaries, the extent to which HGT facilitates the spread of adaptive alleles across strains of the same species among commensal gut microbiota is at present unclear.

Should an adaptive allele spread between microbiomes in a “gene-specific” selective sweep, the same genomic sequence, or haplotype, surrounding the adaptive allele will appear in many otherwise distantly related strains present in different host microbiomes [12, 15, 16]. Such locally shared haplotypes will result in distinct signatures of elevated linkage disequilibrium (LD), or, correlations among variants that have “hitchhiked” to high frequency with the adaptive allele in the vicinity of the adaptive locus, but not in the surrounding genomic region. While elevations in LD have long been leveraged as a signature of selection in eukaryotes [17, 18, 19, 20, 21, 22], to date LD-based scans for selection in bacteria have been limited [23] and instead HGT-mediated sweeps have largely been discovered on a case-by-case basis [15, 24] rather than by systematic application of established statistics, such as iHS [20]. One reason could be that other evolutionary forces including demographic contractions and reduced recombination rates also result in elevations in LD confounding its use in the discovery of adaptation in bacteria [25, 26, 27, 28].

One way to control for these non-selective forces is to compare LD among synonymous and non-synonymous variants. While both types of variants are subject to the same non-selective forces, synonymous variants are far more likely to be neutral. The vast majority of non-synonymous mutations, by contrast, are deleterious in any population [29], and are always found to be preferentially rare [30, 31]. Hitchhikers that are rare prior to the sweep will exhibit high LD with the adaptive mutation during the sweep as they will typically be found only on haplotypes bearing the adaptive mutation. Therefore, we expect non-synonymous variants to have higher LD than synonymous variants in the vicinity of adaptive loci that have swept to high frequency (**Figure 1A**).

**Figure 1:**
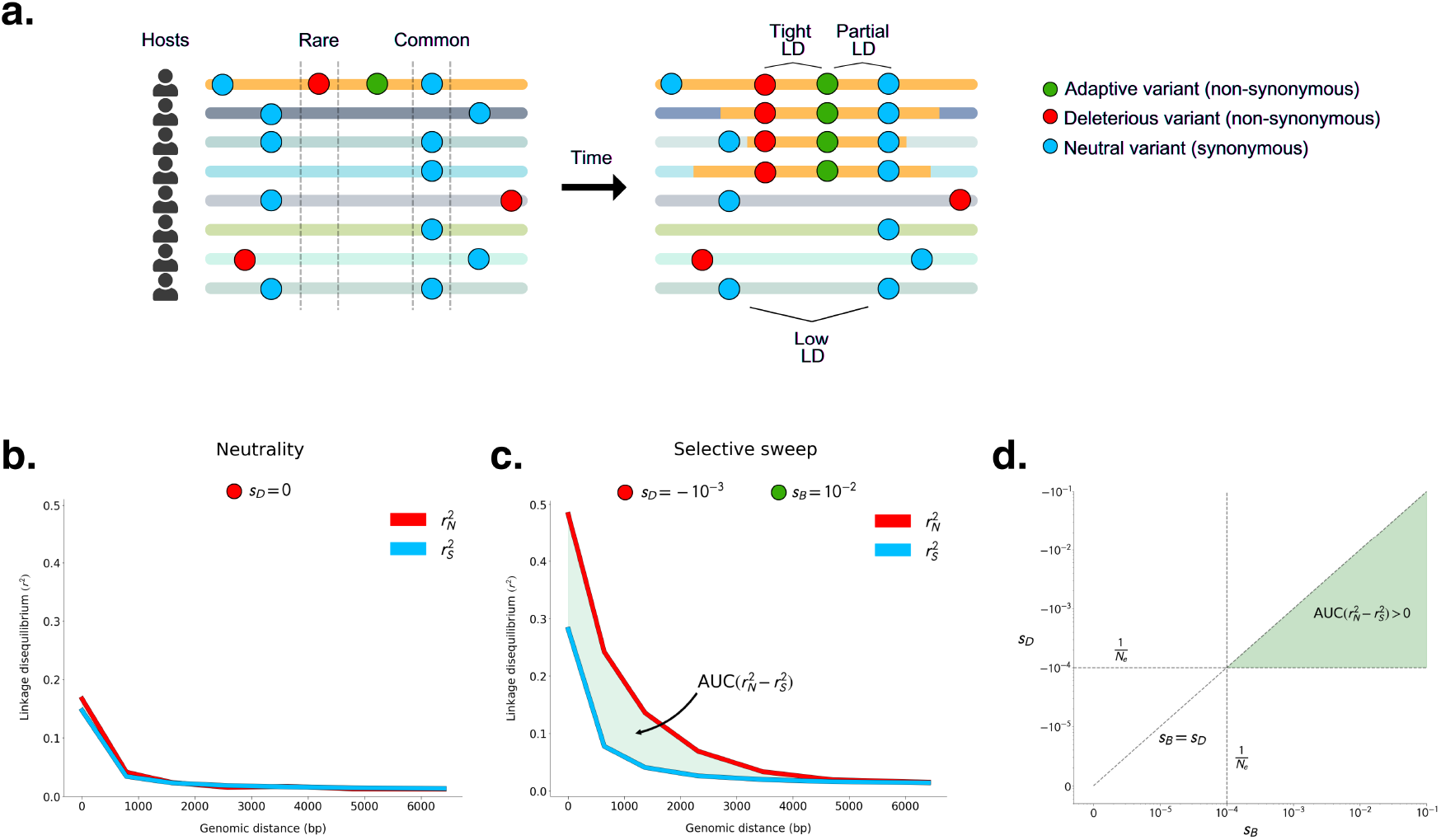
Linkage disequilibrium among common non-synonymous versus synonymous variants during a selective sweep. **(A)** Genomic fragment bearing adaptive variant sweeping across host microbiomes. Each horizontal line represents a bacterial haplotype from a different host’s microbiome. The yellow region of each haplotype represents a fragment that bears an adaptive allele that has recombined onto different lineages’ backgrounds. **(B)** 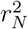 and 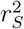 among common variants under neutrality. **(C)** AUC 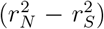 among common variants where *s*_*D*_ = *−*10^*−*3^ and *s*_*B*_ = *−*10^*−*2^. **(D)** AUC 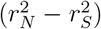 is expected to be greater than zero when *s*_*B*_ *> s*_*D*_ and both *s*_*D*_ and *s*_*B*_ are stronger than the effects of drift (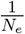, dashed lines). In this schematic and in all simulations (prior to a demographic contraction), *N*_*e*_ = 10^4^. See **Figures S1 S3** for 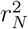 and 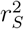 measured across a comprehensive set of simulated evolutionary scenarios.

In this work, we first confirm our hypothesis that deleterious hitchhiking drives an increase in LD among non-synonymous relative to synonymous variants in simulations. We further find that this signal does not manifest under neutrality, as a result of purifying selection alone, or due to low recombination rates or demographic contractions. Next, in a panel of 32 prevalent and abundant gut microbiome species, we find that elevations of LD among non-synonymous variants are common at the whole genome level, suggesting that positive selection is widespread. Lastly, we develop a novel statistic leveraging these insights (iLDS, the integrated Linkage Disequilibrium Score) to detect specific loci under selection in these gut microbial species. Application of iLDS to human metagenomic data from 24 populations around the world reveals more than 300 instances of adaptations that have spread across hosts, as well as differences in the targets of selection between Westernized and non-Westernized microbiomes.

## Results

### Positive selection generates elevated linkage disequilibrium among common non-synonymous variants compared to synonymous variants

We first test whether positive selection can drive an excess of LD between pairs of non-synonymous variants 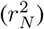 versus pairs of synonymous variants 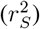 when deleterious variants hitchhike with a positively selected variant. To do so, we performed forward population genetic simulations of selective sweeps in SLiM 4.0 [32] (Supplementary Section 2). While the beneficial variant and any hitchhikers may be expected to become common in the population, deleterious variants not linked to the adaptive variant should remain rare. Assuming all non-synonymous sites are either subject to purifying selection or are adaptive, we expect non-synonymous variants that become common to either be adaptive or to have hitchhiked with and therefore be tightly linked to an adaptive variant. As a result, we expect that 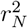 will be elevated relative to 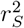 specifically among common variants (**Figure 1A**).

To examine the potential effects of purifying and positive selection on patterns of LD, we analyzed LD among variants that are either rare (minor allele frequency MAF *≤* 0.05) or common (MAF *≥* 0.2) in the broader population, respectively. To quantify whether 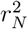 is significantly elevated over 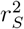, we computed the difference in area under their respective LD distance decay curves (AUC) (**Figure 1C**). This test statistic, which we refer to as AUC 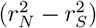, allows us to assess differences in total levels of 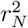 and 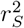 in a manner that controls for genomic distance (and therefore effective recombination rates) between pairs of alleles (Supplementary Section 1.2).

Before assessing if selective sweeps generate excess LD among common non-synonymous versus synonymous variants, we first determined if this pattern can arise under scenarios of neutrality, purifying selection, or demographic contractions. As expected, under neutrality, we observed that AUC 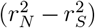 was not significantly different from zero for either common or rare variants (**Figure 1B** and **S1 S6**). Similarly, we found that in populations evolving under purifying selection, in which new non-synonymous mutations experienced purifying selection of strength (*s*_*D*_) varying from *−*10^*−*5^ to *−*10^*−*1^ (encompassing a value weaker than the effect of drift (|*N*_*e*_*s*_*D*_|*<* 1) to very strong selection (|*N*_*e*_*s*_*D*_|*≫* 1)), common variants failed to produce AUC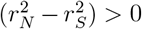, irrespective of the recombination rate. However, in these scenarios of purifying selection rare variants showed a depression in 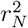 versus 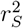 (**Figures S4** - **S6**), consistent with both Hill-Robertson interference [33] or epistasis between deleterious variants, as previously observed by [34, 35, 36, 37, 38]. Finally, given that demographic contractions are known to affect patterns of diversity and linkage disequilibrium in ways that closely resemble sweeps [25, 26, 27], we tested if a population bottleneck could lead to a stochastic increase in the frequency of haplotypes bearing particular combinations of linked deleterious variants, and therefore potentially to an elevation of 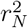 versus 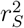 among common variants. However, in two demographic scenarios tested, AUC 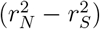 was not significantly different from zero (**Figures S1** - **S3**), irrespective of the recombination rate.

Next, we tested whether selective sweeps could induce AUC 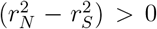 among common variants. To do so, we introduced a novel, beneficial mutation to a population already evolving under purifying selection, and allowed it to rise to intermediate (50%) frequency. The strength of beneficial selection (*s*_*B*_) ranged from nearly-neutral (10^*−*5^) to strongly beneficial (10^*−*1^). First, regardless of *s*_*D*_, 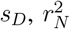 and 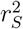 among common variants generally increased monotonically with *s*_*B*_, reflecting the decrease in the expected time for the sweeping variant to reach intermediate frequency relative to neutrality. Second, we found that selective sweeps can in fact produce AUC 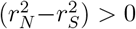; however, this pattern only manifests under particular combinations of *s*_*B*_ and *s*_*D*_. Specifically, the strength of purifying selection must exceed drift (i.e. *s*_*D*_ *>* 1*/N*_*e*_), and the strength of positive selection must exceed that of purifying selection (*s*_*B*_ *> s*_*D*_) (**Figure 1D**). Additionally, AUC 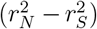 increased with the strength of *s*_*B*_ and *s*_*D*_, as well as with the rate of recombination (**Figures S1** - **S3**). Moreover, 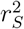 remained elevated over 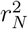 among rare variants during the selective sweep, provided purifying selection exceeded drift (**Figures S4** - **S6**). Thus, when a population experiences both purifying and positive selection, we expect to see differences between synonymous and nonsynonymous LD among both rare and common variants.

### Elevation of LD among non-synonymous variants in gut commensal species

Having established in simulations that LD between common non-synonymous variants can be elevated relative to synonymous variants primarily due to selective sweeps, we next quantified 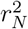 and 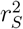 across human gut microbiomes to assess if this signature of positive selection is observed at a genome-wide scale in gut microbiome species. To do so, we analyzed data from metagenomic samples of 693 individuals from North America, Europe, and China [39, 40, 41, 42]. To identify single nucleotide polymorphisms (SNPs) from these samples, we aligned shotgun reads to a database of reference genomes using MIDAS [43] (Supplementary Section 3). We showed previously that samples in which a single dominant strain of a species is present can be confidently ‘quasi-phased’ such that pairs of alleles can be assigned to the same haplotype with low probability of error, and that subsequently LD can be computed between these pairs of alleles [1]. With this quasi-phasing approach, we extracted 3316 haplotypes belonging to 32 species across the 693 individuals we examined. Some of the species examined exhibit considerable population structure, with strong gene flow boundaries between clades, so we focused our analyses only on haplotypes belonging to the largest clade of each species (Supplementary Section 3.6) [1, 12].

First, we examined the dependence of AUC 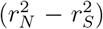 on allele frequency. As purifying selection drives deleterious variants to low frequencies and positive selection tends to elevate allele frequencies, we expect to observe a generally positive relationship between allele frequency (*f*) and AUC 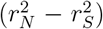 if both purifying and positive selection affect these populations. In **Figure S17**, we see that AUC 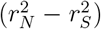 universally increases with allele frequency, as expected. Additionally,we see that AUC 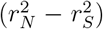 flips from negative to positive when *f ≥* 0.05 in most species. It is possible that the majority of non-synonymous variants with allele frequencies below this threshold are deleterious, while those with allele frequencies above this threshold are more likely to be either beneficial themselves or tightly linked to a beneficial variant.

Shown in **Figure 2A** are examples of genome-wide 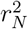 and 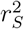 for the species *Ruminococcus bromii* and *Prevotella copri*. Among both rare (*f ≤* 0.05) and common (*f ≥* 0.2) variants, 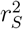 and 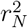 decay with increasing distance between pairs of genomic loci, as expected for recombining species. The rate of decay differs among species; however, for all species, LD appears to eventually saturate to some roughly constant value. In *R. bromii*, for instance, both rare and common variant LD appear to saturate around *∼*10Kb. In Supplementary Section 5.1, we show how the initial decay and eventual saturation of LD can be related to an underlying model of recombination, which in turn can be used to infer the mean tract length of horizontally transferred segments for each species.

**Figure 2:**
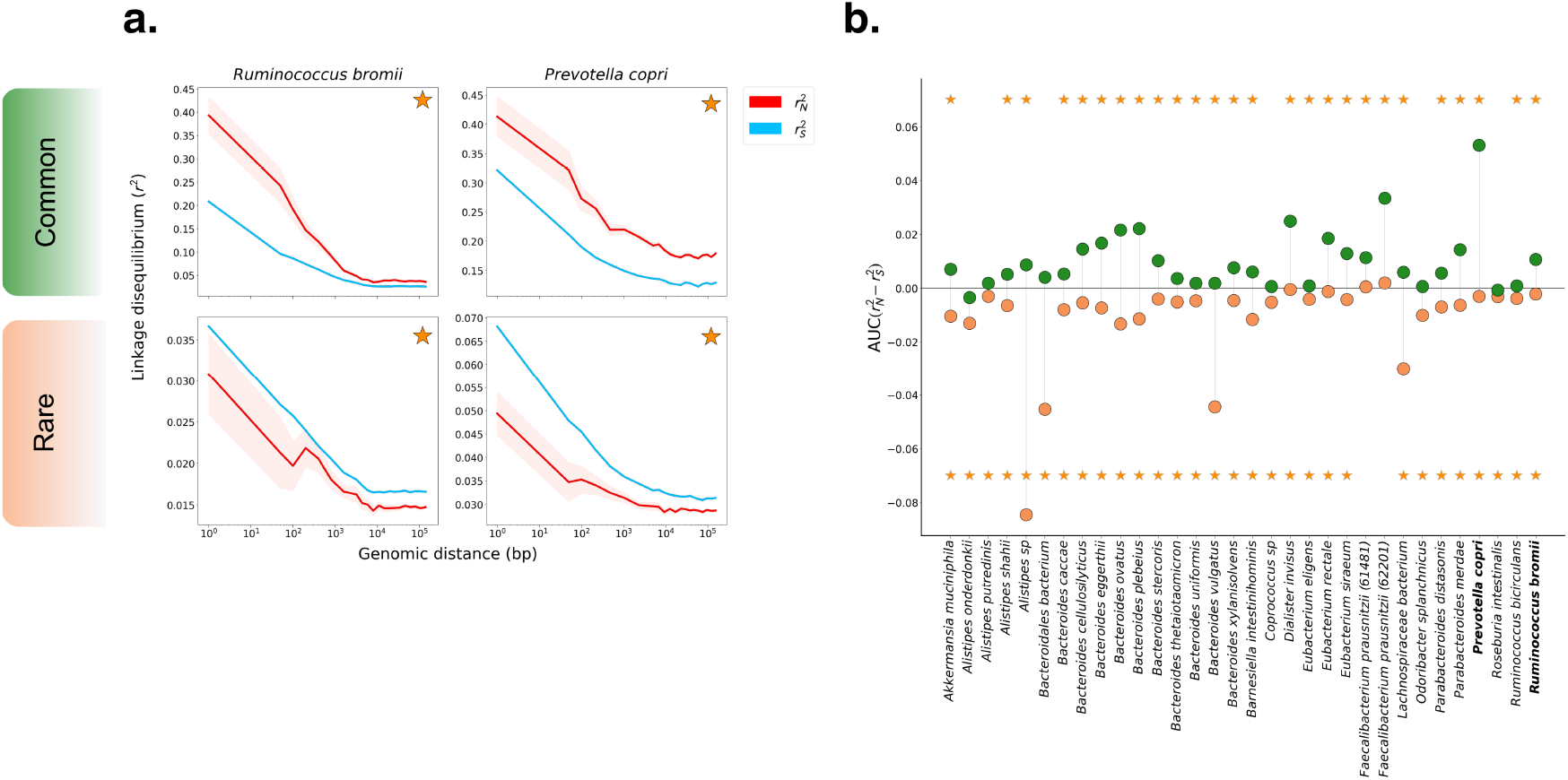
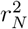 and 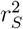 measured in prevalent commensal gut microbiota. **(A)** Decay in LD among common (MAF *≥* 0.2) (top) and rare (MAF *≤* 0.05) (bottom) variants for the species *Ruminococcus bromii* and *Prevotella copri*. Both species show significant differences between 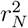 and 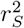 for common and rare variants, as denoted by the orange star. **(B)** AUC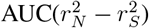 among rare(orange) and common (green) alleles for 32 prevalent gut commensal bacteria species. Among rare variants, AUC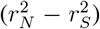 is significantly negative for all but two species (yellow stars, at bottom). Among common variants, AUC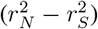 is significantly positive in 26/32 of species (yellow stars, at top).

For both species in **Figure 2A**, AUC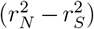 is significantly greater than zero among common variants and less than zero among rare variants. More broadly, across the 32 species analyzed, AUC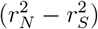 is significantly greater than zero among common variants in 26/32 species. Among rare variants, AUC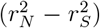 was significantly less than zero for all but two species (**Figure 2B**). Together, these patterns of LD among synonymous and non-synonymous variants are consistent with widespread purifying and positive selection acting on non-synonymous sites in these species.

### Detecting HGT-mediated selective sweeps with iLDS

Genome-wide patterns of LD among synonymous and non-synonymous variants indicate that selection—both positive and purifying—is pervasive at the nucleotide level in gut microbiome species. While only a minority of intermediate frequency non-synonymous sites are likely adaptive, positive selection at these sites is evidently strong enough to create highly significant genomewide linkage patterns. To identify these specific adaptive loci, we developed a novel statistic—the integrated Linkage Disequilibrium Score (iLDS)—which detects genomic regions exhibiting both AUC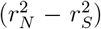*>* 0 and elevated LD relative to the genomic background. By combining these sources of information, we identify regions which have elevated LD due to positive selection and not other non-selective forces.

To detect specific genomic regions under selection, iLDS is calculated in sliding windows across a genome. To calculate iLDS in a genomic window, we first determine AUC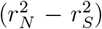 among common SNVs (MAF *≥* 0.2) within the window. Next, to augment our ability to detect selection, we also identify windows with elevated LD overall, which is expected for selective sweeps. To do so, we compute the difference in the area under the LD curve between all intermediate frequency variants in the same window (i.e. AUC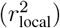), irrespective of whether they are synonymous or non-synonymous, and the area under the average genome-wide LD curve over the same distance defined by the window (AUC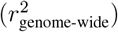). The two components of iLDS are therefore:

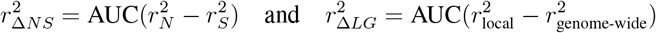

Next, each component is standardized by its mean and standard deviation across all windows along the genome:

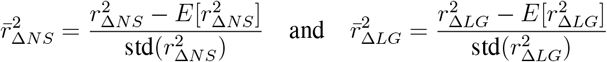

Finally, the statistic is defined as:

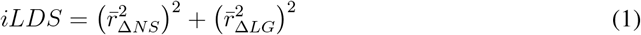

In essence, 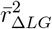 quantifies the increase in total LD within the window relative to the expected level of LD across the whole genomic background for a region of the same size, while 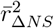 quantifies the local extent of elevation in LD among non-synonymous variants relative to synonymous variants. Both of these terms are expected to be elevated during a sweep; however, iLDS should not be elevated in regions where 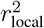 is high due to non-selective factors, as AUC 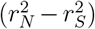 will remain near zero in such regions.

In order for a window to be called as significant, three criteria must be met. First, iLDS must exceed some critical value. In simulations, we found that 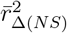 and 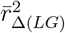 each had approximately standard normal distributions in the absence of positive selection (**Figure S19**). Therefore, iLDS should approximately follow a *χ*^2^ distribution with two degrees of freedom. Sweeps, by contrast, produce iLDS values falling in the upper tail of the *χ*^2^ distribution (**Figure S18**). Thus, we set the critical value of iLDS to be the upper alpha percentile point of the *χ*^2^ distribution (in this work, we employ an *α* = 0.05). If iLDS exceeds this critical threshold, we additionally require that AUC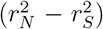*>* 0 and AUC 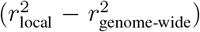 *>* 0. Together, these criteria ensure that LD patterns within windows called as significant are consistent with selection.

We tested iLDS’s ability to correctly detect selective sweeps, as well as its potential for misclassifying genomic regions with elevated LD arising from demographic contractions as selective sweeps (Supplementary Section 5.3). We found that iLDS is powerful in detecting recent and strong selective sweeps (**Figures S7** - **S9**). Further, we found that demographic contractions do not substantially elevate the false positive rate or false discovery rate of iLDS. Finally, we find that in most scenarios where iLDS has strong ability to detect selection, the false discovery rate rarely surpasses 10% (**Figures S10** - **S12**). These simulation results indicate that overall, iLDS is capable of correctly identifying sweeps when *s*_*B*_ is sufficiently strong and rarely identifies non-sweeps as sweeps.

### iLDS reveals pervasive selective sweeps in gut bacteria

We next applied iLDS to gut bacteria. To do so, it is first necessary to define genomic windows to calculate iLDS in. The window size should ideally be large enough that genome-wide LD can be expected to fully decay by the edges of the window, but not so large that the footprint of the sweep is very small relative to the size of the window. To determine this species-specific window size in the bacteria examined here, we estimated a typical upper bound on the size of a horizontally transferred tract *l*_*DD*_ under an idealized model of HGT (Supplementary Section 5.1). LD should fully decay at approximately *l*_*DD*_ as linkage between fragments separated by greater than this distance is always broken by recombination, while variants which are closer may be transferred together horizontally. By visual inspection, we found that the inferred value of *l*_*DD*_ did in fact correspond to the point at which LD fully decayed among common synonymous variants in the data (Supplementary Section 5.1, **Figure S26**, Table S3). To ensure that each window contains both an adequate and comparable number of synonymous and non-synonymous variants with which to calculate 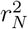 and 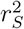 curves, we employed a SNP based windowing approach as opposed to a base-pair defined window. Specifically, we defined each window to consist of the average number (for that species) of consecutive non-synonymous, intermediate frequency SNPs (MAF *≥* 0.2) spanning *l*_*DD*_ (Table S3).

To assess the ability of iLDS to detect known instances of positive selection in a natural population, we applied iLDS to a set of 132 isolates of *Clostridiodes difficile* (**Figure 3A**, Methods), an enteric pathogen that has experienced a recombination mediated selective sweep at the *tcdB* locus, which encodes the toxin B virulence factor [45, 46]. In the majority of windows, iLDS remains close to zero, as expected in the absence of positive selection. However, the value of the statistic peaks sharply in several regions across the genome. Many of these peaked regions contain large numbers of significant iLDS values in consecutive windows. Since consecutive windows may belong to the same selective event, we clustered groups of significant windows into a peak if the SNPs they were centered around were both physically close and tightly linked to one another, as would be expected following a selective sweep (Methods). In total, we identified seven putative selective sweeps in *C. difficile*. One of these peaks overlaps *tcdB*, confirming that iLDS can indeed recover known instances of positive selection (**Figure 3A**).

**Figure 3:**
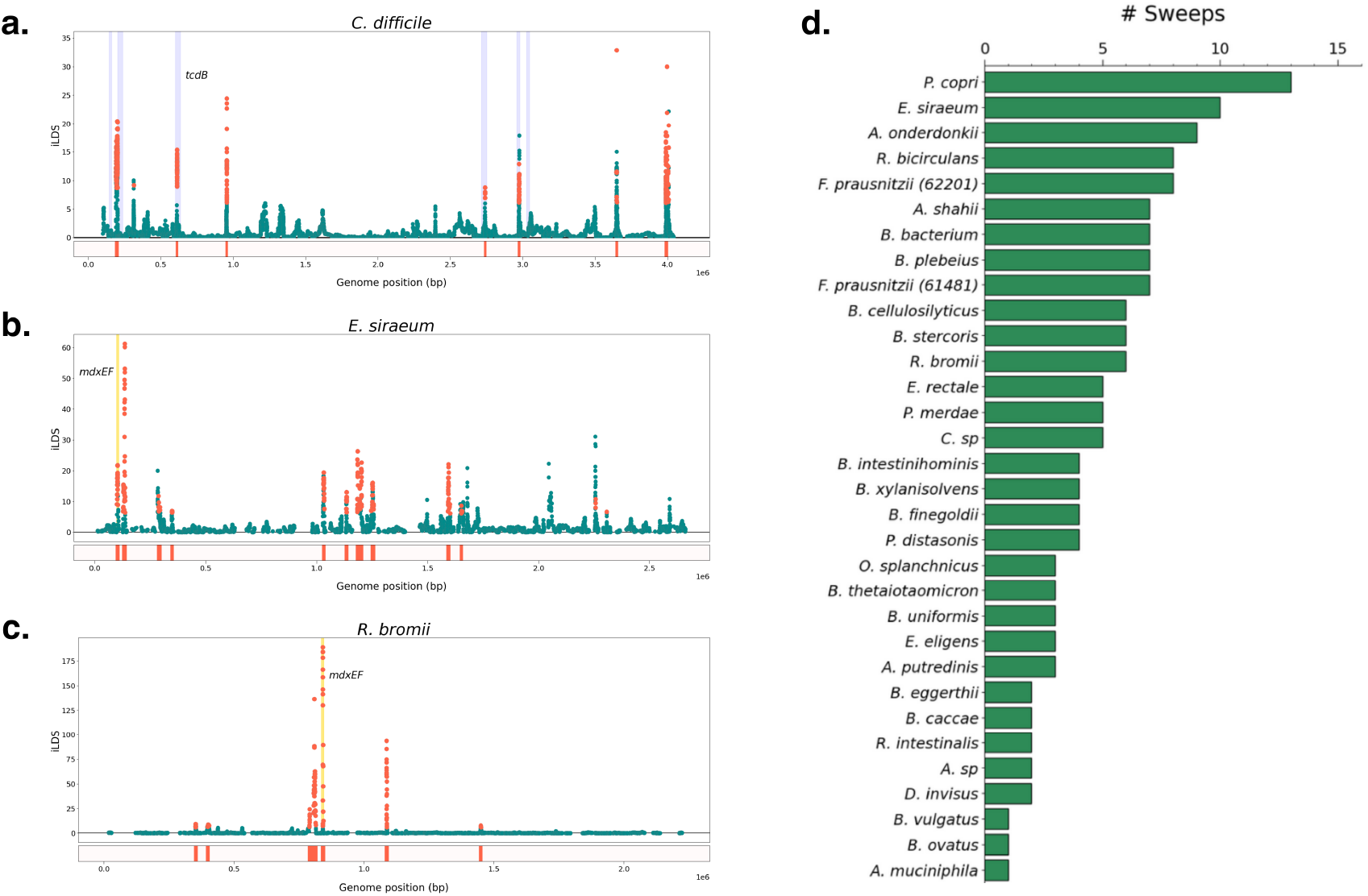
Recombinant selective sweeps in gut bacteria. **(A)** iLDS scan in *C. difficile*. Each point corresponds to an iLDS value for a given genomic window centered around a single intermediate frequency non-synonymous SNP. Significant windows are colored orange, while non-significant windows are colored green. The locations of peaks are shown as orange bars below the scan. Highlighted in blue are the locations of the genes predicted to be virulence factors (Methods [44]). iLDS scans for **(B)** *E. siraeum* and **(C)** *R. bromii*, respectively. Both species exhibit a peak at the genes *mdxEF*, highlighted in yellow, as well as nine other loci in *E. siraeum* and five other loci in *R. bromii*. **(D)** Number of selective sweeps detected per species.

Beyond *tcdB*, we find a striking correspondence between the locations of the putative sweeps and known virulence factors in the *C. difficile* genome (**Figure 3A**, blue bars—see Supplementary Section 4.1). For instance, one peak overlaps the *fli* operon, a virulence factor involved in flagellar biosynthesis, which has been previously hypothesized to be under positive selection [47]. In total, out of the seven regions annotated to have virulence factors, four overlap or are near iLDS peaks, potentially indicating that selection on virulence-associated traits is an important component of *C. difficile* evolution.

Finally, once again confirming the ability of iLDS to recover known positive controls, in the recombinant pathogen *Helicobacter pylori*, iLDS generate a significant peak at the *vacA* virulence factor gene (encoding the vacuolating cytotoxin) (**Figure S23**), which was previously shown to experience positive selection [48, 49]. Overall, we find evidence that virulence factors may be positively selected in these pathogens.

Next, we applied the scan to the 32 gut microbiome species analyzed in **Figure 2B**. We identified a total of 155 unique peaks across all species, with a median of four peaks per species (**Figure 3D**). In total, these peaks spanned 452 genes (Supplementary Table S4). While these genes were functionally diverse, we found certain classes of genes repeatedly under selection. For example, we identified five instances in five unique species of peaks spanning *susC*/*susD* starch utilization system genes, which have previously been found to be under selection within multiple, independent hosts over weeks to months [2, 7]. Among all 452 genes overlapping iLDS peaks, we observed an enrichment for carbohydrate transport and metabolism genes overall (COG category G [50]; adjusted p-value *<* 5 *×* 10^*−*7^; Methods) and specific classes of enzymes involved in carbohydrate metabolism—particularly glycoside hydrolases (EC number 3.2.1 [51]; adjusted p-value = 0.02). These enrichment results provide evidence that genes related to the breakdown and transport of carbohydrates are frequently targeted by selection in the gut microbiome (**Figure S22**).

One particular class of carbohydrate metabolism genes repeatedly detected as under selection were *mdxE* and *mdxF*, ABC transporters capable of metabolizing maltodextrin [52], a starch derivative commonly used as an emulsifier and textural component of ultra-processed foods [53, 54]. The genes mdxEF are present in only four unique species in our dataset, and are identified as under selection by iLDS in two of these species: *R. bromii* and *E. siraeum* (**Figure 3B** and **3C**), both known to metabolize starches in the colon [55, 56]. Indeed, *mdxEF* are overrepresented among targets of selection relative to their genomic frequency (Benjamini-Hochberg adjusted p-value = 0.054). By inspecting the haplotypes at and surrounding *mdxEF*, we see that this putatively adaptive region exhibits evidence of extensive, recent horizontal gene transfer (**Figure S25**).

### Selective sweeps across continents and lifestyles

The shift from traditional to Westernized lifestyle has reshaped the gut microbiome, altering its ecological composition and causing an overall reduction in diversity [57]. Previous work has demonstrated that the shift to Westernization has also altered evolutionary trajectories within species, with Westernization driving elevated rates of HGT [14] as well as differentiation in the pool of mobile genetic elements [58]. We hypothesized that the targets of adaptation in Westernized and non-Westernized microbiota are distinct, as a consequence of selective pressures specific to each group. To test this hypothesis, we performed iLDS scans in metagenome assembled genomes from the Unified Human Gastrointestinal Genome catalog [59] for 16 species present in healthy hosts from both Westernized populations in Europe, Asia, and North America, as well as non-Westernized populations from Fiji [58] Tanzania [60, 61], Madagascar [62], Peru [63], El Salvador [64], and Mongolia [65] (**Figure 4A**). In total, we analyzed 24 populations around the world (19 Westernized, five non-Westernized).

**Figure 4:**
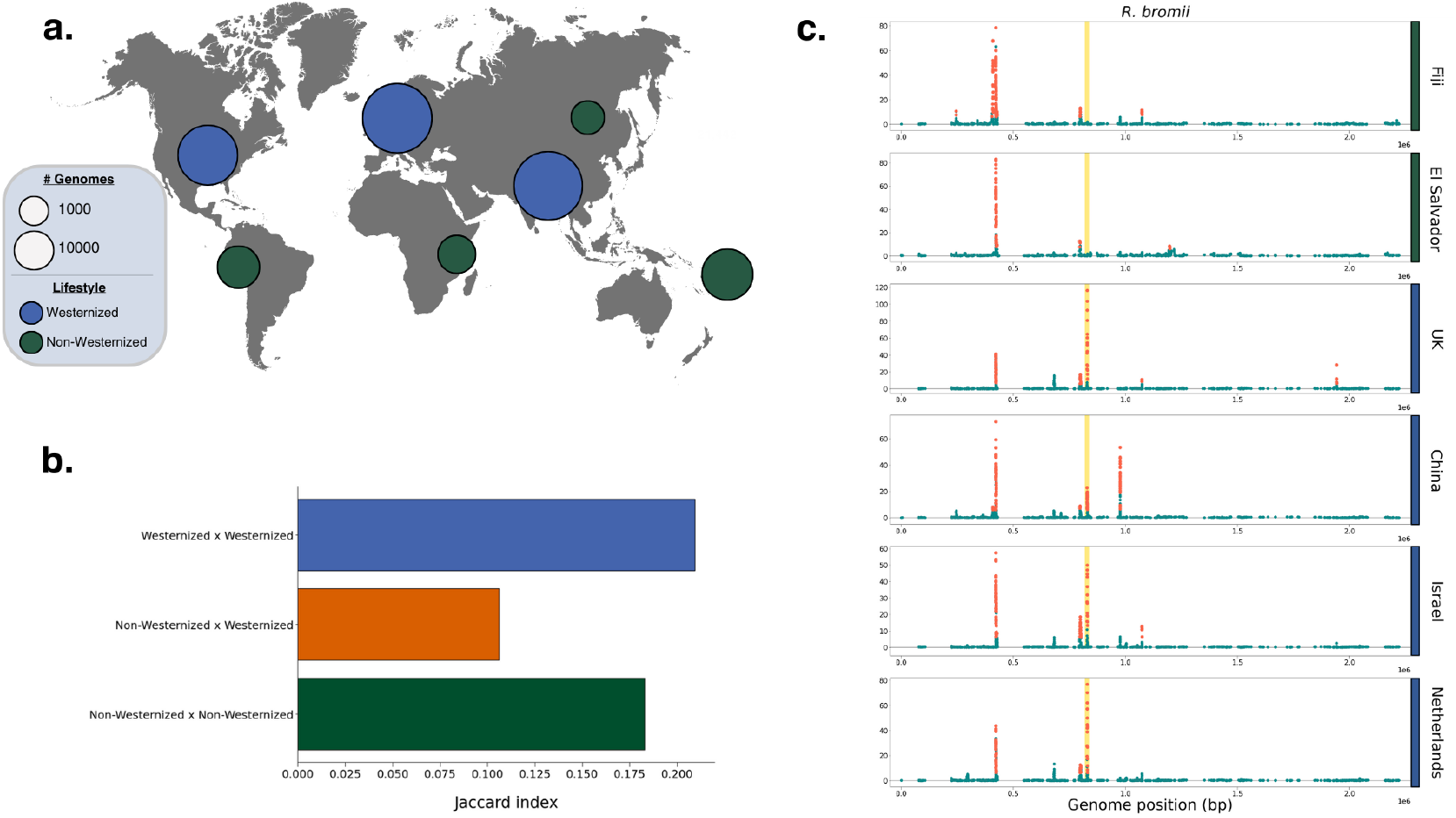
Selective sweeps across continents and lifestyles. **(A)** Numbers of bacterial genomes analyzed per continent for Westernized vs non-Westernized populations, as indicated by circle size. **(B)** Overlap in the locations of peaks between Westernized and non-Westernized populations, as determined by the Jaccard index (Methods) **(C)** Selective sweeps in *R. bromii* in two non-Westernized populations and four Westernized populations from around the world. The *mdxEF* genes are highlighted in gold. For the full set of scans across all 16 populations analyzed, see **Figure S31**.

We found first that many selective sweeps have spread between countries and across continents. Across the 24 populations and 16 species studied, we detected a total of 309 unique selective sweeps. While the majority of sweeps were unique to a single population, 35% were shared across populations, with 108 sweeps found in more than one country, and 83 found on multiple continents (**Figure S29**). Some sweeps were extremely broadly distributed, with 26 sweeps present in 50% or more of populations and eight present in 80% or more.

Next, we assessed whether sweeps were more commonly shared between Westernized populations. To do so, we calculated a Jaccard index (*J*) to quantify the proportion of sweeps shared between populations (Methods). Consistent with our hypothesis, we found that Westernized populations share sweeps with one another at more than double the frequency (*J* = 0.21, p-value = 0.047, permutation test, Methods) than they do non-Westernized populations (*J* = 0.11, p-value *<* 10^*−*4^) (**Figure 4B, Figure S30**). Similarly, non-Westernized populations also shared sweeps with one another (J = 0.18, p-value = 0.67) at greater frequency than with Westernized populations, though this elevated sharing was not statistically significant. Together, these results indicate that there are shared selection pressures experienced across Westernized populations that drive evolutionary differentiation from non-Westernized populations.

Beyond the evident aggregate difference in sweeps between Westernized and non-Westernized populations, we also identified specific selective sweeps which were unique to one group or the other. In total, we identified 32 sweeps present in 50% or more of populations of one type (i.e. Westernized or non-Westernized) but absent in populations of the other type. Of these, 24 were unique to Westernized populations and eight to non-Westernized populations. By contrast, only three sweeps were found to be present in *≥* 50% of both Westernized and non-Westernized populations, underscoring the lack of shared selective pressures between these populations.

The *R. bromii mdxEF* locus, discussed in the preceding section (**Figure 3C**), was among the sweeps which exhibited the strongest pattern of differential selection between Westernized and non-Westernized populations. Indeed, *mdxEF* was found to be under selection in all fourteen Westernized populations but neither non-Westernized population in which the species was present (Fiji and El Salvador) (**Figure 4C**), suggesting that this species may be adapting specifically to Westernized lifestyles. Furthermore, this locus had the largest value of iLDS for this species in six out of 14 Westernized populations (in addition to being the highest peak in **Figure 3C**), indicating that these genes may be under particularly strong selection.

While some targets of selection may differ between Westernized and non-Westernized microbiota, we also found that the total number of selective sweeps per population were similar, indicating the gut microbiota of non-Westernized populations may be adapting at a similar rate to those of Westernized populations. In particular, we found Westernized populations tended to harbor slightly more sweeps (3.46 sweeps/population) than non-Westernized populations (3.25 sweeps/population); however, this difference was not statistically significant (permutation test, p-value = 0.6). We note that the non-Westernized populations studied here are heterogeneous in lifestyle—including pastoralist (Mongolia) and agrarian (Fiji) populations—and may also differ in both the extent of their contact with Westernized populations as well as their adoption of Westernized dietary and lifestyle practices.

## Discussion

In this paper we perform the first comprehensive scan for adaptive fragments that have swept across human gut microbiomes via horizontal gene transfer. To do so, we develop a novel selection scan statistic, iLDS, that is sensitive to elevations in LD between pairs of common nonsynonymous SNPs vs pairs of common synonymous SNPs. We show in simulations that this signature is expected when deleterious variants hitchhike to high frequency along with beneficial variants. Application of iLDS to metagenomic data from 24 populations around the world revealed more than 300 candidate selective sweeps across more than 30 bacterial species. Among these sweeps, we find a strong enrichment of loci involved in carbohydrate transport and metabolism, suggesting that host diet may play an outsize role in driving adaptive evolution in these species. Additionally, present in all Westernized populations and absent from all non-Westernized populations analyzed is a selective sweep at a locus potentially involved in the metabolism of a synthetic starch added to highly processed foods. Taken together, our results indicate that recombination between strains fuels pervasive adaptive evolution among human gut commensal bacteria, and strongly implicate host diet as a critical selection pressure for these species.

Our work adds to a growing literature suggesting that host diet not only changes the species composition of the microbiome, but also selects for specific genetic variants within species. Indeed, human populations consuming diets that are rich in seaweed glycans [8], red meat [66], and plant starches [58] appear to select for genes in particular bacterial species which facilitate the metabolism of these substrates within the host. Additionally, mouse experiments [67, 68] have shown that adaptations arise within hosts on short time scales of weeks to months in response to Western-style high fat and high sugar diets. We build on these findings by demonstrating that adaptations to host diet are broadly distributed across many pathways in many different species.

We also uncover striking instances of adaptation at specific loci. For instance, we found a single selective sweep at the *mdxEF* locus in *R. bromii* which was ubiquitous in Westernized populations but absent from non-Westernized populations. While the precise selection pressures driving the spread of *mdxEF* variants across Western populations are unclear, these genes are known to facilitate growth on maltodextrin—a synthetic starch derivative commonly used as an emulsifier and textural component of ultra-processed food [53, 54]—raising the possibility that this selective sweep represents an adaptation to a novel source of dietary starch in the Western diet. As *R. bromii* occupies a unique ecological niche within the gut microbiome [55], where it is a keystone species for the metabolism of resistant starches (i.e. starches which escape digestion by the host), adaptations in this species may have outsize effects on the ecological structure of the microbiome and the production of resistant starch fermentation byproducts, such as short-chain fatty acids. Future work investigating the functional and ecological consequences of the selective sweeps we have identified, likely via experimental studies, will be important for understanding the role of each genetic variant in the microbiome.

Previous attempts to scan for signatures of positive selection across the genome in gut microbiome species have tended to employ single locus approaches—looking for signatures such as parallelism or elevated dN/dS ratios [1, 2]. Such approaches are ideally suited to detect selective sweeps within hosts from *de novo* mutations, for instance, but are underpowered to detect gene-specific sweeps as they do not leverage LD between sites. Gene-specific sweeps in natural populations of bacteria have been instead discovered via searching for dramatic reductions in local diversity coupled with preservation of SNP densities in flanking regions [15, 24]. However, thus far, examples of gene specific sweeps in the literature have largely been discovered by case-by-case studies as opposed to systematic application of a haplotype-based selection scan statistic, such as iHS [20]. The precise reasons for why such statistics have been rarely applied are unclear, though we do find that iHS exhibits limited ability to detect known HGT-mediated sweeps in *C. difficile* (**Figure S24**). By contrast, iLDS is highly successful in detecting positive controls in multiple species of bacteria (**Figure 3**A and **Figure S23**). Moreover, we find that iLDS is versatile enough to be applied to any recombining species, and as such we demonstrate it is capable of detect positive controls in *Drosophila melanogaster* (**Figure S32**, Supplementary Section 7). iLDS may have power across a range of species because it exploits a common signature associated with selective sweeps: deleterious variants hitchhiking to high frequency with a beneficial driver. To our knowledge, the tight linkage of beneficial variants with hitchhiking deleterious variants, which has been shown to be a common feature of adaptation both in theory and in numerous systems [69, 70, 71, 72, 73, 74, 75, 76, 77], has not been explicitly incorporated into any selection scan statistic.

We note that others have also observed that elevated LD among non-synonymous variants relative to synonymous variants can be a signature of adaptation [78, 79, 80, 81]; however, the connection with deleterious hitchhiking had not previously been noted. Stolyarova *et al*. [78] and Callahan *et al*. [81] found that epistatic interactions could generate elevated LD among non-synonymous variants in the highly polymorphic fungus *Schizophyllum commune* and also in *Drosophila* species, respectively, while Arnold *et al*. [79] concluded that epistasis was not necessary to generate this signal in *Neisseria gonorrhoeae*, and that adaptive inter-specific HGT of short genomic fragments bearing multiple positively selected non-synonymous alleles was the likely driving factor. We emphasize that our findings are fully consistent with those of Stolyarova *et al*., Callahan *et al*., and Arnold *et al*. But crucially, our results suggest that elevated LD among common non-synonymous variants is not by itself sufficient to establish that all such variants are adaptive or epistatically interacting. Because purifying selection at the vast majority of non-synonymous sites is well-established to be a pervasive feature not only of bacterial genomes [1, 82, 83, 84, 85], but also in the genomes of most other species [86, 87, 88]. We believe it is highly likely some proportion of common non-synonymous polymorphisms will be deleterious hitchhikers in any adapting population, with this proportion growing, paradoxically, as the strength of positive selection increases. In future work, disentangling the effect of epistasis versus hitchhiking on deleterious alleles will be important for understanding the relative contributions of different population genetic forces driving selective sweeps in bacteria and other natural populations.

Most of the selective sweeps we identify are likely real and are not false positives given low false discovery rates frequently on the order of 10% for recent and strong selective sweeps (**Figures S10** - **S12**). However, the FDR is likely much lower than what we have measured from simulations, which treats each analysis window as independent from one another. In data, we require that multiple adjacent windows are significant, and, all windows supporting a putative sweep have central SNPs that are tightly linked. Additionally, the low TPR measured from simulations even for the strongest and most recent sweeps (60%), likely due to our stringent criteria for identifying a sweep, suggests there are actually many more sweeps that iLDS has not yet detected, which may be weaker or older, where iLDS loses power (**Figures S7** - **S9**). This raises the question of how truly pervasive selection is in gut commensal bacteria. Future work expanding the sensitivities of selection scan statistics will be crucial for quantifying the frequency and also targets of adaptation among gut commensal bacteria as well as other organisms.

The high rate of recovery of positive controls and moreover the discovery of hundreds of selective sweeps suggest that that recombination is a major mechanism by which adaptive DNA spreads in human gut microbiome populations. While previous work has found extensive transfer of DNA via HGT across species boundaries [13, 14] and also between strains of the same species across hosts [1, 12], we definitively establish here that certain fragments cannot be spreading due to neutral recombination alone, but rather are being repeatedly transferred in a gene-specific selective sweep. We emphasize that the success of iLDS critically depends on the fact that recombination is ubiquitous across these species’ genomes [1, 12], allowing us to distinguish non-selected regions of the genome—where recombination breaks up LD between even nearby variants—from selected regions.

In conclusion, development and application of iLDS to metagenomic data from diverse populations may help us to learn about unique selective pressures especially relevant to certain human conditions. For example, in future work, application of iLDS to diseased vs healthy cohorts may reveal disease-relevant microbiome loci [89]. The stringent criteria for significance as well as our ability to cleanly detect positive controls in multiple organisms ranging from bacteria to eukaryotes suggests that several of the candidates for selection that we have identified are likely real. Thus, future molecular studies investigating the functional importance of selected loci identified by iLDS may provide mechanistic insight into how microbiome genotypes confer phenotypes to hosts, improve our ability to diagnose and treat diseases associated with specific microbiome variants, and potentially allow us to deploy existing natural, adaptive variation in the design of rational probiotics.

## Acknowledgments

This work was funded by NIGMS NIH award R35GM151023, NSF CAREER award (no. 2240098), and a Paul Allen Research Foundation grant to NRG, as well as a UCLA EEB departmental fellowship to RW. The authors would like to thank Dmitri Petrov for helpful conversations early in the project, Kirk Lohmueller for his help, as well as all members of the Garud lab and Emma Derrick for helpful discussions and feedback on the manuscript.

## Code availability

The source code used to process data, perform analyses, and generate figures is available on GitHub: https://github.com/garudlab/iLDS.

