## Supplementary figures and text for "Pervasive selective sweeps across human gut microbiomes"

### Supplementary materials

### Table of Contents

#### List of Supplementary Tables

**Table S1: Metadata for metagenomic samples.** This table contains a list of the 1,013 shotgun metagenomic samples collected from 693 unique, healthy individuals which we used in this study, and were originally analyzed by Garud & Good (2019) [1]. This includes 250 individuals from Lloyd-Price (2017) (accession numbers PRJNA48479 and PRJNA275349) [2], 250 from Xie *et al* (2016) (accession number PRJEB9576) [3], 185 from Qin *et al* (2012) (accession number PRJNA422434) [4], and 8 from Korpela *et al* (2018) (accession number PRJEB24041) [5]. For each sample used, the subject identifier, sample identifier, accession number, country of the study, continent of the study, and visit number are included, as well as the study from which the sample was drawn.

**Table S2: Clade definitions.** This table contains the top-level clades which were manually defined for this dataset by Garud & Good (2019) [1]. Each row details the species and clade label of a single QP sample.

**Table S3: iLDS scan parameters.** This table shows the parameters used to perform the iLDS scans. In particular, for each species, it shows the tract length $l_r$, decay distance $l_{DD}$, window size, number of bins per window, and the total number of intermediate frequency variants non-synonymous variants present genome-wide. For further methodological details on how these quantities were obtained, see Supplementary Section 5.1.

**Table S4: Peaks.** This table gives the locations and annotations of all intermediate frequency non-synonymous variants inferred to be in a peak of iLDS. An intermediate frequency non-synonymous variant is considered to be in a peak if it is the center of a significant window and is both tightly linked and physically proximate to variants lying at the center of other significant windows (for further details on our approach to obtaining peaks, which are each meant to reflect a single selective sweep, see Supplementary Section 5.4). The columns of the table are:

- **Species**: Name of the species
- **Peak number** the unique peak to which the variant belongs (peaks are numbered from left to right for each species)

- **Contig**: the contig the variant lies on (in the case of peaks for *Drosophila melanogaster*, this is the chromosome)
- **Gene**: the gene the variant lies in
- **SNV position (contig)**: the position of the variant along the given contig
- **SNV position (genome)**: the inferred position of the variant along the entire genome (used only for plotting purposes)
- **Gene product**: the gene protein product
- **GO annotation**: the gene ontology annotation
- **Pathway**: the inferred pathway type the gene lies in
- **Data type**: the dataset the scan was performed on (QP metagenomes, UHGG, or Drosophila)
- **iLDS**: value of iLDS for the variant

**Table S5: Metadata for UHGG samples.** Metadata for strains coming from UHGG used in this study. These sequences, and the metadata presented in this table, were downloaded from UHGG [6]. The SNV catalogs used here are available from the MGnify FTP site: `http://ftp.ebi.ac.uk/pub/databases/metagenomics/mgnify_genomes/`.

**6 Gene enrichment analysis** **61**

**7 Analyses with *Drosophila melanogaster*** **62**

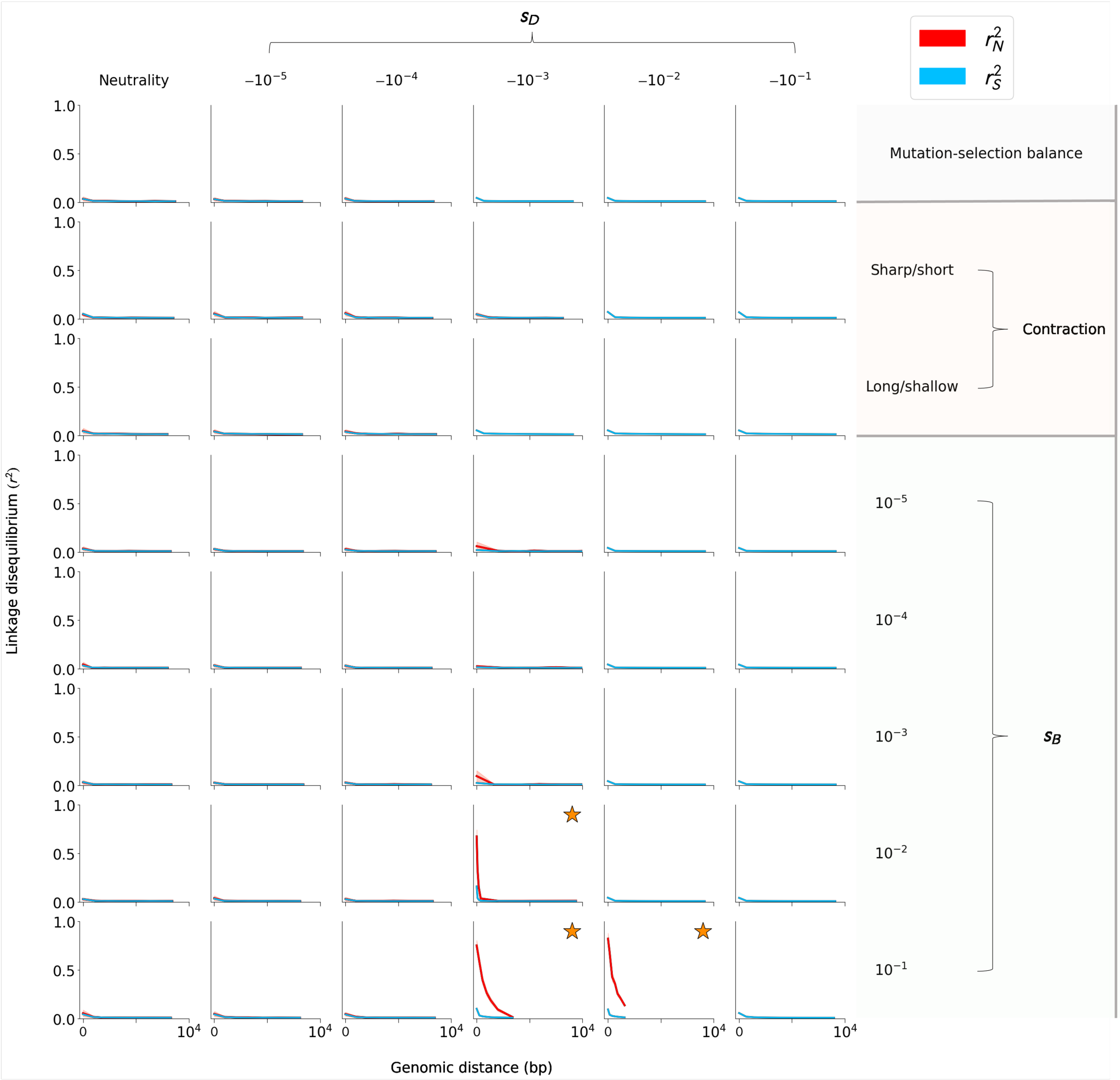

**Figure S1: $r_N^2$ and $r_S^2$ measured among common variants in a range of simulated evolutionary scenarios, where $\rho = 10^{-5}$.** Populations of size $N_e = 10^4$ were simulated with a range of deleterious selection coefficients ($s_D$), and with $\rho/\mu = 0.1$ (see Figures **S2** and **S3** for $\rho/\mu = 1$ and $\rho/\mu = 10$). Top row: $r_N^2$ and $r_S^2$ measured in populations at mutation-selection balance. When $s_D \geq 10^{-2}$, no intermediate frequency non-synonymous variants were observed, and therefore no $r_N^2$ curves are plotted in these columns. Second and third rows: $r_N^2$ and $r_S^2$ measured in populations experiencing a sharp, short population size contraction (second row) and a long, shallow contraction (third row). Fourth row onwards: $r_N^2$ and $r_S^2$ measured in populations experiencing a selective sweep. A total of 250 replicate burn-in populations were simulated until mutation-selection balance had been reached ($10N_e$ generations), followed by five replicates for each burn-in population for the bottleneck and selective sweep scenarios (Section 2). Orange stars denote significantly elevated $r_N^2$ compared to $r_S^2$. Under the strongest selective conditions ($s_B = 10^{-1}, s_D = \{-10^{-3}, -10^{-2}\}$), nearly all common deleterious variants were physically close to the adaptive variant, resulting in apparently truncated LD curves. 99% confidence intervals are plotted as a shaded region around each non-synonymous LD curve, though this region is narrow around some curves.

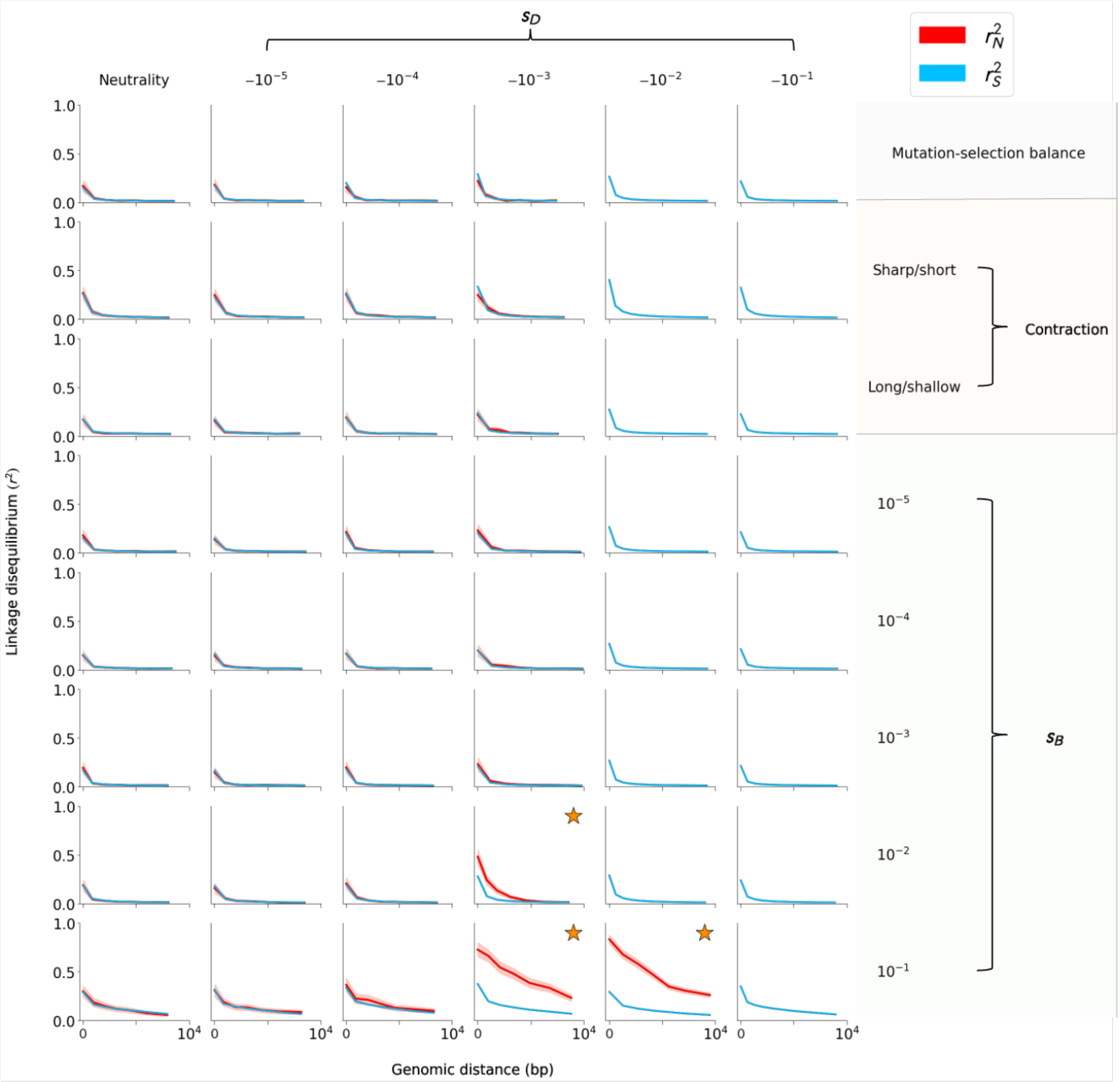

**Figure S2: $r_N^2$ and $r_S^2$ measured among common variants in a range of simulated evolutionary scenarios, where $\rho = 10^{-6}$.** Analogous to Figure **S1**, but for simulations performed with $\rho = 10^{-6}$ (i.e. $\rho/\mu = 1$). As expected, LD decays more rapidly in this regime than with $\rho/\mu = 10$. As with $\rho/\mu = 10$, AUC($r_N^2 - r_S^2$) was significantly greater than zero when $N_e s_D > 1$ and $s_B > s_D$.

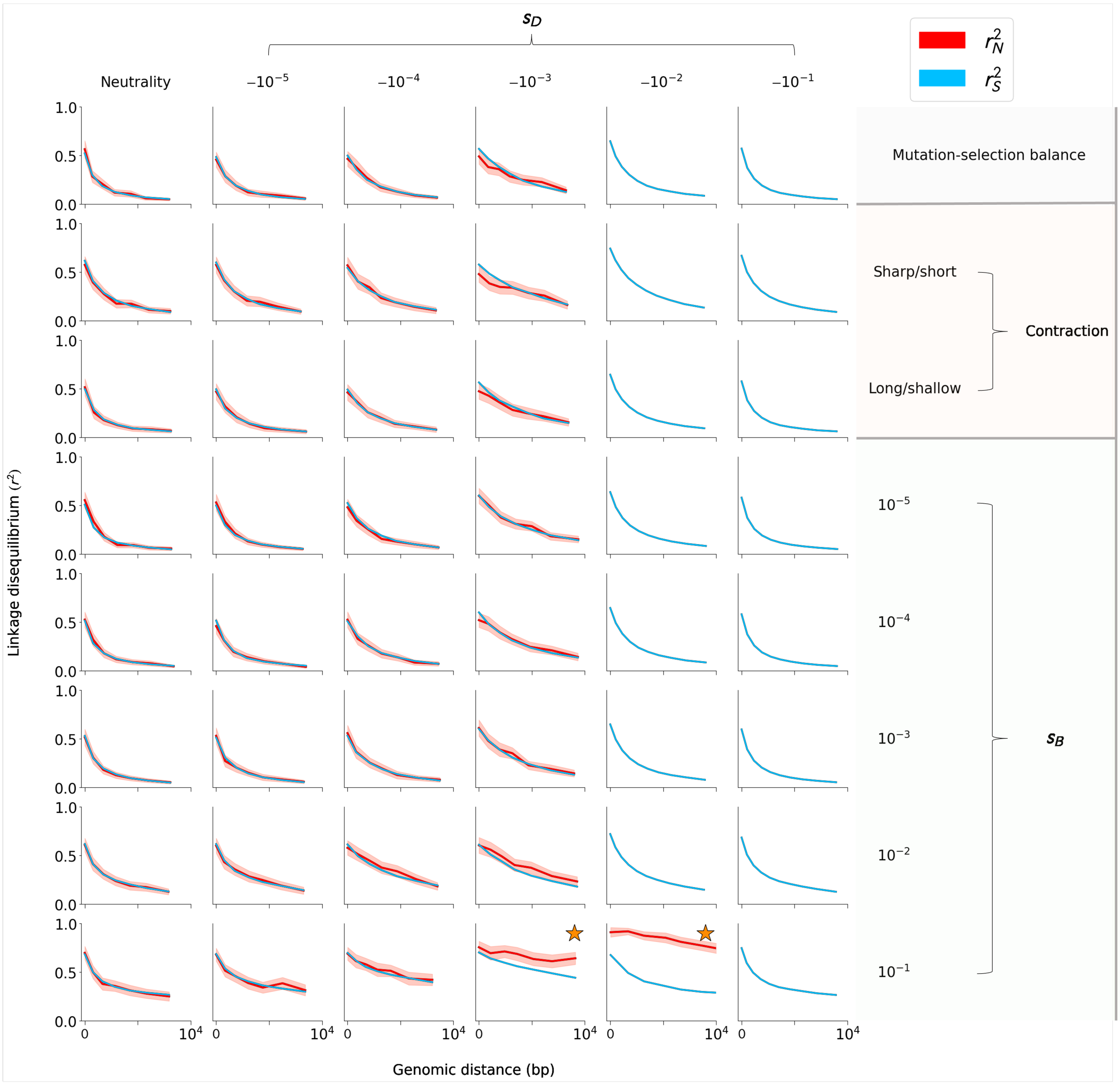

**Figure S3: $r_N^2$ and $r_S^2$ measured among common variants in a range of simulated evolutionary scenarios, where $\rho = 10^{-7}$.** Analogous to Figure **S1**, but for simulations performed with $\rho = 10^{-7}$ (i.e. $\rho/\mu = 10^{-1}$). As expected, LD decays much more slowly in this regime than with $\rho/\mu = 1$. Moreover, AUC($r_N^2 - r_S^2$) was significantly greater than zero only when $s_B = 10^{-1}$. While AUC($r_N^2 - r_S^2$) > 0 when $s_B = 10^{-2}$ and $s_D = -10^{-3}$, the difference was not significant.

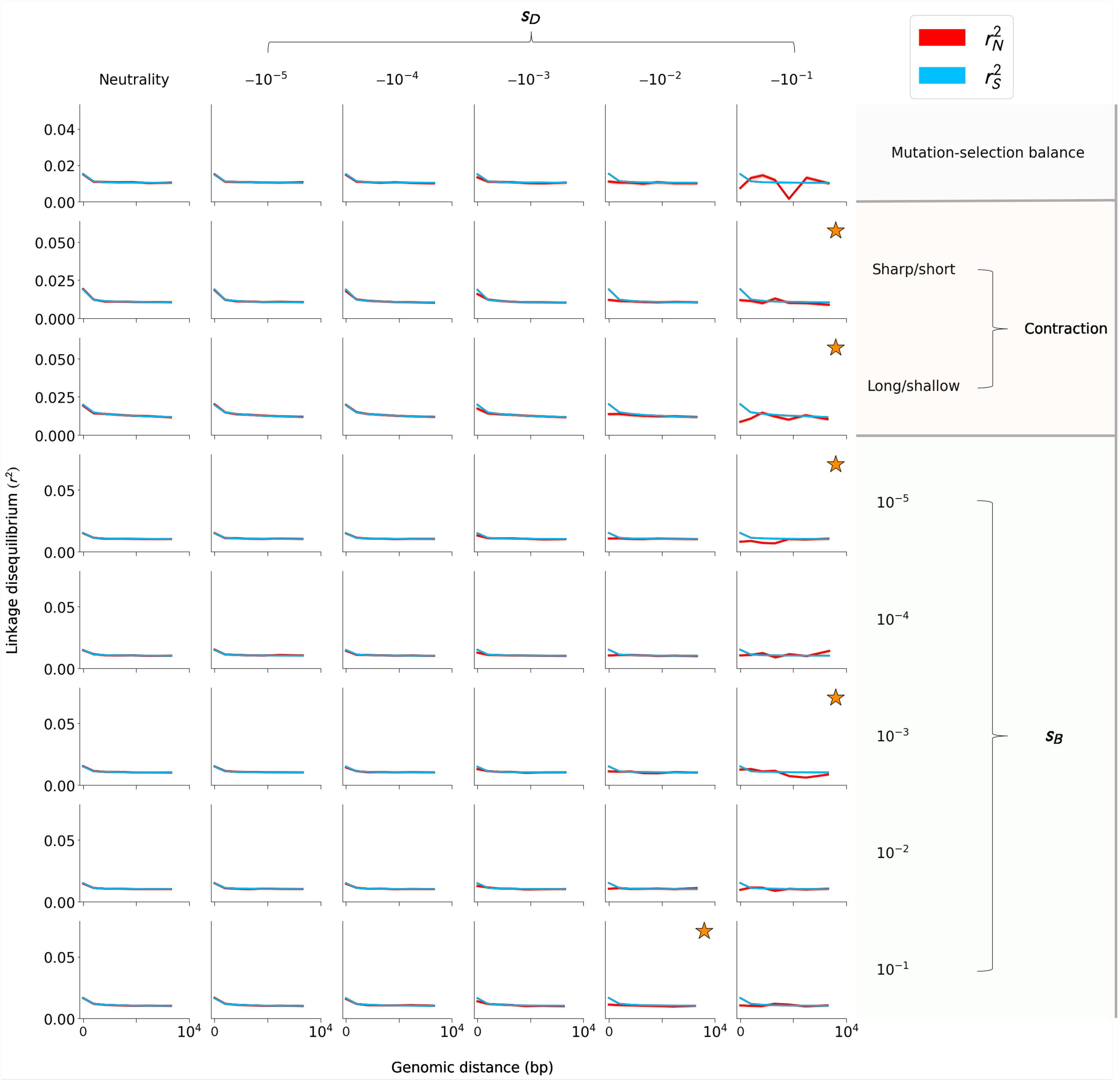

**Figure S4: $r_N^2$ and $r_S^2$ measured among rare variants in a range of simulated evolutionary scenarios, where $\rho = 10^{-5}$.** LD patterns among rare variants for the evolutionary scenarios discussed in the Main Text, with $\rho = 10^{-5}$ (i.e. $\rho/\mu = 10$). LD among both synonymous and non-synonymous variants decays extremely rapidly, and differences between $r_N^2$ and $r_S^2$ are typically only apparent over very short length scales.

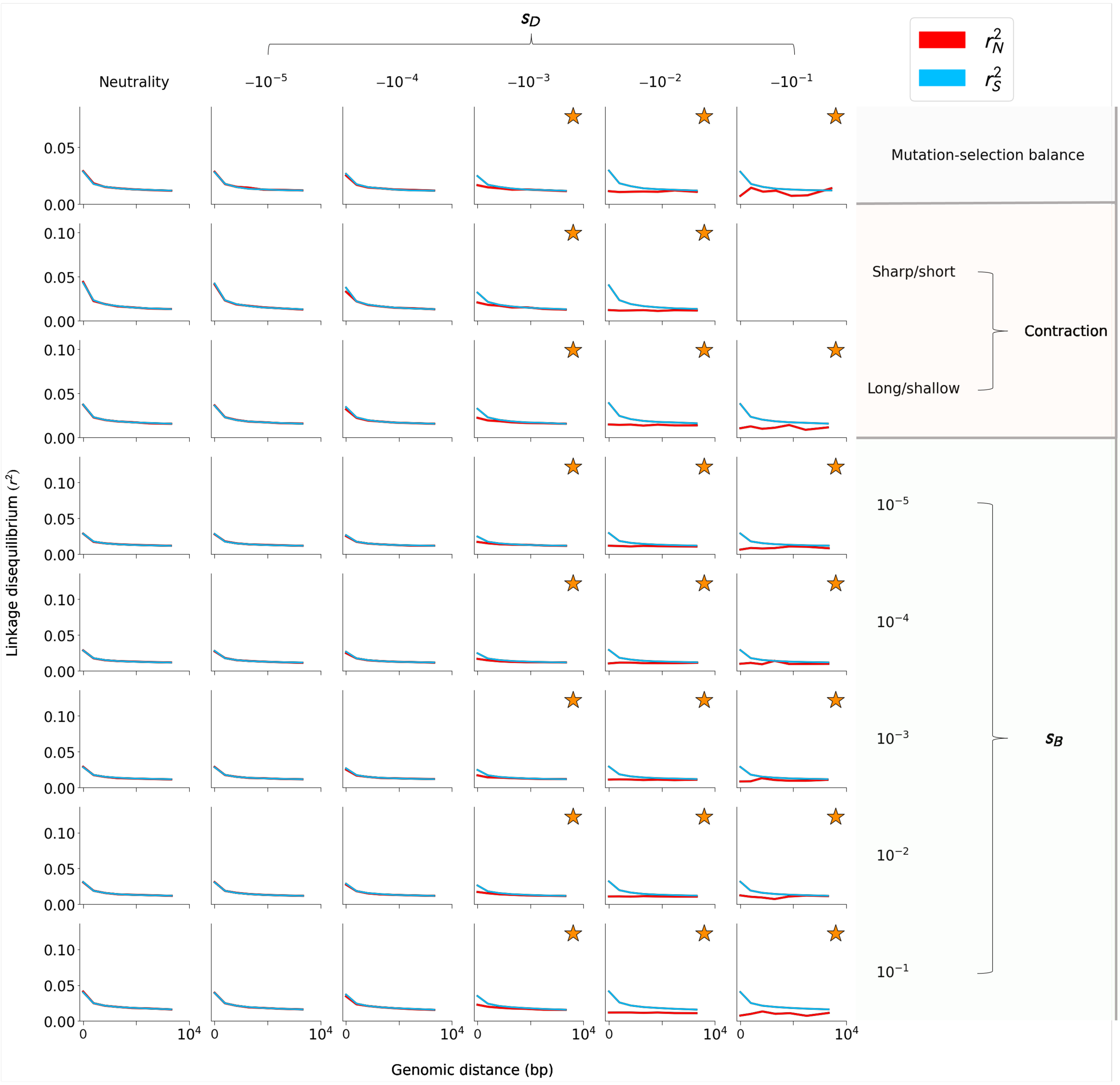

**Figure S5: $r_N^2$ and $r_S^2$ measured among rare variants in a range of simulated evolutionary scenarios, where $\rho = 10^{-6}$.** LD patterns among rare variants for the simulations discussed in the Main Text. AUC($r_N^2 - r_S^2$) is significantly less than zero in all simulations where $s_D > 1/N_e$ except for the short, sharp contraction with $s_D = -10^{-1}$, in which there were an insufficient number of both rare synonymous and non-synonymous variants to create LD curves.

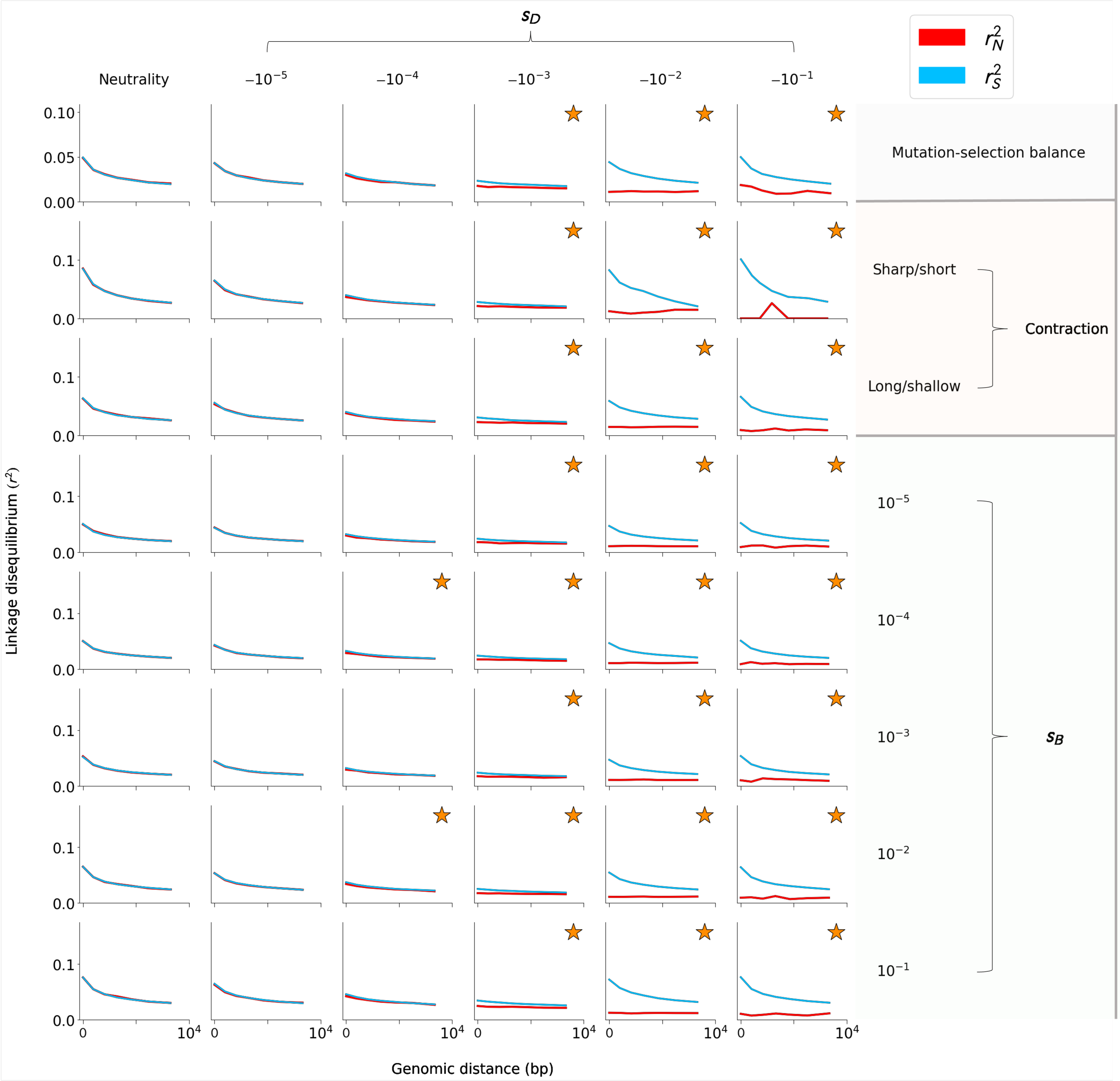

**Figure S6: $r_N^2$ and $r_S^2$ measured among rare variants in a range of simulated evolutionary scenarios, where $\rho = 10^{-7}$.** LD patterns among rare variants for the evolutionary scenarios discussed in the Main Text, with $\rho = 10^{-7}$ (i.e. $\rho/\mu = 10^{-1}$). AUC($r_N^2 - r_S^2$) is significantly less than zero in all simulations where $s_D > 1/N_e$, and in two instances also when $s_D = 1/N_e$.

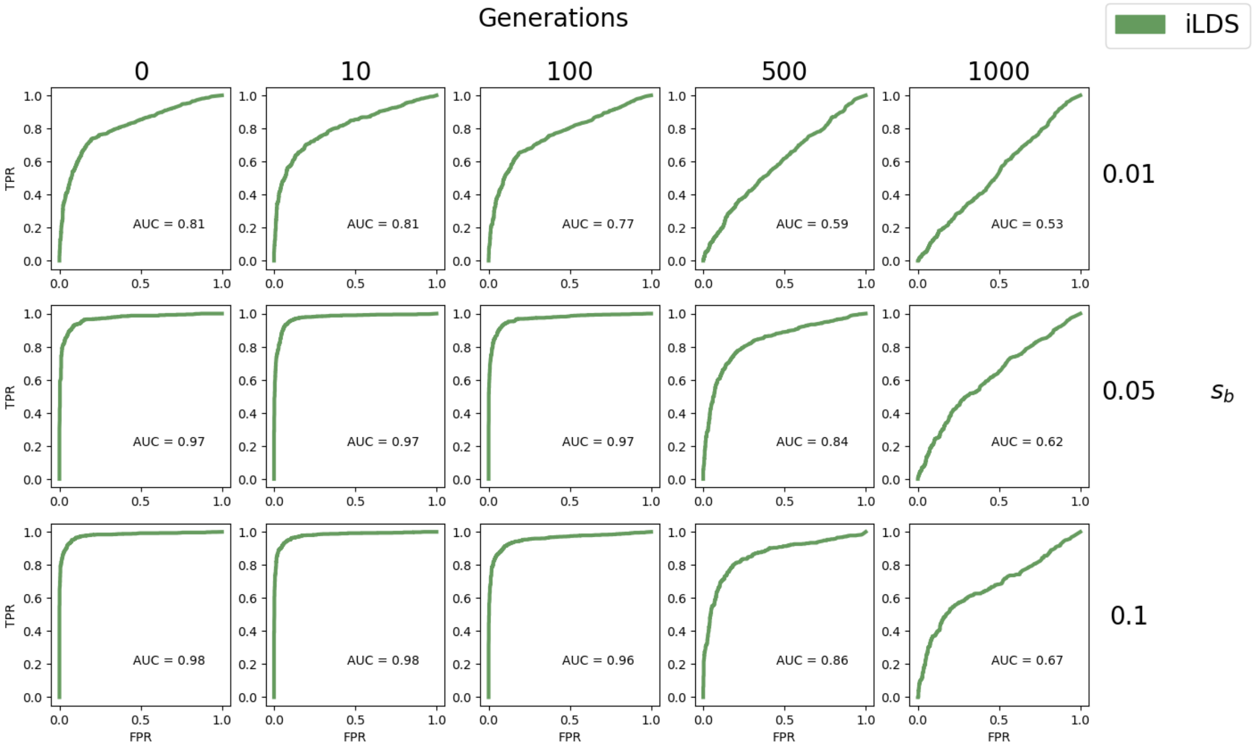

**Figure S7: Power analysis for iLDS under various sweep scenarios versus a constant $N_e$ population at mutation-selection balance**. Receiver operating characteristic (ROC) curves showing true (TPR) and false positive rates (FPR) of iLDS for sweeps of various strengths (rows) and number of generations since the sweep ended (columns). All populations experienced constant deleterious selection strength of $s_D = -10^{-3}$

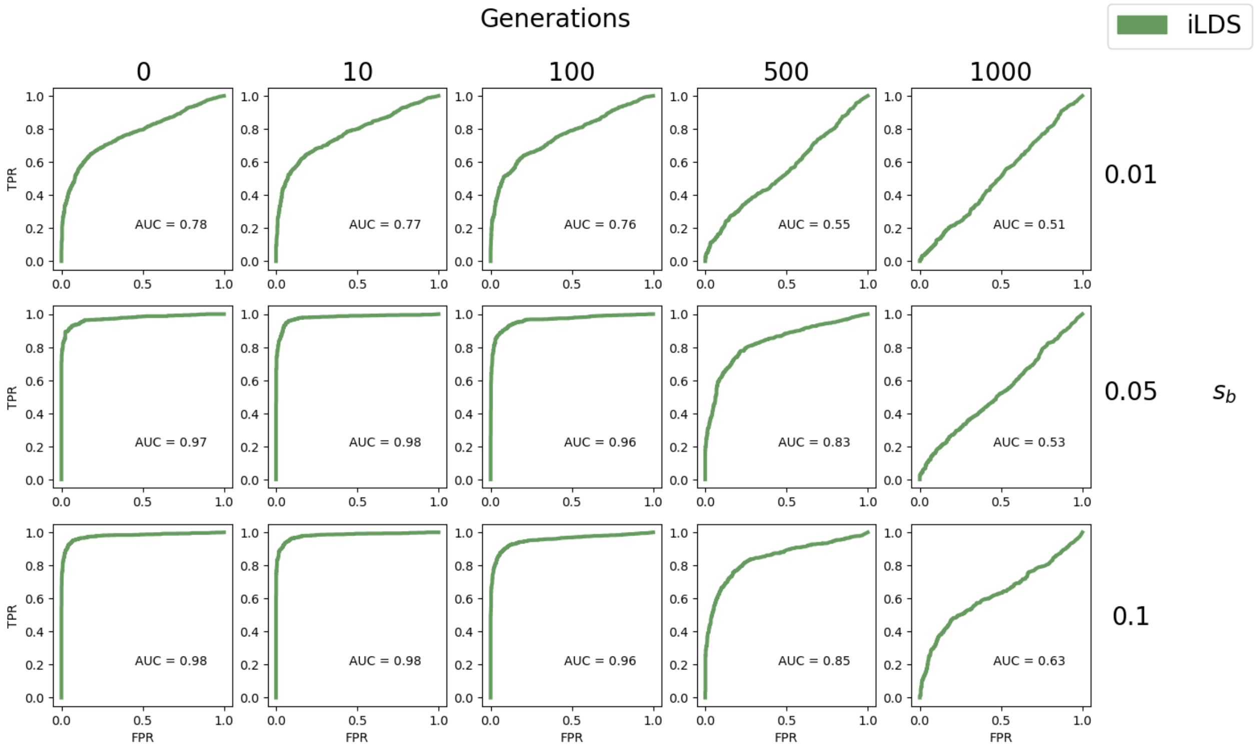

**Figure S8: Power analysis for iLDS under various sweep scenarios versus populations experiencing a sharp, short bottleneck under mutation-selection balance.** Receiver operating characteristic (ROC) curves showing true and false positive rates of iLDS for sweeps of various strengths (rows) and number of generations since the sweep ended (columns). Here, the false positive rate is assessed by the rate of mis-classifying populations experiencing a short, sharp bottleneck but no positive selection (see Section 5.3). All populations experienced constant deleterious selection strength of $s_D = -10^{-3}$.

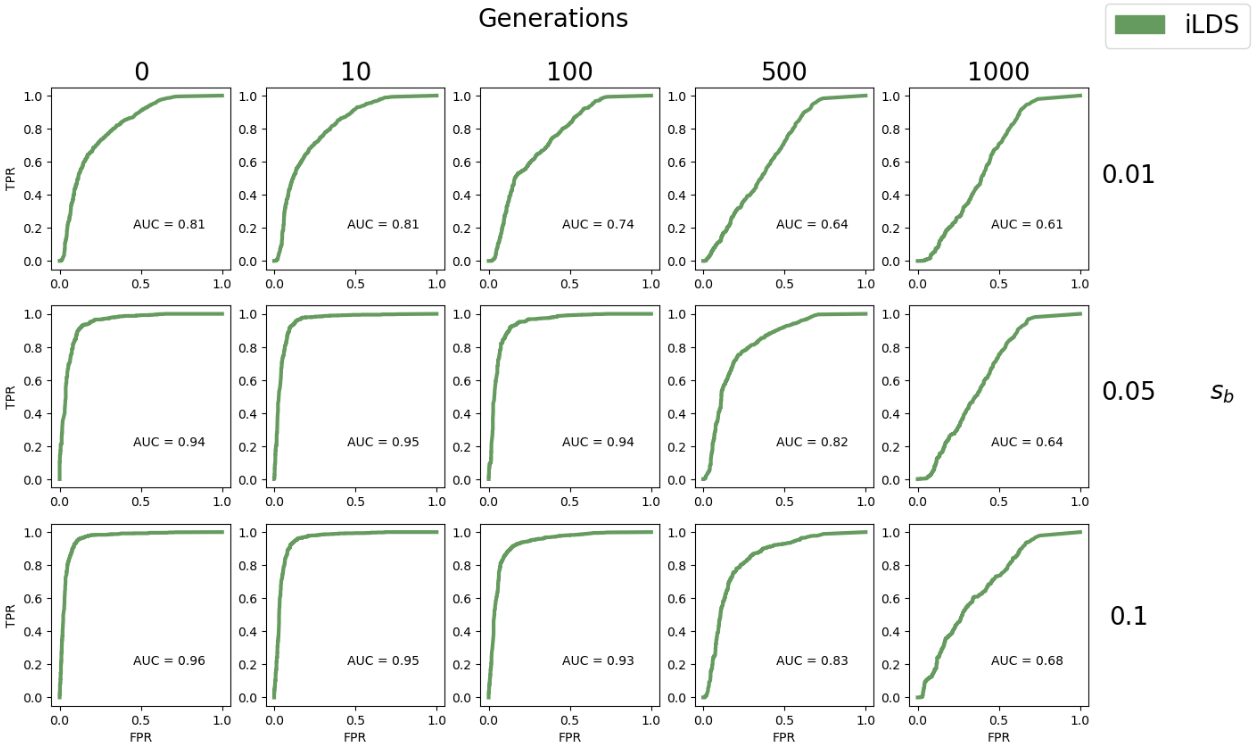

**Figure S9: Power analysis for iLDS under various sweeps scenarios versus populations experiencing long, shallow bottleneck**. Receiver operating characteristic (ROC) curves showing true and false positive rates of iLDS over a range of significance values, for sweeps of various strengths (rows) and number of generations since the sweep ended (columns). Here, the false positive rate is assessed by the rate of mis-classifying populations experiencing a long, shallow bottleneck but no positive selection (see Section 5.3). All populations experienced constant deleterious selection strength of $s_D = -10^{-3}$.

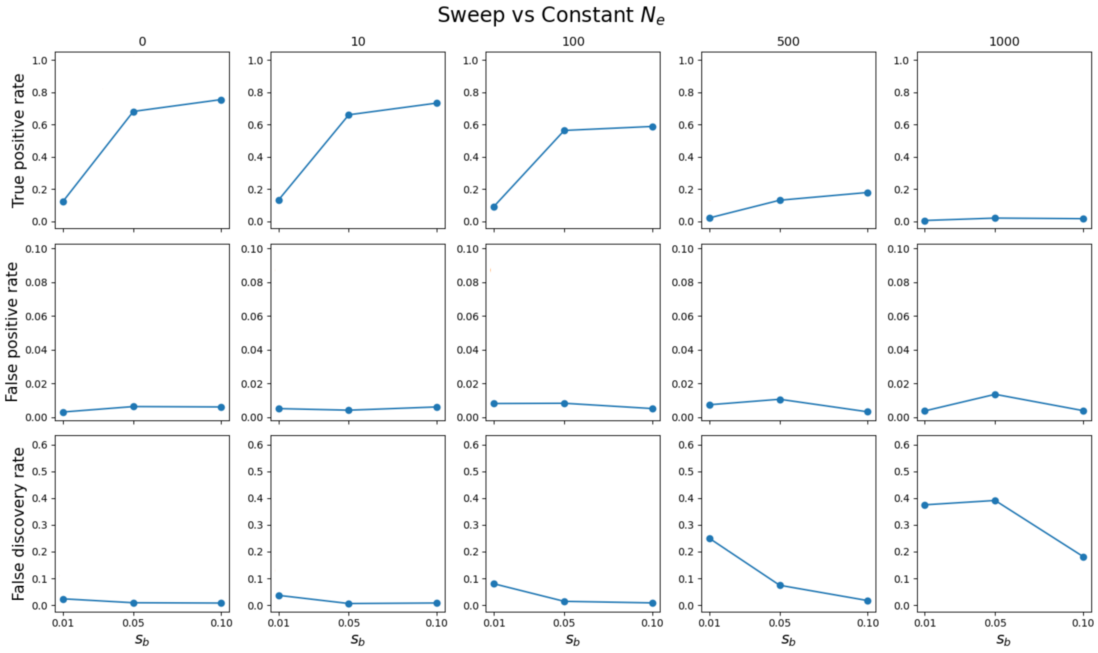

**Figure S10: True positive, false positive, and false discovery rates for sweeps against constant $N_e$.** True positive, false positive, and false discovery rates of iLDS at $\alpha = 0.05$ threshold when detecting selective sweeps against the genomic background of a population of constant size at mutation-selection balance.

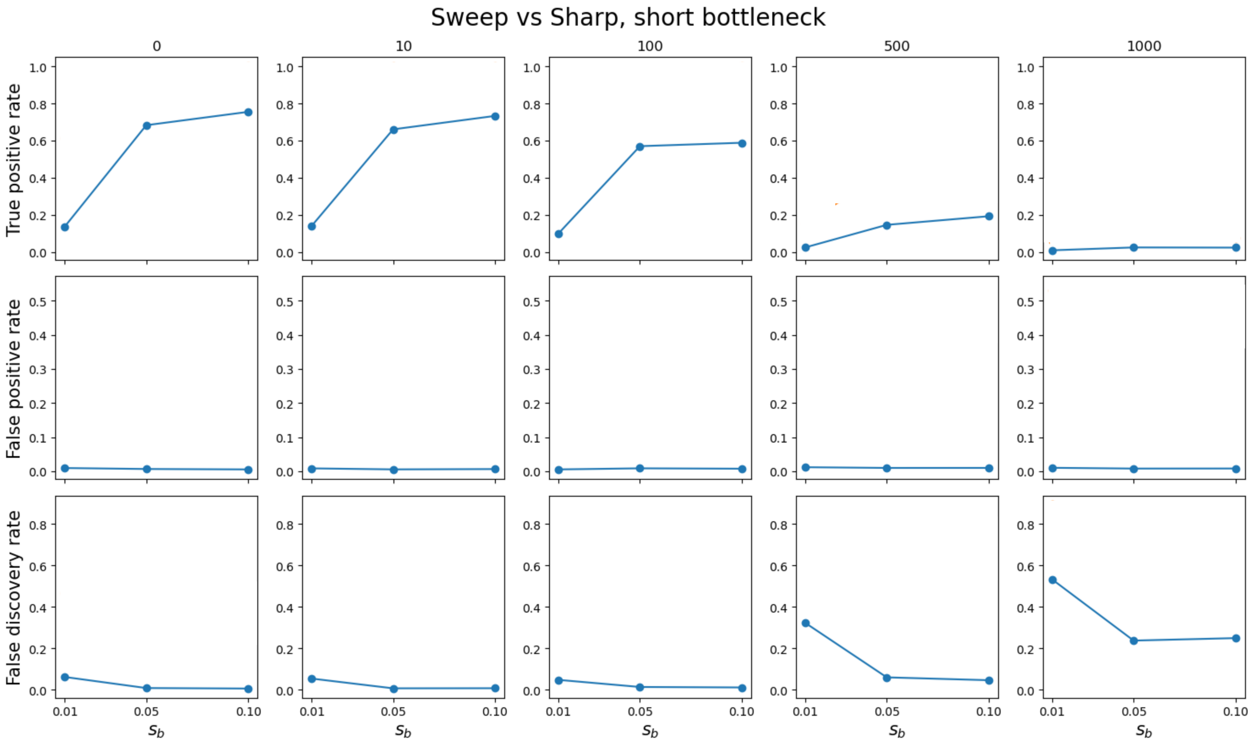

**Figure S11: True positive, false positive, and false discovery rates for sweeps against sharp, short bottleneck**. True positive, false positive, and false discovery rates of iLDS at $\alpha = 0.05$ threshold when detecting selective sweeps against the genomic background of a population experiencing a sharp, short bottleneck.

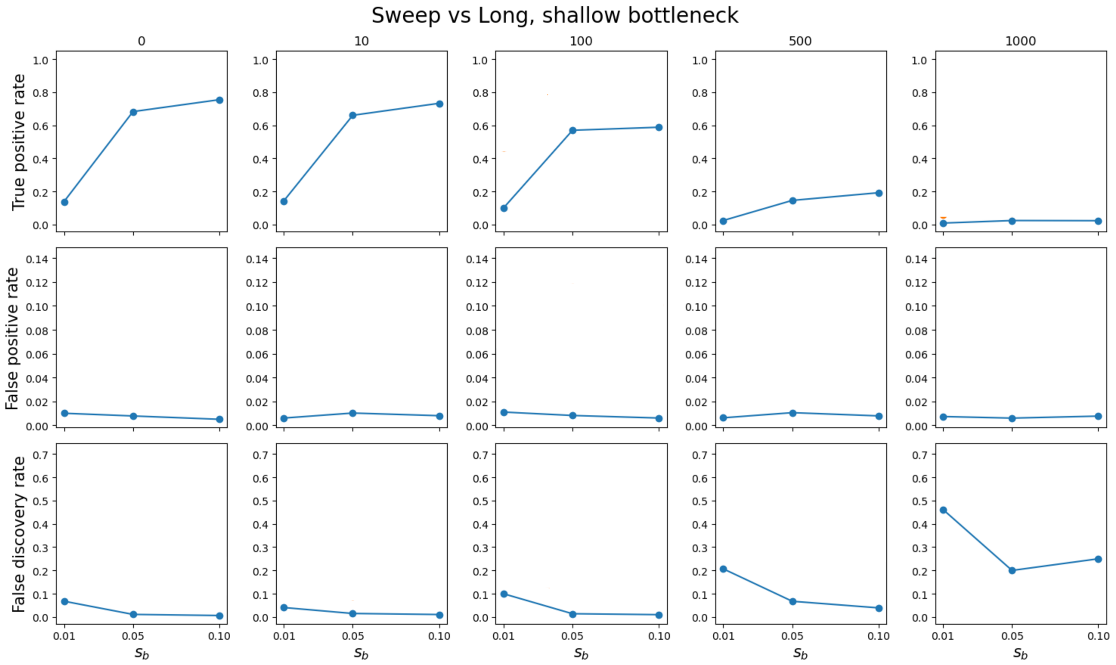

**Figure S12: True positive, false positive, and false discovery rates for sweeps against long, shallow bottleneck**. True positive, false positive, and false discovery rates of iLDS at $\alpha = 0.05$ threshold when detecting selective sweeps against the genomic background of a population experiencing a long, shallow bottleneck.

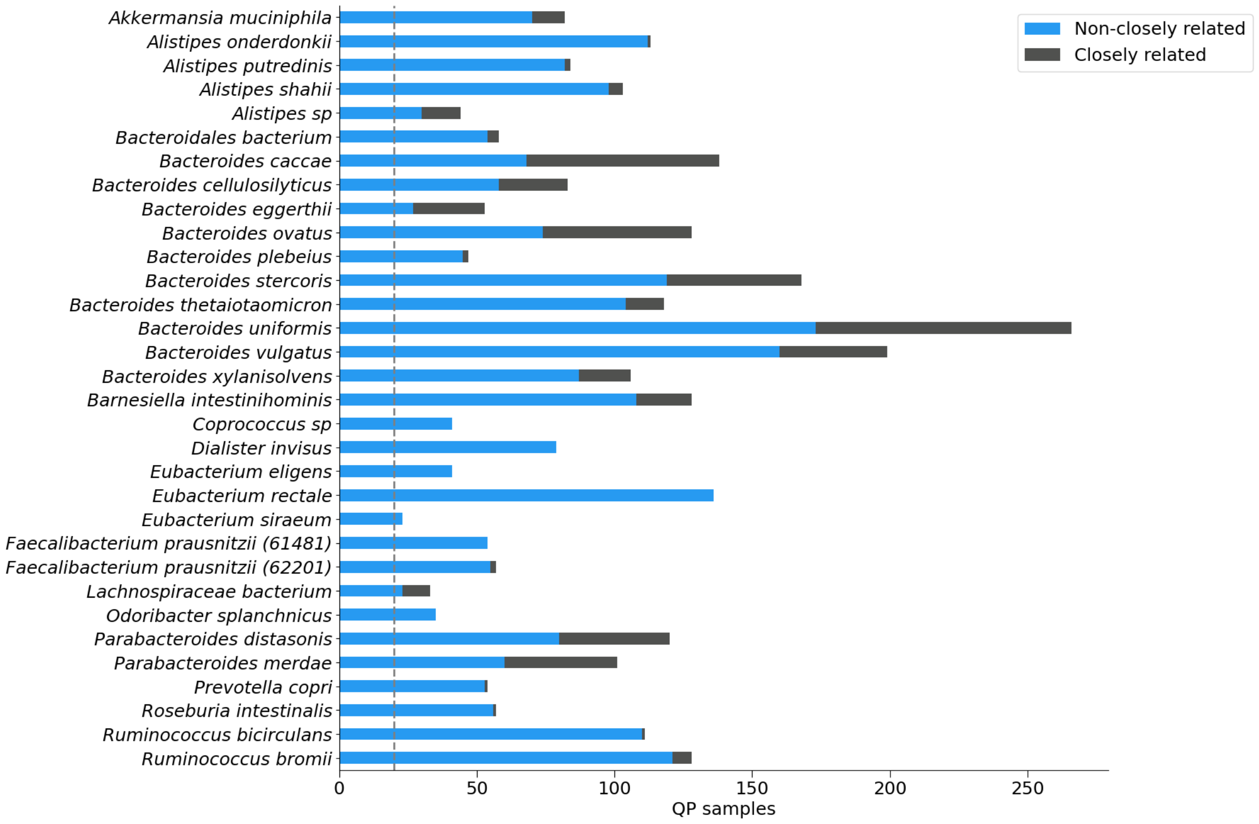

**Figure S13: Number of QP samples by species.** The number of QP samples (Supplementary Section 3.4) belonging to the largest clade for each of the 32 species analyzed. Samples are divided between closely related ($d < 5 \times 10^{-4}$ at 4D sites) and non-closely related samples.

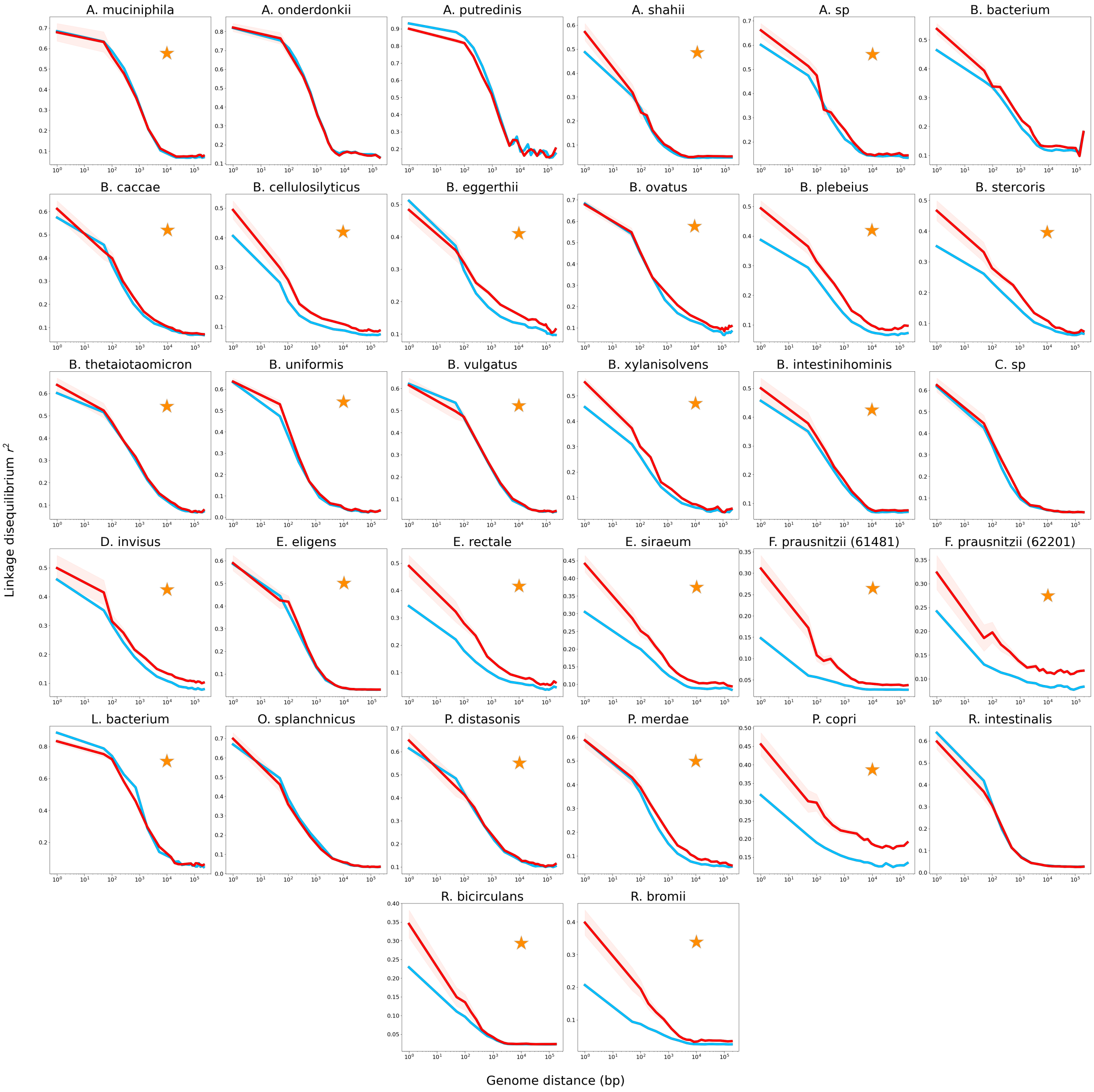

**Figure S14: $r_N^2$ and $r_S^2$ measured in prevalent commensal gut microbiota for common variants.** This figure is analogous to Figure 2A (top panel) in the Main Text. $r_N^2$ is shown in red, while $r_S^2$ is shown in blue. While $r_N^2 < r_S^2$ at certain distances in some species (e.g. at short distances in *Alistipes putredinis*, overall $r_N^2$ typically exceeds $r_S^2$ for all species.

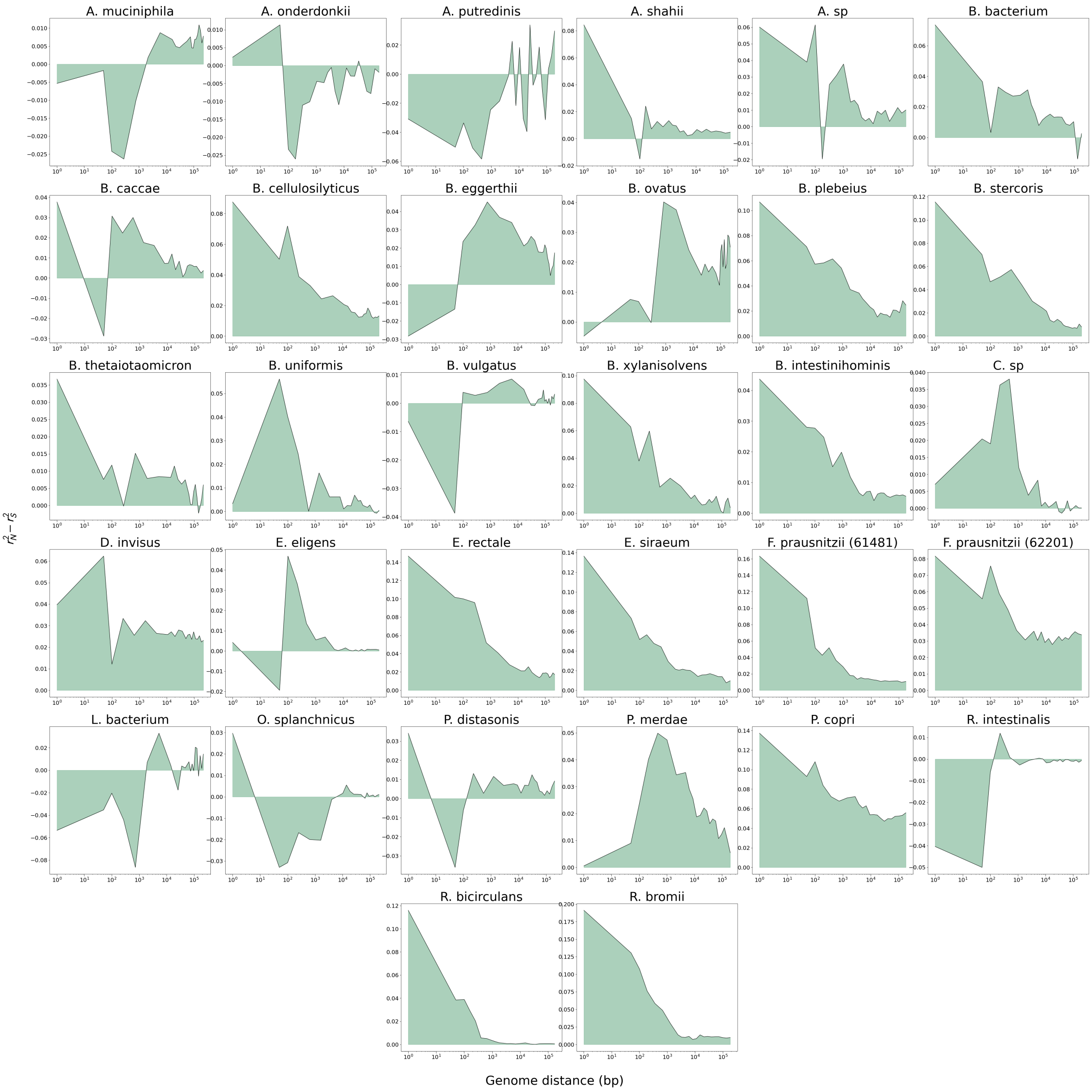

**Figure S15:** $(r_N^2 - r_S^2)$ **measured in prevalent commensal gut microbiota for common variants.** Here, the function $(r_N^2 - r_S^2)$ is plotted (black line), and AUC$(r_N^2 - r_S^2)$ is shown with the green shaded region. In a majority of species, $(r_N^2 - r_S^2)$ tends to decrease as a function of distance, or reach a maximum at some intermediate distance. A minority of species (such as *Akkermansia municiphilia*, top left) show an increase of $(r_N^2 - r_S^2)$ with distance.

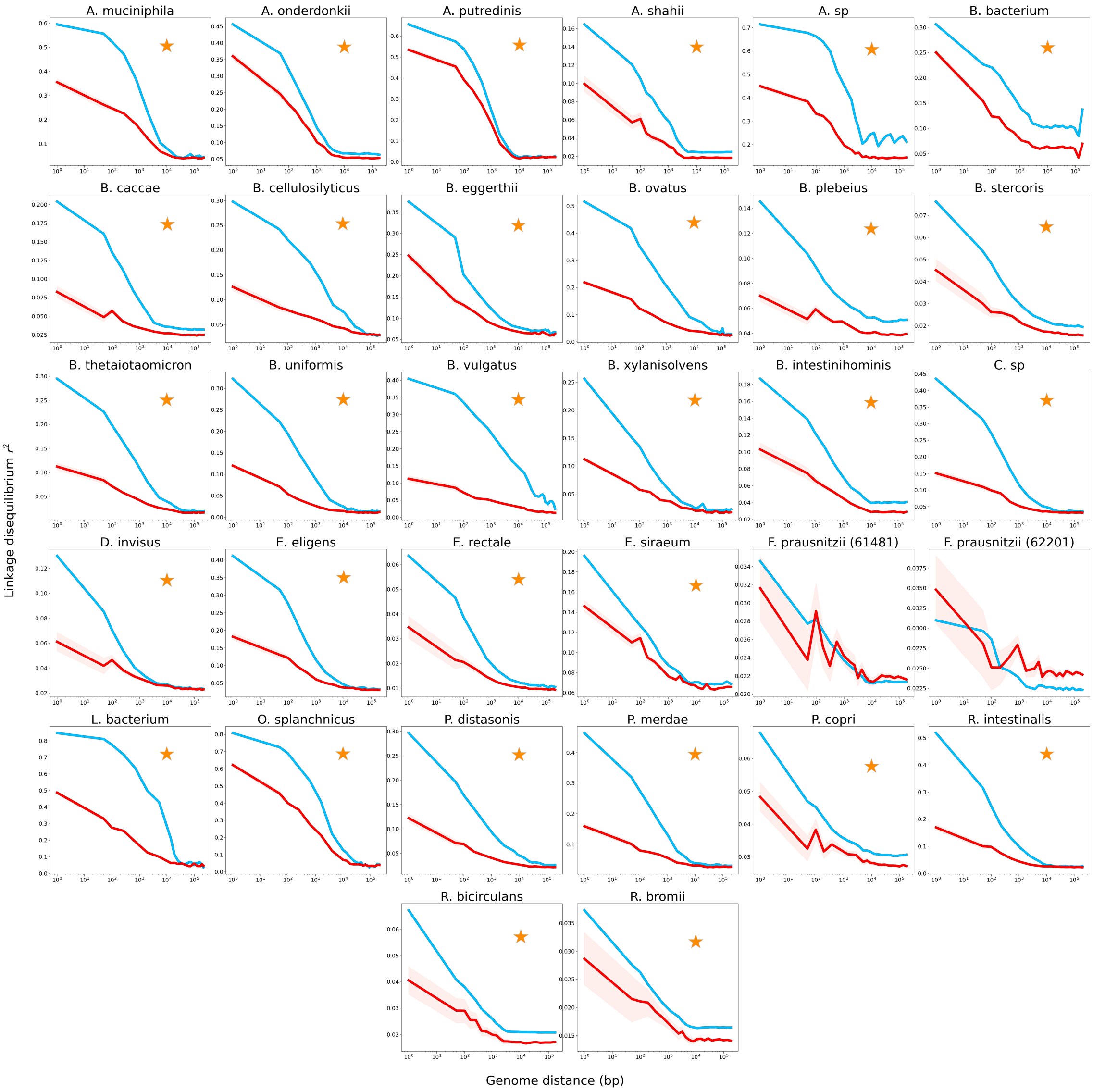

**Figure S16: $r_N^2$ and $r_S^2$ measured in prevalent commensal gut microbiota for rare variants.** This figure is analogous to Figure 2A (bottom panel) in the Main Text. $r_N^2$ is shown in red, while $r_S^2$ is shown in blue. $r_S^2$ is significantly elevated above $r_N^2$ in all but one species.

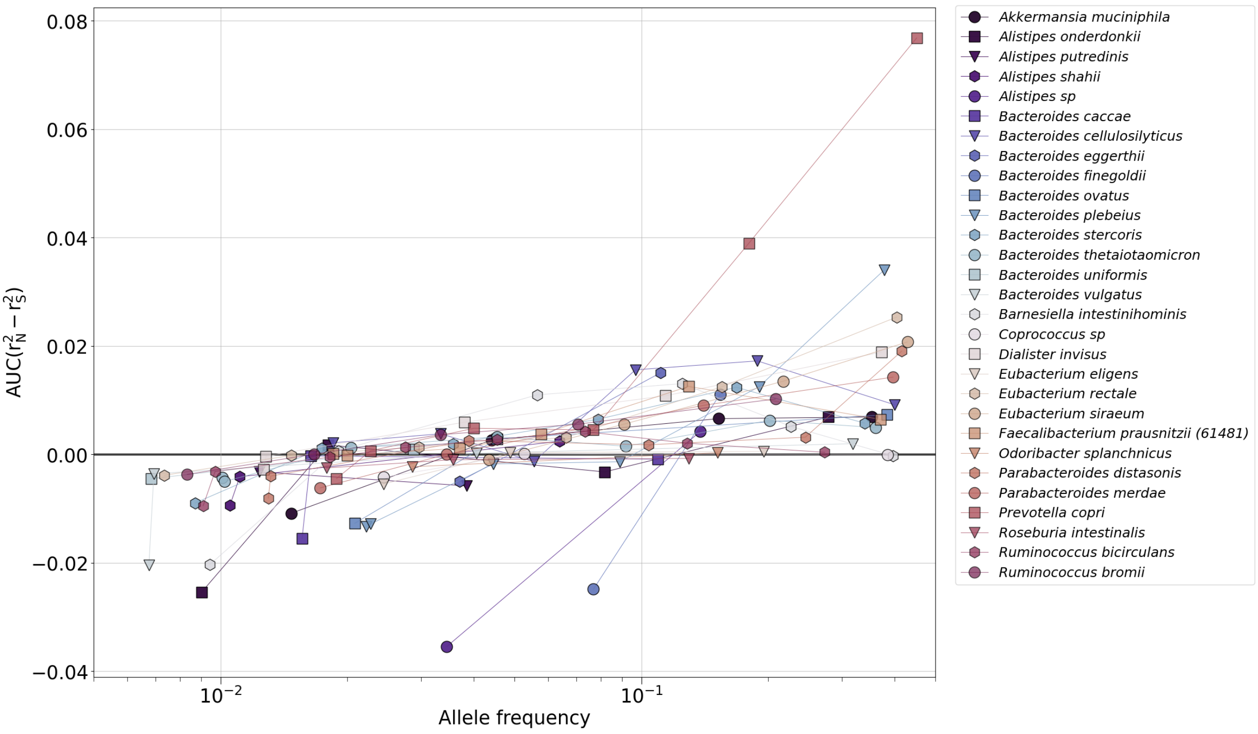

**Figure S17: AUC($r_N^2 - r_S^2$) measured in prevalent commensal gut microbiota for varying allele frequencies.** For each species, all variants were binned by allele frequency interval, and AUC($r_N^2 - r_S^2$) calculated for each bin. AUC($r_N^2 - r_S^2$) is universally an increasing function of allele frequency, with a transition between negative AUC($r_N^2 - r_S^2$) and positive AUC($r_N^2 - r_S^2$) typically occuring around $f \approx 0.05$.

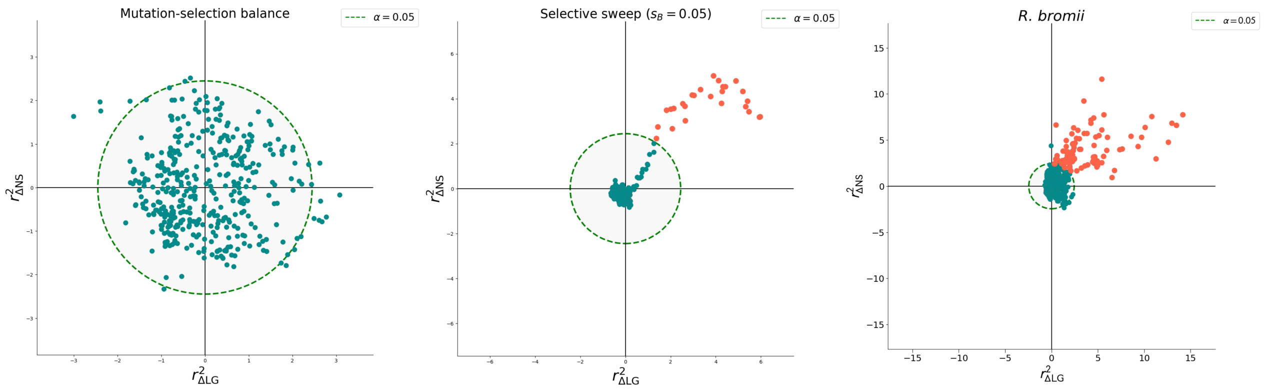

**Figure S18: $r^2_{\Delta\text{NS}}$ versus $r^2_{\Delta\text{LG}}$ in simulations and data.** Here, we plot the two components of iLDS ($\bar{r}^2_{\Delta\text{NS}}$ and $\bar{r}^2_{\Delta\text{LG}}$, the normalized differences between non-synonymous versus synonymous LD and local versus genome-wide LD) for simulations and data. Significant windows are colored orange, and non-significant windows green. Because the formula for iLDS is $\left(\bar{r}^2_{\Delta(NS)}\right)^2 + \left(\bar{r}^2_{\Delta(LG)}\right)^2$, the value of the statistic can be thought of as the square of the distance from the origin of a point in ($\bar{r}^2_{\Delta\text{NS}}$, $\bar{r}^2_{\Delta\text{LG}}$) space. The green dashed line is a circle with a radius corresponding to the square root of the $95^{\text{th}}$ percentile of a $\chi^2$ distribution with two degrees of freedom—an iLDS value exceeding this threshold is necessary but not sufficient for a window to be called as significant. In a simulation at mutation-selection balance (left) the iLDS statistic may surpass the critical threshold, but the additional checks of significance (non-synonymous LD exceeding synonymous LD, local LD exceeding genome-wide LD) prevent false inferences of selection. By contrast, in a simulated selective sweep (center) and real data (right), there is a notable positive correlation between $\bar{r}^2_{\Delta\text{LG}}$ and $\bar{r}^2_{\Delta\text{NS}}$ in a tail of windows which also tend to be called as significant.

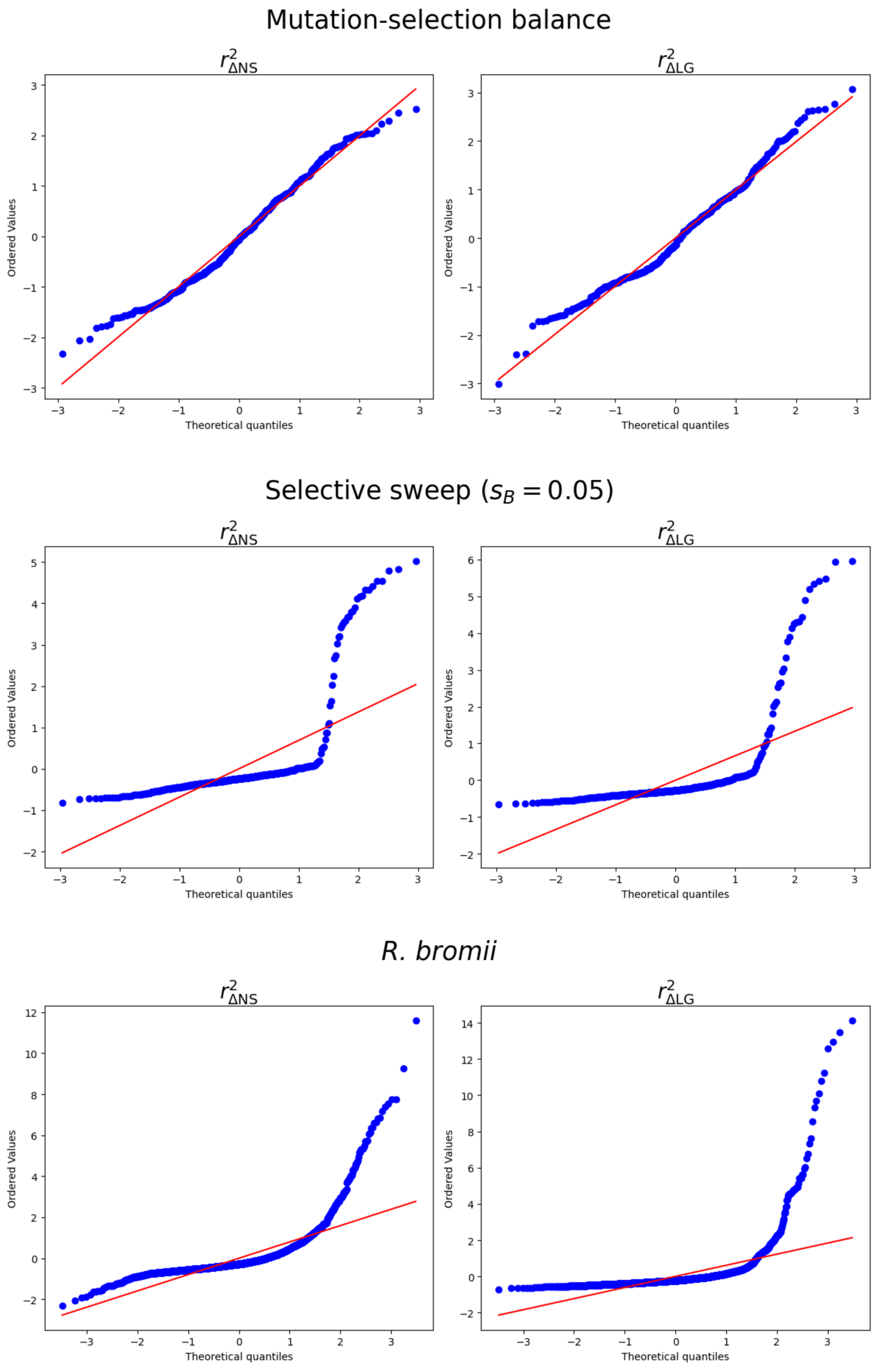

**Figure S19: QQ plots of $\bar{r}^2_{\Delta \mathrm{NS}}$ versus $\bar{r}^2_{\Delta \mathrm{LG}}$ in simulations and data.** Quantile-quantile plots showing the quantiles of $\bar{r}^2_{\Delta \mathrm{NS}}$ and $\bar{r}^2_{\Delta \mathrm{LG}}$ versus the expected quantiles if each statistic had a standard normal distribution for a simulation at mutation-selection balance (top), a simulated selective sweep (middle), and real data for *R. bromii* (bottom). While each statistic roughly follows a standard normal distribution at mutation-selection balance, the simulated selective sweeps and real data exhibit substantial departures from the mutation-selection balance scenario, particularly driven by a long right tail of large $\bar{r}^2_{\Delta \mathrm{NS}}$ and $\bar{r}^2_{\Delta \mathrm{LG}}$ values.

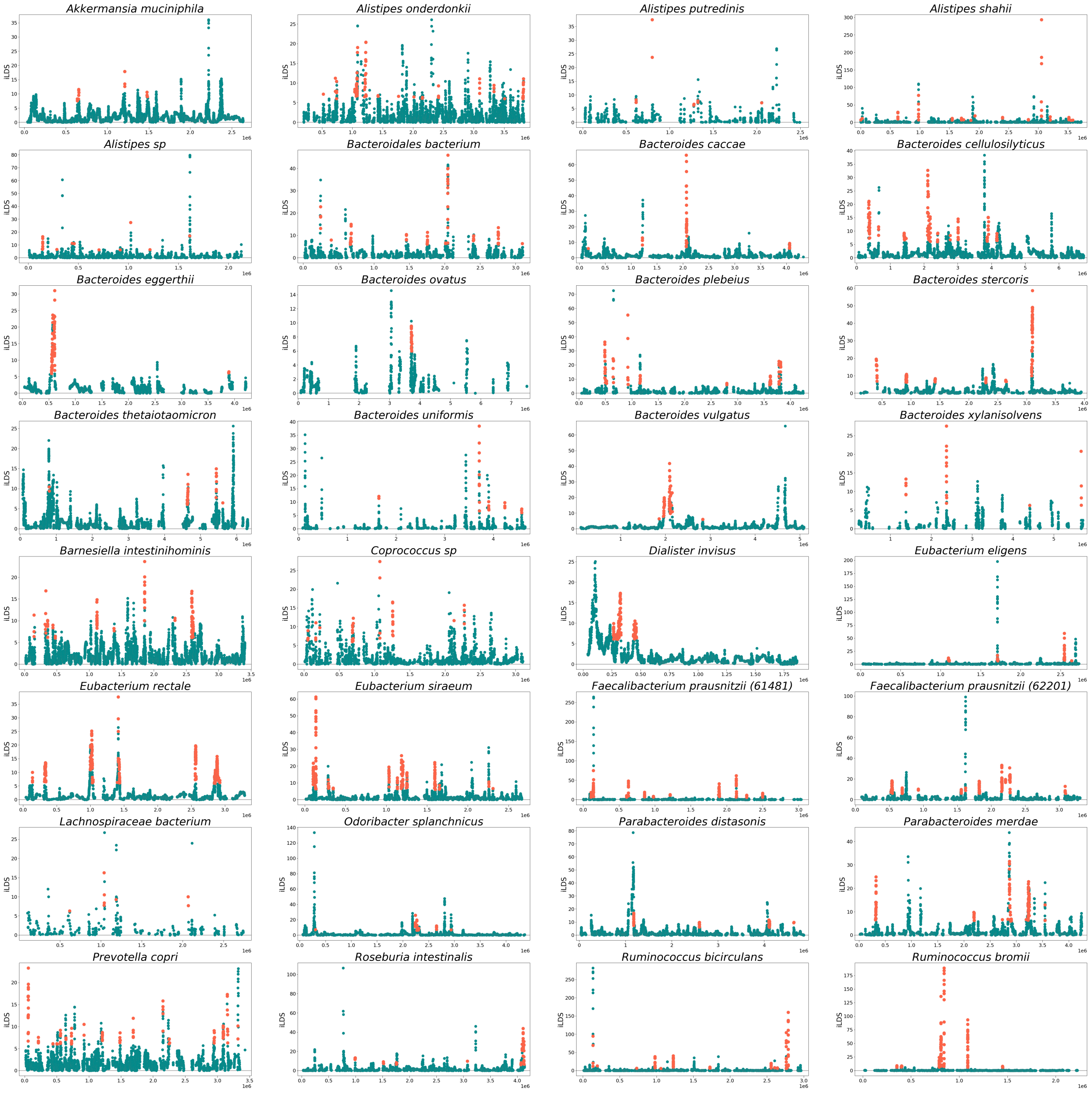

**Figure S20: iLDS scans for all species analyzed (HMP).** Analogous to Figure 3B, iLDS scans are shown here for all 32 species analyzed in our dataset (metagenomes that we quasiphased). Significant windows are shown in orange, while non-significant windows are shown in green.

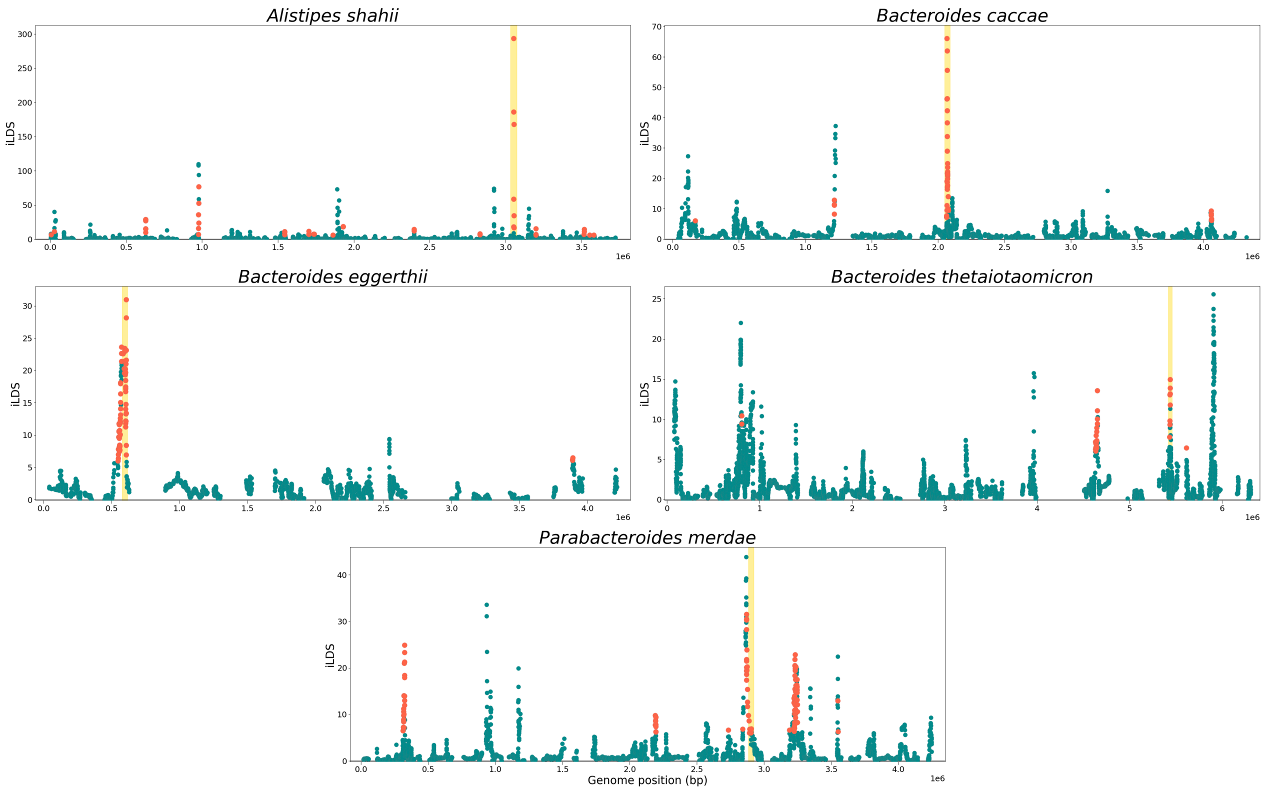

**Figure S21: *susC/susD* genes under selection.** *susC/susD* genes were detected as being under selection in five unique species. Here, the iLDS scans for these species are plotted, and the location of the *susC/susD* locus under selection in that species is shown with a gold bar.

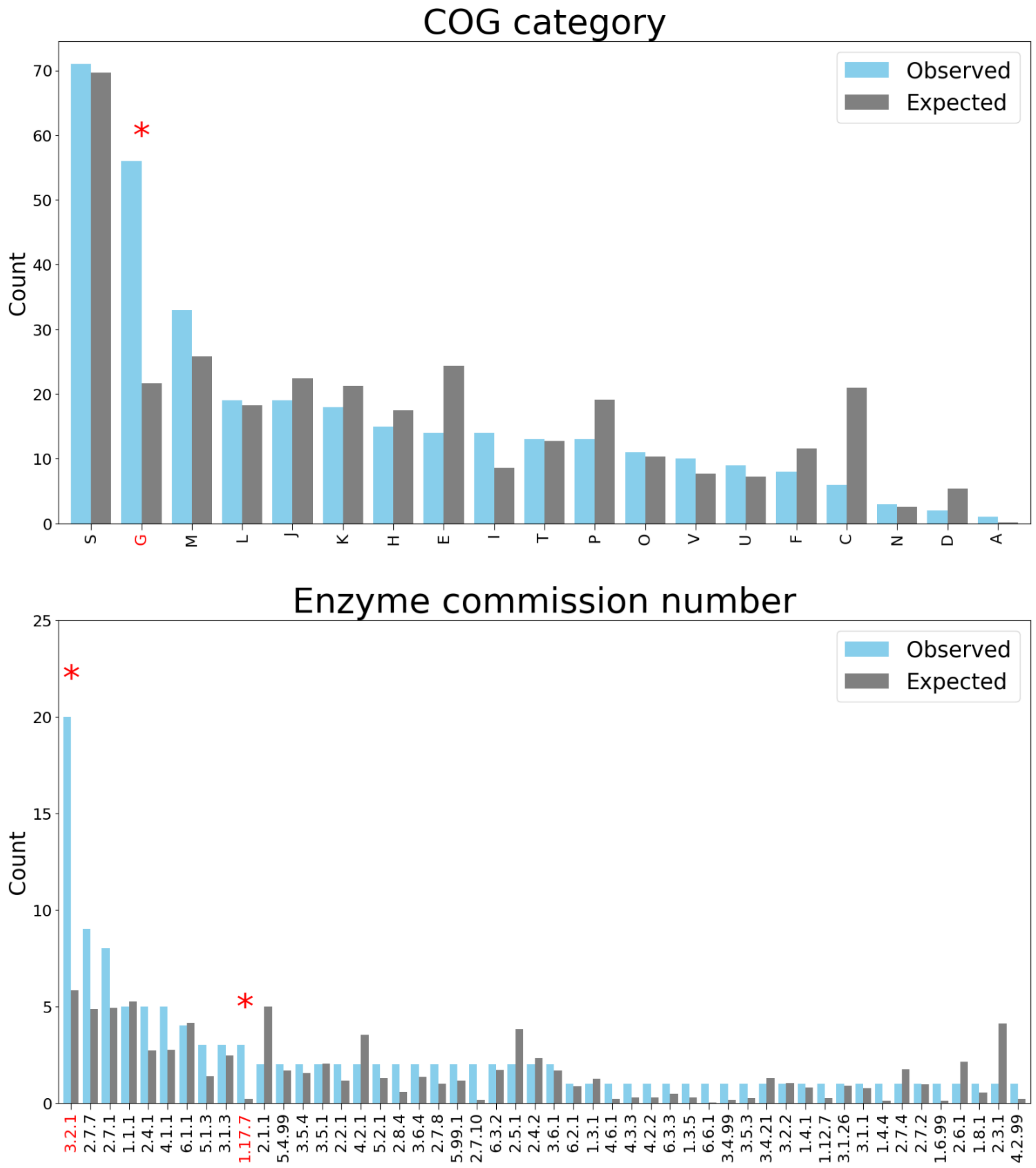

**Figure S22: Enrichment of functional categories among sweeps.** Number of genes expected to be under selection, based on the frequency of such genes across the core genomes of the 32 species analyzed, versus the number of genes observed to be under selection for two kinds of functional annotations: COG categories (top) [7] and Enzyme Commission (EC) numbers (bottom) [8]. Functional categories exhibiting a statistically significant enrichment among selected genes are denoted by red asterisks and labels (see Section 6 below for details of significance test). Among COG categories (top), we observed a statistically significant enrichment of genes related to carbohydrate metabolism and transport (category G). Among ECs, we observed an enrichment of glycoside hydrolases (EC 3.2.1) and oxidoreductases using ferredoxin as an electron acceptor (EC 1.17.7).

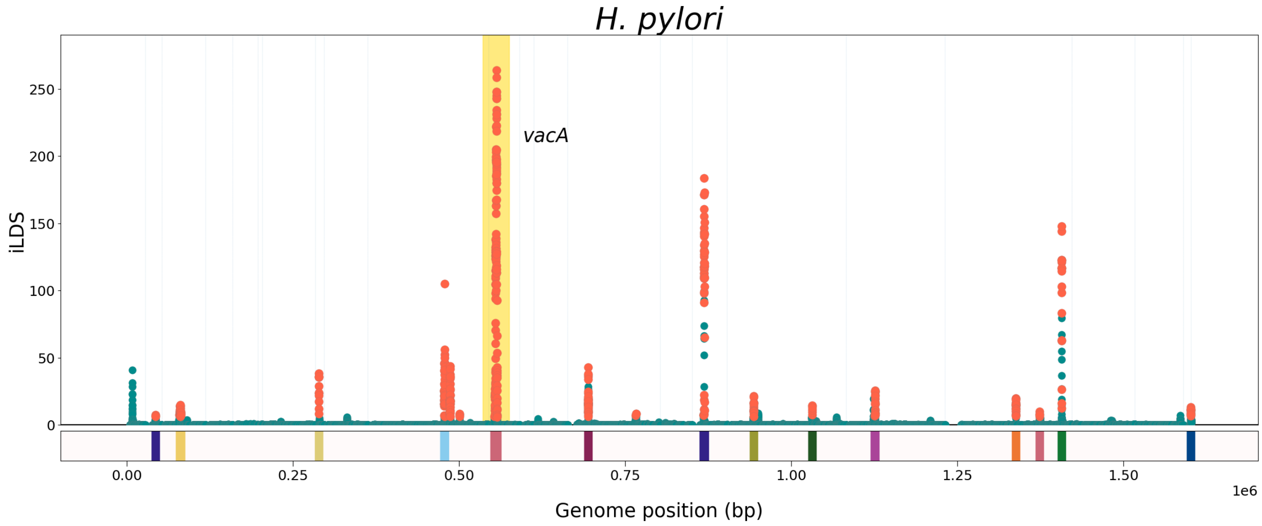

**Figure S23: iLDS scan of *H. pylori*.** iLDS scan for the highly recombinant gastric pathogen *Helicobacter pylori*. Much like *C. difficile* (**Figure 3A** Main Text), iLDS detects that a key virulence factor (here, the *vacA* gene, which encodes the vacuolating cytotoxin) is under positive selection in this species. Previous studies have likewise reported that *vacA* is experiencing positive selection [9, 10].

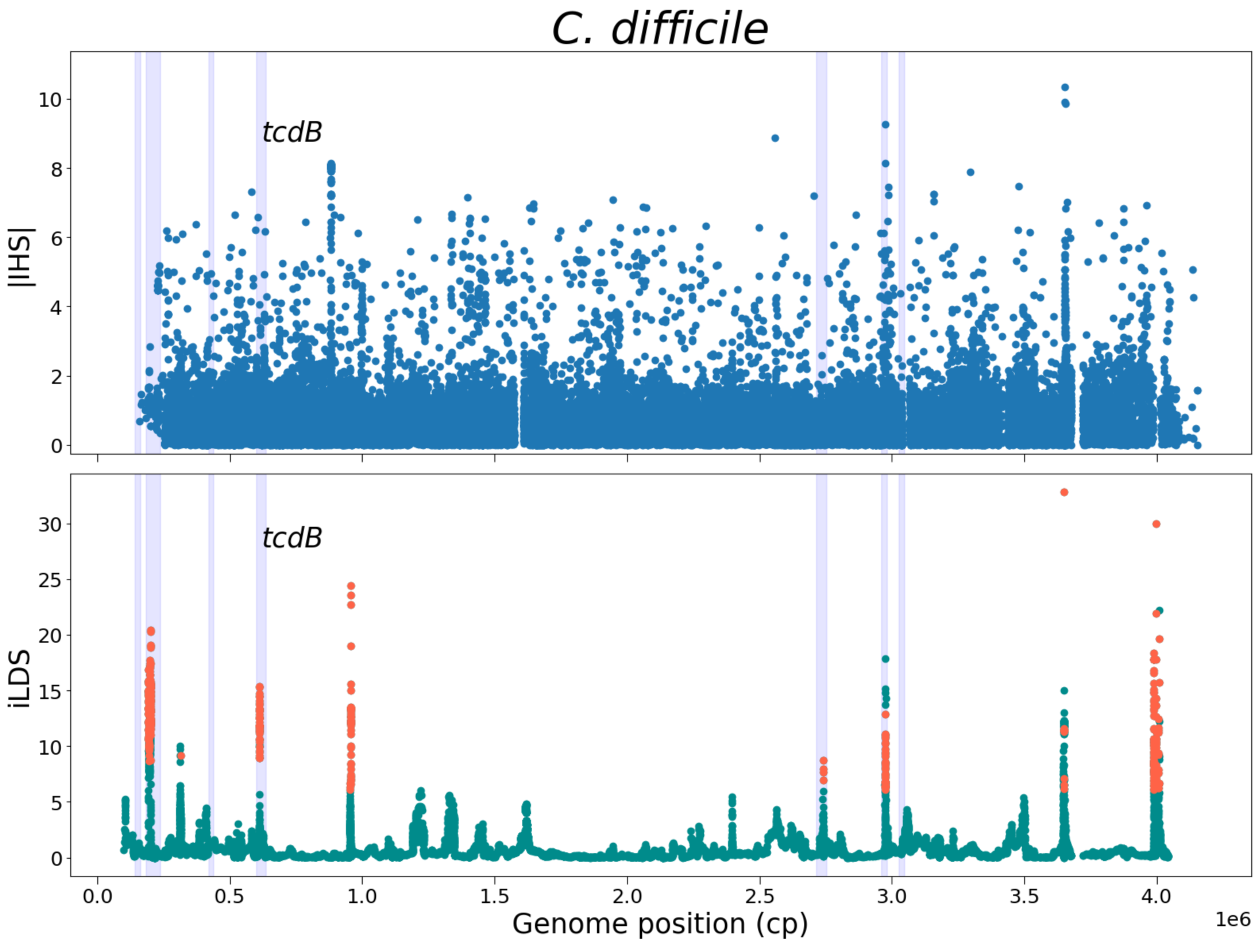

**Figure S24: Comparison iHS and iLDS scan of *C. difficle*.** We compare the performance of iLDS with iHS [11], a standard selection scan which detects extended haplotype homozygosity, using the same set of isolates of *C. difficle* as in Main Text **Figure 3A**. In constrast to the iLDS scan, iHS peaks are less sharply defined. In particular, iHS shows no clear signature of a selective sweep at the positive control *tcdB* locus or at other virulence factor loci, highlighted in light blue. iHS was implemented using the scikit-allel software [12], with values normalized in 10% allele frequency bins. Application of iHS to bacterial genomes may be challenging because iHS natively requires highly complete genomes with little to no missing data or few accessory genes. Concretely iHS looks for extended haplotype homozygosity on either side of a central SNP, and the absence of a gene would hinder the ability to detect these stretches. iLDS, by contrast, does not require such complete genomes given that it is applied in windows.

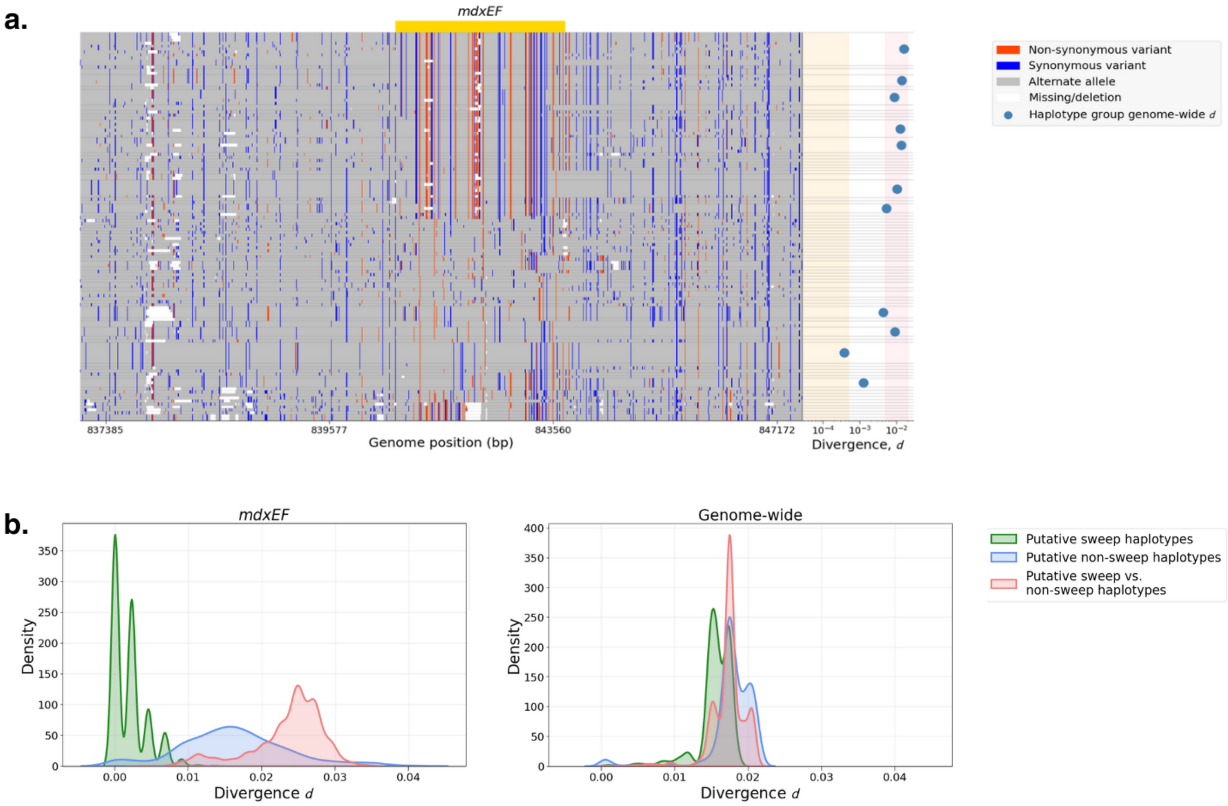

**Figure S25: Genomic diveristy at the *mdxEF* locus versus genome-wide for the species *R. bromii*.** **(A)** Haplotype plot of a ~10Kb region surrounding the sweep candidate *mdxEF* genes. Non-synonymous variants are colored red, while synonymous variants are colored blue, and missing sites are colored white. Horizontal lines separate strains into haplotype groups that are genetically identical at *mdxEF*. Haplotypes are ordered based on their genetic distance to the largest haplotype at the *mdxEF* locus. To aid in visualization, intermediate frequency variants (MAF $> 0.2$) are colored based on the identity of the first strain in the first haplotype group (i.e. the top row), while the color of all other sites is determined by minor allele at that position. The alternate allele at each site assigned by this polarization scheme is colored gray. At right, mean genome-wide divergence $d$ within each haplotype group containing three or more strains is shown with blue dots, while the pink region shows the typical range of $d$ (within a factor of two of the mean), and the orange region denotes the region $d < 5 \times 10^{-4}$/bp, indicative of close-relatedness among strains. **(B)** For the species *R. bromii*, divergence $d$ at 4D sites is shown at *mdxEF* (left) and genome-wide (right). The distribution of $d$ among samples is shown for the putatively sweeping haplotype group in green (the 26 haplotype groups at the top in **(A)**, above), for the remaining haplotype groups in blue, and between these two groups in red. At *mdxEF*, the putatively sweeping haplotype groups have very low $d$ relative to their genome-wide average, while the remaining haplotype groups have $d$ comparable to their genome-wide averge (though with greater variance, as is expected given that $d$ is calculated over far fewer sites). Moreover, the putatively sweeping haplotypes are far more genetically distinct from the remaining haplotypes at *mdxEF* than genome-wide.

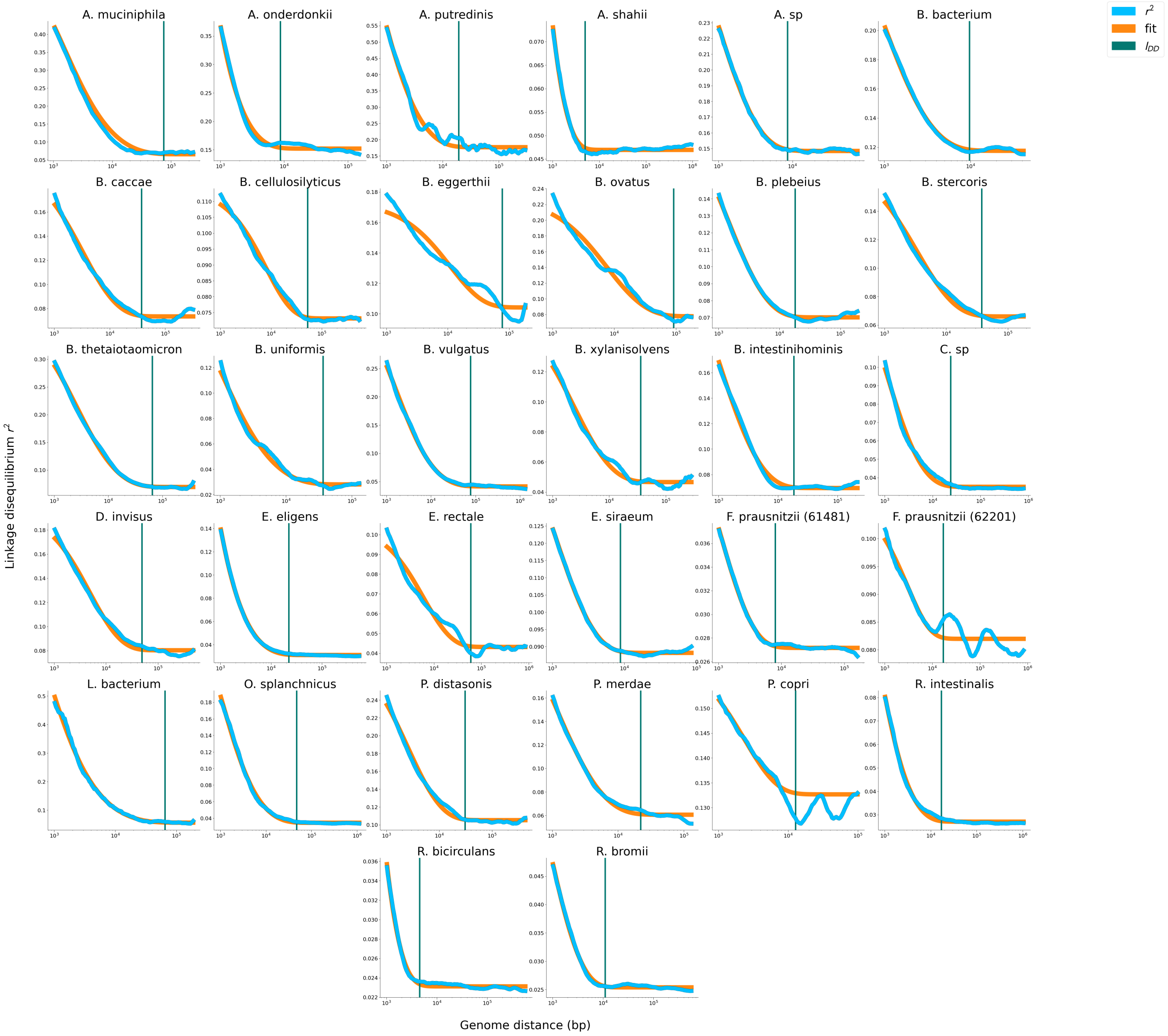

**Figure S26: Estimates of $l_{DD}$ for common commensal gut microbiota.** Using the procedure outlined in Supplementary Section 5.1, we fit Equation S8 to $r_S^2$, and used the parameters of this fit to infer the decay distance $l_{DD}$ (vertical lines). In general, $r_S^2$ appears to saturate at some long distance constant value in most species, and $l_{DD}$ appears to approximate the length scale over which this saturation occurs.

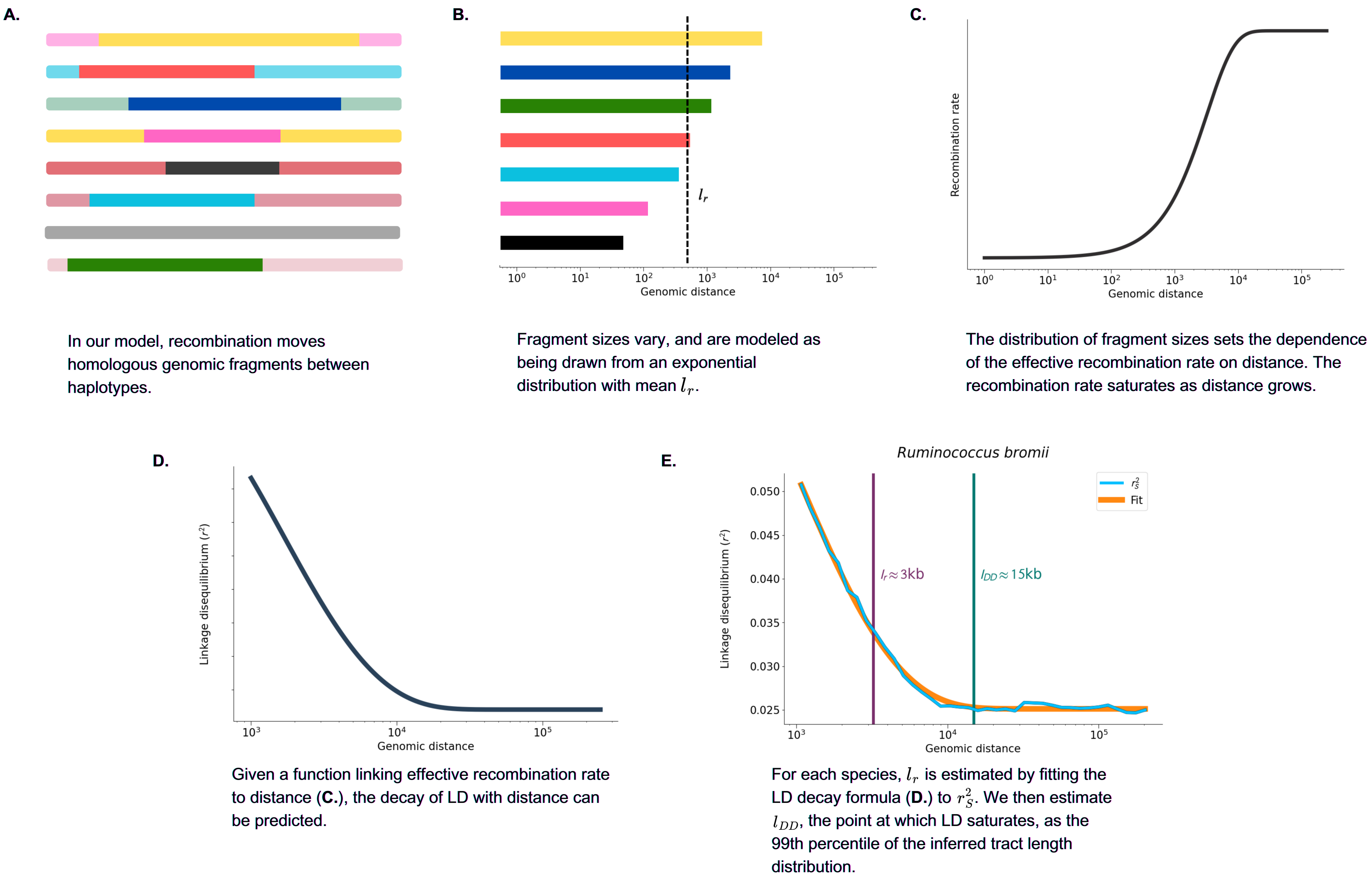

**Figure S27: Schematic of method for inferring $l_r$ and $l_{DD}$.** Visual illustration of the procedure outlined in Supplementary Section 5.1, where Equation S8 is fit to $r_S^2$, and the parameters of this fit to infer the decay distance $l_{DD}$ (in turn used to determine the window size for iLDS) in the microbial species examined here.

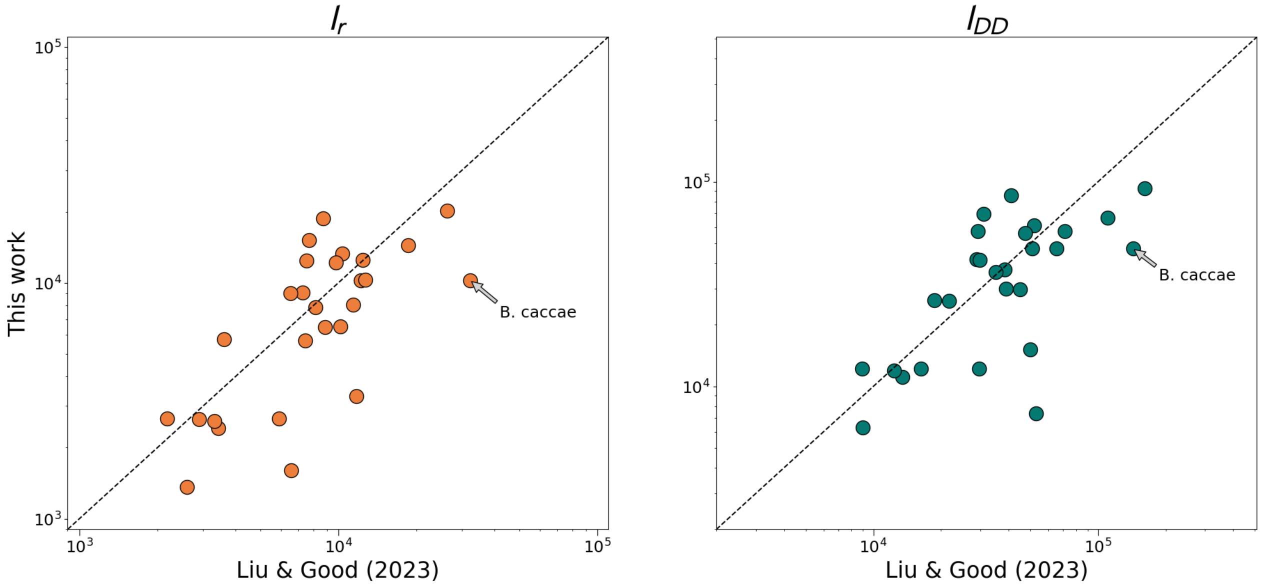

**Figure S28: Comparison of $l_r$ and $l_{DD}$ estimates with Liu & Good (2024).** We compared the values of $l_r$ (left) and $l_{DD}$ (right) obtained using the procedure outlined in Supplementary Section 5.1 with estimates of these values inferred from Supplementary data in Liu & Good (2024) [13]. The dashed line denotes the 1:1 line.

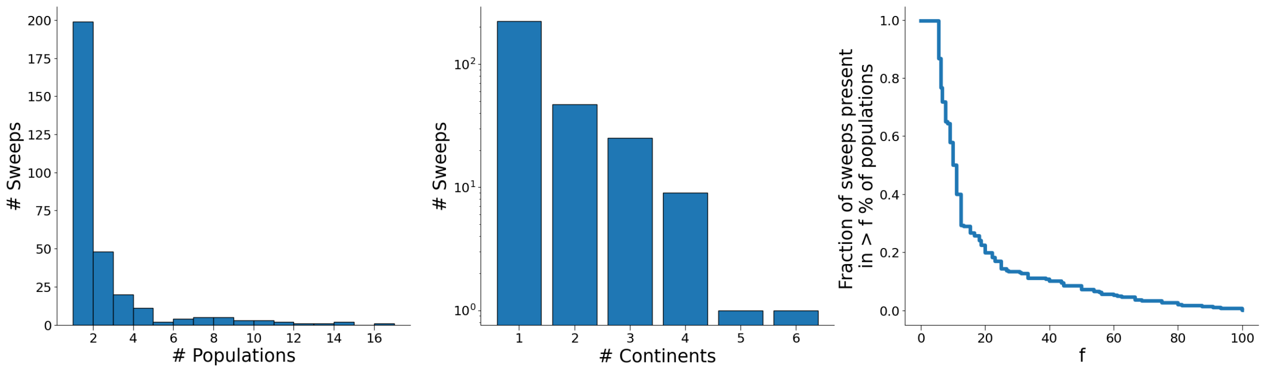

**Figure S29: Spread of sweeps across populations.** Distributions of the number of populations (left) and continents (center) sweeps have spread to. At right, the fraction of sweeps which have spread to at least a given percentage of the total number of populations in which each species was present. In each plot, it is clear that while it is most common for sweeps to be detected in a small number of populations, there are a substantial number of sweeps which have spread between populations and across continents.

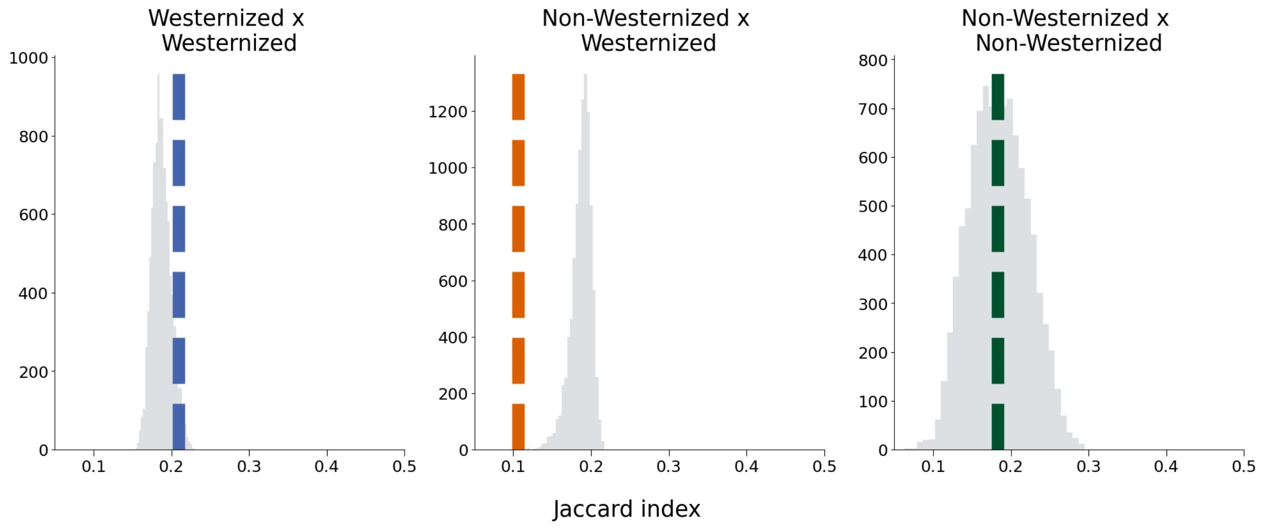

**Figure S30: Jaccard permutation test.** Distribution of Jaccard index after permuting population labels (grey distributions), compared with empirically measured Jaccard index (dashed colored lines). Westernized populations share a greater proportion of sweeps with one another than expected by chance (p-value $= 0.047$). Similarly, Westernized and non-Westernized populations share fewer sweeps than expected by chance (p-value $< 10^{-4}$). See Section 4.3 for further details.

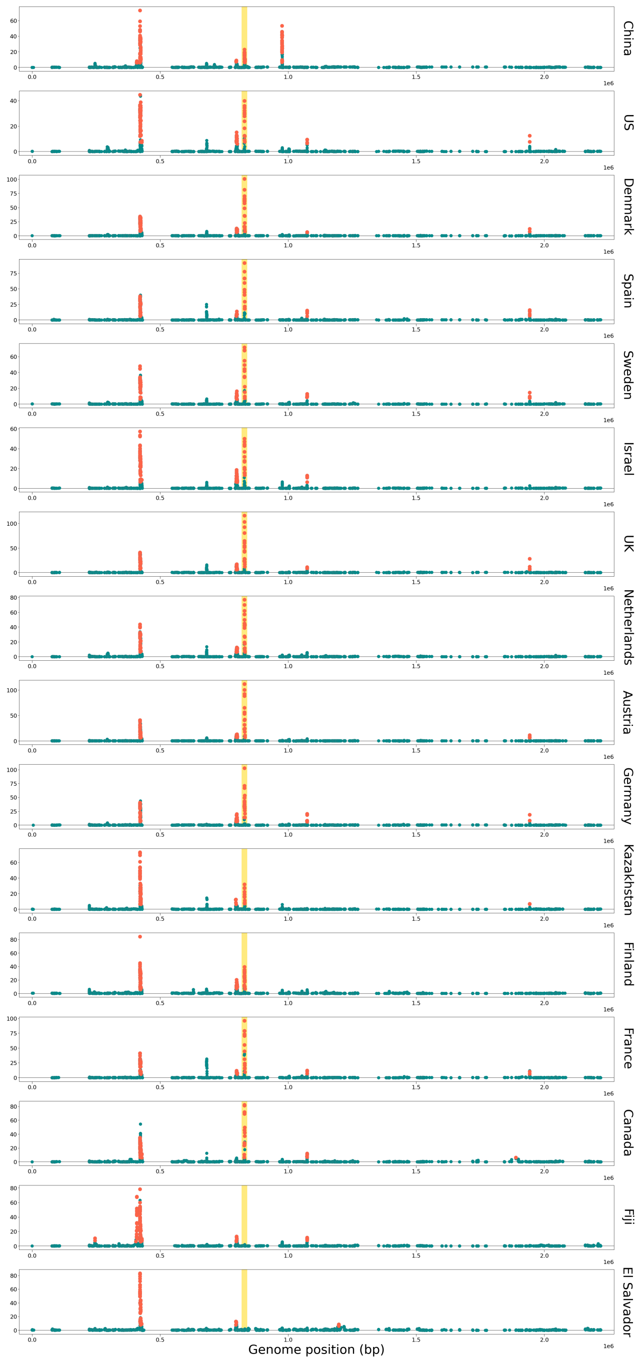

**Figure S31:** ***R. bromii*** **scans across 16 populations.** Expanding on Figure 4C of the Main Text, iLDS scans for all populations of *R. bromii* present in UHGG. As noted in the Main Text, both non-Westernized populations (Fiji and El Salvador—bottom) lack the sweep at the *mdxEF* locus (highlighted in gold), which is present in 14/14 Westernized populations. Note that the the order of the contigs present in the reference genome of *R. bromii* in UHGG differed from that of the reference genome in the MIDAS database [14] to which metagenomes were mapped to from HMP, Qin *et al.*, Korpela *et al.*, and Xie *et al.* For ease of visualization and compatibility with Main Text **Figure 3C**, we have matched the ordering of the contigs in UHGG to that of the reference genome in the MIDAS database for these iLDS scans.

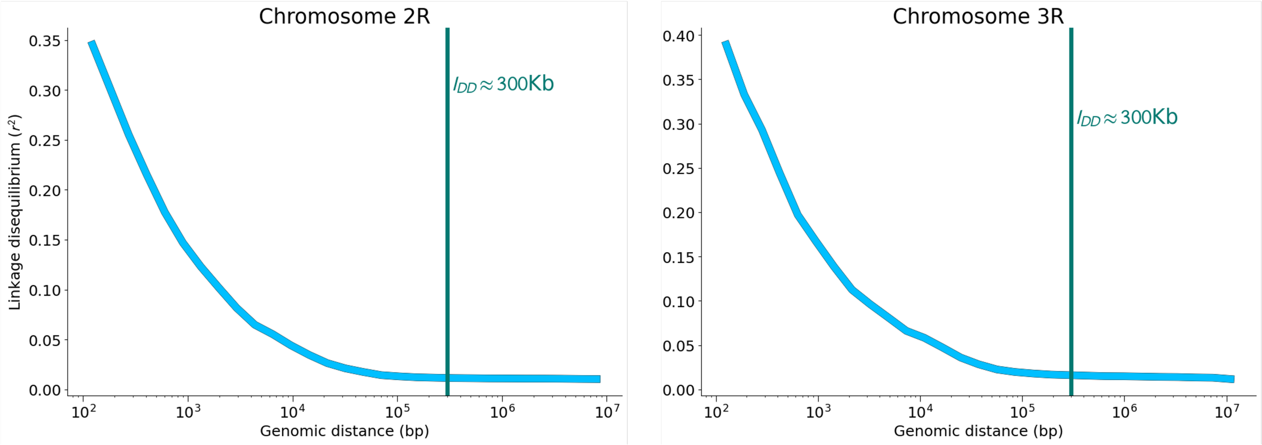

**Figure S32: Decay of LD among common variants in *D. melanogaster*.** Common variant $r_S^2$ curves and $l_{DD}$ points used for iLDS scans for *D. melanogaster* chromosomes 2R and 3R. As recombination in *D. melanogaster* is governed by crossing-over rather than tract exchange, $l_{DD}$ was estimated visually as the approximate point at which LD had fully decayed, rather than by using the scheme described in Section 5.1. As with the scans in bacteria, the window size in number of common non-synonymous SNPs was taken to be the median number of such SNPs necessary to cover $l_{DD}$.

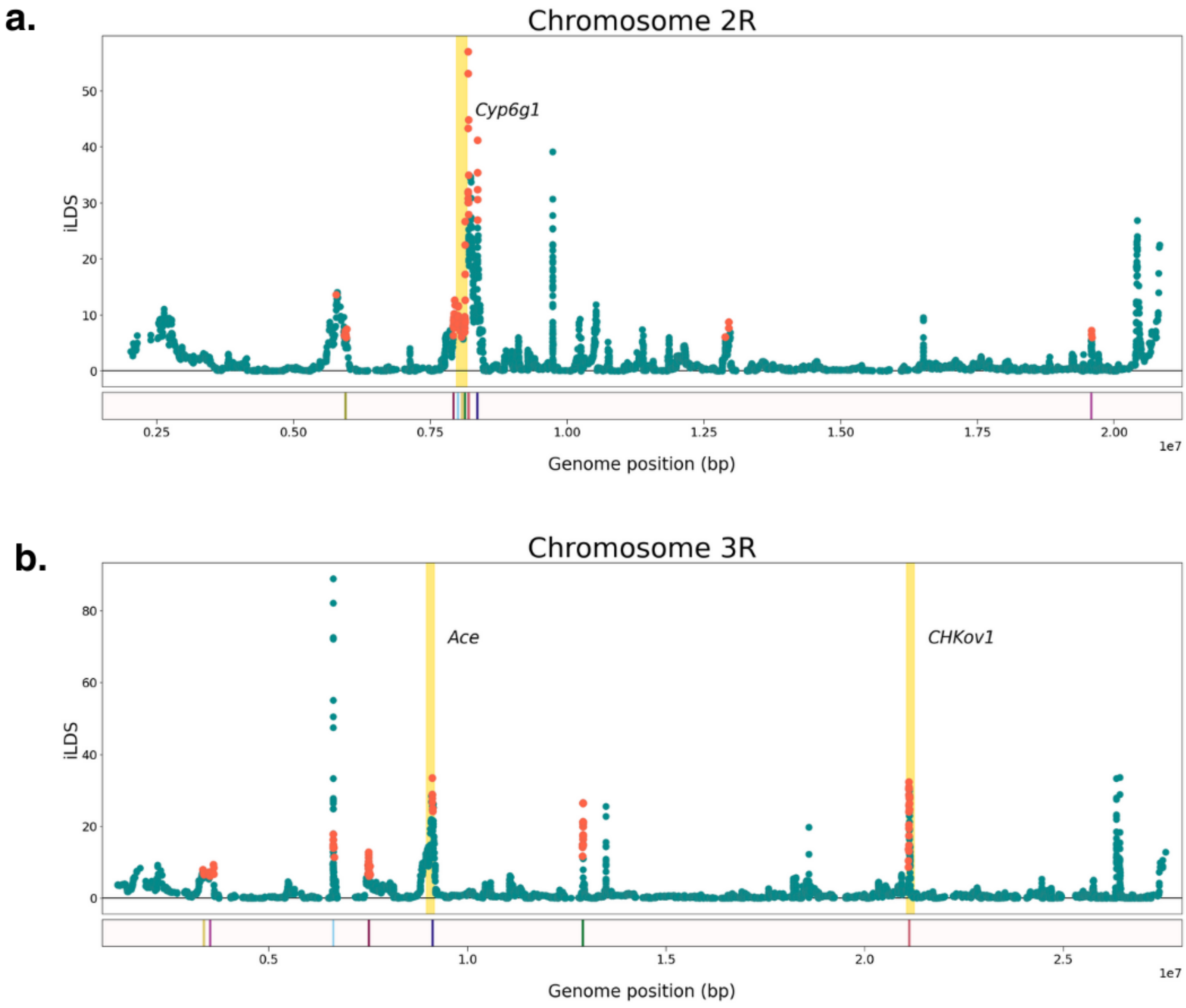

**Figure S33: iLDS scans in *D. melanogaster*.** iLDS scans for chromosomes 2R and 3R of *D. melanogaster*, where $l_{DD}$ was set at 300Kb (Figure **S32**). The location of the yellow vertical line corresponds to the location of known selective sweeps at *Cyp6g1* (2R) and *Ace* and *Chkov1* (3R) plus 50Kb to either side of the locations [15]. Significant windows are shown in orange, while non-significant windows are shown in green. Locations of peaks are shown in the bar, at bottom.

# 1 LD

#### 1.1 LD calculations

To calculate $r^2$ between two loci, we used the standard formula:

$$r^2 = \frac{(f_{AB} - f_A f_B)^2}{(f_A(1 - f_A)f_B(1 - f_B))} \tag{S1}$$

where $f_A$ is the frequency of the minor allele at locus 1, $f_B$ is the frequency at locus 2, and $f_{AB}$ is the frequency of the $AB$ haplotype. $r^2$ measurements were binned by genomic distance (bp) between the two loci, and mean $r^2$ within each bin was computed to determine decay in $r^2$ as a function of distance. We denote the location of the $i^{\text{th}}$ bin to be $d_i$, and the mean $r^2$ within this bin $\langle r_X^2(d_i)\rangle$.

#### 1.2 AUC

To compute AUC($r_X^2$)—the normalized area under the LD decay curve—we first approximate the total area under the curve using the trapezoidal rule and then divide by the total distance horizontal distance of the curve:

$$\text{AUC}(r_X^2) = \frac{1}{d_M - d_1} \sum_{i=1}^{M-1} \frac{\langle r_X^2(d_{i+1})\rangle - \langle r_X^2(d_i)\rangle}{2}(d_{i+1} - d_i) \tag{S2}$$

where $M$ is the total number of bins. The trapezoidal rule was implemented computationally with the trapz function in numpy [16].

#### 1.3 LD confidence intervals

To quantify whether one LD curve is significantly elevated relative to another (e.g. $r_N^2$ versus $r_S^2$), we estimate confidence intervals (CIs) for the more elevated LD curve and assess if area under the other LD curve is lower than the area under the lower CI. Once LD measurements are binned by distance and the mean value ($\langle r^2(d_i)\rangle$) and standard deviation ($\sigma^2_{r^2(d_i)}$) computed for each bin $d_i$, we compute a confidence interval for the sample mean within each bin:

$$\text{CI}(\langle r^2(d_i)\rangle) = \langle r^2(d_i)\rangle \pm z^* \frac{\sigma^2_{r^2(d_i)}}{\sqrt{n_i}} \tag{S3}$$

where the z-score $z^*$ sets the width of the interval used to assess significance, and $n_i$ is the number of measurements in bin $i$. To determine if $r_N^2$ significantly exceeded $r_S^2$, we first computed a lower CI of $r_N^2$ ($r_N^{2\,(LCIB)}$) as follows:

$$r_N^2(d_i)^{(LCIB)} = r_N^2(d_i) - z^* \frac{\sigma^2_{r_N^2(d_i)}}{\sqrt{n_i}} \tag{S4}$$

then calculated AUC($r_N^{2\,(LCIB)}$) and compared this quantity with AUC($r_S^2$). If AUC($r_N^{2\,(LCIB)} - r_S^2$) $> 0$ (that is, if the total area between the lower bound of $r_N^2$ and $r_S^2$ is positive), we considered $r_N^2$ to be significantly elevated above $r_S^2$. When determining if $r_N^2$ exceeded $r_S^2$ at the whole-genome level (Main Text Figure 2B), significance was assessed at the $\alpha = 99.9\%$ level ($z^* = 3.291$). In essence, we assess whether $r_N^2$ overall is greater than $r_S^2$, on average, taking into account the variability of $r_N^2$ at each distance. It is important to note that in order for $r_N^2$ to significantly exceed $r_S^2$, it is neither necessary nor sufficient that $r_N^2$ be greater than $r_S^2$ in all bins. For rare variants, purifying selection is expected to decrease LD among non-synonymous variants relative to synonymous ones, and significance is therefore assessed with respect to the upper bound of $r_N^2$ (i.e. AUC($r_N^{2\,(UCIB)}$) - AUC($r_S^2$) $< 0$.)

To ensure that all species had comparably sized confidence intervals surrounding $r_N^2$ in our genome-wide assessment of significance, we downsampled the number of non-synonymous LD measurements to 1,000,000 for common variants (roughly 1,500 $\times$ 1,500 pairwise measurements) and 25,000,000 (roughly 7,000 $\times$ 7,000) for rare variants. Additionally, as the circular genome size in the species studied here varies, we only considered LD measurements of variants lying on the same contig and within 250Kb of one another. LD was binned in 20 logarithmically spaced bins.

### 2 Simulations

To evaluate the conditions under which LD between non-synonymous variants might be elevated relative to LD between synonymous variants, we performed evolutionary simulations in SLiM 4.0 [17] using custom recipes (Figures **S1 - S6**).

While the species examined in the empirical portion of this study are haploid (with the exception of *D. melanogaster*, Section 7 below), simulations were performed with diploids. Diploid evolution was chosen as a simulation paradigm for two reasons. First, diploid simulations in SLiM employ a standard Wright-Fisher models of evolution, aiding in the interpretability of the simulation results. Second, we aimed to maximize computational efficiency for the large number of simulations we performed ($4500$ burn-in simulations, $135,000$ sweeps, $32,500$ contractions) with population sizes of $N = 10^4$.

While haploid and diploid evolution differ in important ways, simulations were tailored to minimize the effect of these differences, while still retaining the performance advantages offered by simulations of diploidy in SLiM. First, all mutations were assigned a dominance coefficient of 1, making them fully visible to selection regardless of the presence or absence of any mutation on the opposite chromosome. In particular, this means that deleterious variants are always exposed to selection, as is the case with haploids. Second, while the crossing-over recombination typical of diploids differs from the homologous donation of tracts typical of bacterial evolution (at least in the core genome [13, 18]), for our purposes these differences manifest primarily in the efficiency with which genomic elements at large distances can be unlinked. Variants which are physically close to one another relative to the average recombined tract length $l_r$ will be affected identically by crossing-over versus homologous recombination (see Section 5.1). As individuals in our simulated populations have chromosomes of length 10Kb, the decay of linkage should resemble that of a bacterial species with a tract length considerably larger than 10Kb, such as *B. vulgatus* ($l_r = 23$Kb).

To calculate LD, we first sampled 100 individuals from each simulated population. As with our genome-wide assessments, we next sampled a constant number of LD measurements between non-synonymous variants in our simulations to ensure that confidence intervals around $r_N^2$ were comparable across evolutionary scenarios. Specifically, for common variants, we sampled 1250 LD measurements between non-synonymous variants with replacement, for an average of one measurement per simulation for sweeps and contractions. Among rare variants, we sampled 1,000,000 LD measurements. We found, in general, that LD among rare variants was more variable (less smooth decay, wider confidence intervals) than LD among common variants for a given number of LD measurements, likely reflecting the more limited range of potential LD values that can be taken on between pairs of rare variants. Pairs of singletons, for instance, may only take on $r^2$ values of zero or one.

If fewer than 100 LD measurements were available for a given parameter combination and mutation type, we did not calculate an LD decay curve (e.g. among non-synonymous variants for $s_B = 10^{-5}, s_D = 10^{-1}$, where the adaptive variant was always the only intermediate frequency non-synonymous variant following the sweep.) Following sampling, $r_N^2$ and $r_S^2$ were calculated using 8 bins each containing an equal number of non-synonymous variants. Significance was assessed in these simulations at the $\alpha = 99.9\%$ level, again mirroring the genome-wide measurements.

#### 2.1 Burn-in/mutation-selection balance

In each replicate population of size of $N = 10^4$ individuals, burn-in proceeded for $10N$ generations, allowing the population to reach mutation-selection balance. Populations were constituted of individuals with diploid genomes of size $L = 10^4$ bp. Within each simulation, $s_D$ was held constant at all non-synonymous sites, and between simulations, $s_D$ was varied from a nearly-neutral regime ($N_e s_D << 1$: $s_D = 0, -10^{-5}, -10^{-4}$) through a weakly deleterious regime ($N_e s_D \approx 1$ : $s_D = -10^{-3}$ to a strongly deleterious regime ($N_e s_D \gg 1$: $s_D = \{-10^{-2}, -10^{-1}\}$). Following the degeneracy of the codon table, we assumed a ratio of 2.31 non-synonymous sites per synonymous site in our simulations [19], with a per-basepair mutation rate $\mu = 10^{-6}$. Finally, the scaled recombination rate $\rho/\mu$—i.e. the quotient of the recombination and mutation rates—was varied from 0.1 to 1 to 10 ($\rho = 10^{-7}, 10^{-6}, 10^{-5}$). For each parameter combination, 250 replicate populations were simulated.

#### 2.2 Demographic contractions

To simulate demographic contractions, we began by first reading in a burn-in population with the desired $s_D$. In short, sharp contractions, the population size was reduced from $N = 10^4$ to $N = 10^3$, a 90% reduction, and the simulation was allowed to proceed for 20 generations. In the long, shallow contractions, the population size was reduced from $N = 10^4$ to $N = 5 \times 10^3$, a 50% reduction, and the simulation was then allowed to proceed for 5000 generations. For each burn-in population, five replicate contractions were performed, yielding a total of 1250 contraction simulations for each parameter combination.

#### 2.3 Partial sweep

To simulate partial sweeps, we began by first reading in a burn-in population with the desired $s_D$, and then introducing a novel non-synonymous mutation at position zero on the chromosome. For each burn-in population, five replicate sweeps were performed, yielding a total of 1250 simulations per ($\rho/\mu$, $s_B$, $s_D$) combination. Simulations proceeded until the focal, adaptive mutation reached 50% frequency, and were restarted from the burn-in state if the focal variant was lost from the population.

### 3 Metagenomic pipeline

#### 3.1 Overview

In this work, we identify signatures of selection primarily in haplotypes obtained from publicly available fecal shotgun metagenomes of healthy Western adults. While such short read data can yield very accurate estimates of intra-sample allele frequencies, it inherently fragments the relatively long-range LD patterns which are of interest to us. To be able to measure linkage disequilibrium on length scales exceeding that of the length of a short read, we first map reads to reference genomes, and then identify particular sample/species combinations in which a dominant haplotype can be extracted with high confidence and minimal probability of error (Section 3.4). Below, we describe the pipeline used to reconstruct these haplotypes from this metagenomic data.

#### 3.2 Data

The raw sequencing reads for the 1,013 metagenomic samples used in these study were collected from 693 unique individuals (though only a single sample was used per person), including 250 individuals from the Human Microbiome Project (HMP) [20], a further 250 from Xie *et al* (2016) [3], 185 from Qin *et al* (2012) [4], and 8 from Korpela *et al* (2018) [5]. Metadata for each sample is available in Table S1.

#### 3.3 Estimation of species, gene, and SNV content of shotgun metagenomic samples

Species abundances, gene, and SNV content for each metagenomic shotgun sample were previously quantified in Garud & Good (2019) [1] using MIDAS (Metagenomic Intra-Species Diversity Analysis System, version 1.2, downloaded on November 21, 2016) [14]. Here we briefly summarize again the steps followed to obtain the data relevant for this work.

##### *3.3.1 Estimation of species content*

Species abundances for each sample were estimated by mapping reads to a set of single copy marker genes belonging to the 5,952 species in the MIDAS database. A species was considered present if it had an average marker gene coverage $\geq 3$. Since some hosts from the Human Microbiome Project dataset were sampled over multiple timepoints, we next determined a single reference database for each host by including all species present at one or more timepoints with coverage $\geq$3. In doing so, we were as inclusive as possible of any species present at any time point to avoid 'read donating' to species that may resemble true species that are present, but also selective enough to prevent 'read stealing' from a species truly present.

##### *3.3.2 Estimation of CNV content*

CNV content was next estimated by mapping reads to the pangenome for each species in each per-host reference database using Bowtie 2 [21] with default MIDAS settings (local alignment, MAPID$\geq$94.0$\%$, READQ$\geq$20, and ALN_COV$\geq$0.75). MIDAS estimates the copy number of any gene ($c$) as the ratio between its coverage and the median single-copy marker gene coverage.

For each species, we identified "core genes", defined as genes present in at least 90$\%$ of samples belonging to the largest clade (described in the Main Text and in Garud & Good (2019) [1]). Copy number values were used to estimate the prevalence of genes across hosts, defined as the fraction of samples with copy number $c \leq 3$ and $c \geq 0.3$ (conditional on the mean single gene marker coverage being $\geq 5\times$). In our calculation of LD, as well as for any iLDS scans, we only considered core genes. Since genes that are shared across species boundaries can result in read 'stealing' and 'donating', we excluded those genes part of a 'blacklist' of shared genes that we previously inferred [1]. In addition, since some genes may be absent from the MIDAS database, we also excluded genes with $c \geq 3$ in at least one sample in our cohort as in Garud & Good (2019) [1], to avoid examining genes that may have high copy number due to their being present in multiple species.

##### *3.3.3 Estimation of SNV content*

Next, to estimate SNV content, reads were mapped to a single reference genome per species previously selected in the default MIDAS software. Read mapping was performed with Bowtie 2 [21] using default MIDAS thresholds (global alignment, MAPID$\geq$94.0$\%$, READQ$\geq$20, ALN_COV$\geq$0.75, and MAPQ$\geq$20). As per the defaults in MIDAS, species for a given sample were excluded from further analysis if $\leq$ 40$\%$ of their genome had any coverage or if they had median read coverage $\bar{D} < 5$ at protein coding sites with nonzero coverage. To further avoid read stealing and donating, sites were masked in a given sample if $D < \bar{D}/3$ or $D > 3\bar{D}$, as these sites harbor coverage anomalously low or high compared to the genome-wide average coverage $\bar{D}$. An additional coverage threshold requirement of 20 reads/site was imposed for calling SNVs for analyses below.

#### 3.4 Quasi-phasing

Once SNVs had been determined within each sample, we obtained "quasi-phased" (QP) haplotypes from individual hosts following the procedure originally outlined in Garud & Good *et al.* (2019) [1]. Briefly, quasi-phasing estimates one of the dominant haplotypes for a given species in samples for which that species has high coverage ($\bar{D} > 20$) and relatively few intermediate frequency variants, allowing each allele to be confidently assigned to a single haplotype with low (and bounded) probability of polarization errors due to sampling noise. As our paper and Garud & Good (2019) analyze the same data, we leveraged the same sample x species classified as quasi-phaseable in that work. Any sites with intermediate frequency ($0.2 < f < 0.8$) polymorphism in the sample were treated as missing data, and the dominant major alleles ($f \geq 0.8$) at all sites passing the site-specific coverage thresholds outlined above were taken to constitute the haplotype. Further methodological details on QPing can be found in Garud & Good (2019). In total, our analyses were performed on 2641 QP haplotypes belonging to the largest clade of each of species considered, as seen in Figure **S13**.

#### 3.5 Inclusion criteria

Once samples had been quasi-phased, we included only species which had a sufficient number of non-closely related ($d > 5 \times 10^{-4}$) haplotypes and had reference genome in MIDAS that was not fragmented into a large number of contigs. In particular, we included all species which had at least 20 non-closely related samples and an N50 > $10^5$ bp. In total, 31 species passed these thresholds.

#### 3.6 Population structure

Several of the gut microbial species examined here exhibit strong population structure [1, 13, 22], with 'clades' of lineages that recombine far more frequently with one another than with lineages belonging to other clades. In species with strong clade structure, LD is elevated genome-wide and shows comparatively little decay with distance [1, 13]. Genetic diversity is therefore dominated by fixed differences between clades, masking signatures of ongoing recombination within clades. To control for population structure, we analyzed LD only among lineages belonging to the largest clade (as identified previously in [1]) of each species.

Additionally, we controlled for the potential effects of elevated LD due to the presence of occasional closely related lineages, as measured by the per base-pair divergence $d$ at fourfold degenerate sites (that is, sites where a nucleotide difference does not result in an amino acid change). Pairwise divergence $d$ was determined by first calculating the number of fourfold degenerate sites at which two haplotypes differed, and then dividing this number by the total number of such sites in the core genome. While on average random pairs of lineages belonging to the same clade have $d$ on the order of $10^{-2}$/bp, unrelated hosts occasionally harbor very closely related lineages, with pairwise $d$ two orders of magnitude smaller than typically observed across hosts [1, 13]. To avoid elevations in LD arising from these anomalously closely related lineages, we choose only a single representative for any group of lineages $d < 5 \times 10^{-4}$, as in our previous analysis [1].

### 4 UHGG

To augment our analyses using quasi-phased genomes, we downloaded alignments from Unified Human Gastrointestinal Genome collection [6]. We wrote custom scripts to annotate each SNV falling in a coding region as synonymous or non-synonymous by using coordinates for the reading frame. Metadata for these samples, including accession numbers, can be found in Table S5.

Next, following the pipeline developed for metagenomic samples, we identified top-level clades for each species we analyzed in UHGG by creating a UPGMA dendrogram of all samples using $d$ at four-fold degenerate (4D) sites as a distance metric. Four-fold degenerate sites are sites at which all single nucleotide mutations yield the same amino acid. We then manually cut the dendrogram if there appeared to be multiple deep clades. As with HMP, we focused our subsequent analyses only on samples belonging to the largest clade identified, and retained only a single representative from each group of closely related samples ($d < 5 \times 10^{-4}$/bp).

For our analyses using UHGG data to identify peak overlap between populations (Figure 4, Main Text) we only used genes which were core to all continents for which a species was present. We took a gene to be core if 90% of polymorphic sites were present in 90% of non-closely related samples. Thus, our analyses only focus on genes which are truly present in all populations surveyed.

Species composition varies considerably between Westernized and non-Westernized microbiomes. We chose to focus only on species where 20 or more non-closely related ($d > 5 \times 10^{-4}$) at 4D sites, as in our analysis with HMP) genomes were present in at least one Westernized and one non-Westernized population. As Westernized populations are far more extensively sampled, these populations tended to have considerably more genomes for a given species than did any non-Westernized population. Therefore, to ensure that each iLDS scan had the same sensitivity, we downsampled so that for each species, all populations had the same number of genomes.

For the pathogens analyzed (*C. difficile* and *H. pylori*), we did not make comparisons in the locations of peaks between continents, and therefore ran our analyses on all sites which were present in 90% or more of non-closely related samples, rather than defining core genes. Because these pathogens had large numbers of isolates ($\mathcal{O}(10^2)$), we restricted our analyses only to isolate genomes to further minimize any source of mapping error.

#### 4.1 Virulence factors

To identify virulence factors in *C. difficile* (Figure 3A, Main Text), we performed a blast search of its reference genome assembly from UHGG against the VFDB database [23]. We annotated any *C. difficile* gene as a virulence factor if it had more than 90% amino acid identity across $\geq$ 333 codons with a known VFDB virulence factor.

#### 4.2 Identifying Westernized samples

To identify a sample as belonging to a Westernized or non-Westernized population, we relied on the annotations available in the curatedMetagenomicData package [24], which provides a range of metadata on publicly available human gut microbiome metagenomic data. All samples from Fiji, Tanzania, Madagascar, Peru, and El Salvador were identified as non-Westernized, while only a subset of samples from Mongolia were identified as non-Westernized. We excluded all Mongolian samples identified as Westernized from analysis.

#### 4.3 Jaccard index

To quantify the overlap in number of peaks between cohorts, we calculated the Jaccard index between all pairs of populations. The Jaccard index measures the overlap between two sets, and is the ratio of the size of the intersection of the sets to the size of the union of the sets.

$$J = \frac{\text{\# elements shared}}{\text{\# elements total}} \tag{S5}$$

To determine the Jaccard index for a pair of populations, we first calculated the number of peaks shared and the number of peaks in total for each species. To determine a mean Jaccard index across Westernized populations, for example, we took the sum of the number of peaks shared across all species and all pairs of Westernized populations, and divided this quantity by the total number of peaks considered for each pair of populations. As the precise non-synonymous variants detected as significant by iLDS may vary, we counted peaks as shared between populations if a variant within one of the peaks was within 1000bp of a variant in the other, and not shared otherwise.

Next, to determine whether the value of the Jaccard index between each cohort was statistically significant, we performed a permutation analysis. First, we randomly shuffled the Westernized and non-Westernized status. Then, we repeated the Jaccard index calculation as described above 10^4 times. Finally, to obtain a p-value, we compared the true value of J for each pair of groups with the distribution of values of J for that pair, taking the p-value to be the frequency of permuted values more extreme than the observed value. Because the groups are of different sizes—with the Westernized/Westernized group containing nearly 10 times as many measurements as the Non-Westernized/Non-Westernized group—the permuted distributions have different variances (though identical means). Histograms showing the distributions of permuted values relative to the true values of the Jaccard index can be found in Figure **S30**

### 5 iLDS

#### 5.1 Determining the window size for computing iLDS

To perform an iLDS scan, LD is calculated in sliding windows across the genome, and LD measurements within each window subsequently binned by genomic distance between variants, so that AUC($r_N^2$), AUC($r_S^2$), and AUC($r^2$) within each window may be calculated. To ensure that all windows contained both an adequate and comparable number of synonymous and non-synonymous variants with which to calculate $r_N^2$ and $r_S^2$ curves, windows were defined in terms of SNPs rather than base-pairs. Specifically, each window was defined by the region spanning $n$ consecutive non-synonymous, intermediate frequency SNPs. The value $n$ was chosen such that the average size, in base pairs, of all windows corresponded as closely as possible to the distance over which LD fully decays genome-wide for that species. To determine AUC($r^2_{genome-wide}$) in each window, we first downsampled to 1,000,000 LD measurements total, and then binned these measurements using the bins defined for that window.

To infer the distance over which LD decays, we model recombination as moving tracts of DNA from a donor genome into a recipient genome, where they replace homologous segments in a process known as homologous recombination in bacteria. While bacteria exchange genomic material in a variety of ways, including through the transfer of mobile genetic elements such as plasmids, homologous recombination is thought to be important in breaking up LD in the core genome, which is the focus of this work [13, 18]. The size of these homologously exchanged tracts vary, but numerous experimental studies have found that, within a species, tract lengths are approximately exponentially distributed [25, 26, 27, 18]. The speed at which loci are unlinked therefore depends crucially on the average size of these recombinant fragments ($l_r$), which in turn characterizes the total (exponential) distribution of tract lengths. In order to obtain the length scale on which LD decays ($l_{DD}$), we will first obtain an estimate of $l_r$, the average tract length.

In our model, two loci recombine when one or the other of the sites, but not both simultaneously, is homologously replaced by an imported fragment. Assuming a uniform per base-pair rate $r$ at which a recombination event may begin at each site along the genome, and an exponential distribution of tract lengths with average length $l_r$, the effective recombination rate $R$ between loci separated by a distance $l$ is

$$R(l) = rl_r(1 - e^{-l/l_r}) \tag{S6}$$

This functional form suggests that the effective recombination rate increases linearly as $\sim rl$ at short distances (specifically, when $l \ll l_r$), and ultimately saturates to a single value $rl_r$ for pairs of variants separated by $l \gg l_r$. The transition between these two regimes occurs when $l \approx l_r$.

In a neutrally evolving, panmictic population of size $N$, a classic result [28] states that expected LD obeys the following relationship:

$$E[r^2] = \frac{10 + 2NR}{22 + 26NR + 4(NR)^2} \tag{S7}$$

where $R$ is the effective recombination rate between sites, provided this expectation is taken with respect only to intermediate frequency alleles (e.g. $0.1 < f < 0.9$ [28]). The ultimate cause of the decay of LD is the dependence of the effective recombination rate $R$ on genomic distance—in our model, this relationship is given by Equation S6. We should in principle be able to estimate the parameters $N$, $r$, and $l_r$ by fitting Equation S7 to the empirical LD decay curves for intermediate frequency synonymous variants ($r_S^2$). To obtain an estimate of the effective distance over which LD fully decays—and thereby a window size for iLDS—we are primarily interested in determining $l_r$.

While Equation S7 describes the behavior of $E[r^2]$ over all distances, its value is, in theory, independent of $l_r$ over short distances ($l \ll l_r$) as $R(l)$ is itself independent of $l_r$ in this regime. Therefore, as our goal is first to estimate $l_r$, we disregard short-distance LD and focus only on fitting when $l \approx l_r$ and greater. Using a completely different methodology, a previous study investigating the dynamics of recombination in human gut bacteria (Liu & Good (2024)) estimated $l_r$ for many of the species under consideration here, and obtained values which were $O(10^3) - O(10^4)$. Therefore, we restrict our attention to $l > 10^3$ bp.

In particular, we fit our $r_S^2$ curves with the following model:

$$E[r^2(l)] = C\frac{10 + 2NR(l)}{22 + 26NR(l) + 4(NR(l))^2} \tag{S8}$$

where $C$ is a normalization constant allowing us to fit the relative change in $r_S^2$ with distance, rather than absolute values of LD strictly. Fits were obtained using the scipy.optimize.curve_fit function in scipy [29].

To assess the performance of our model visually, we show the empirical $r_S^2$ and the fit obtained from Equation S8 in Figure **S26**. Overall, we see that the theoretical prediction (Equation S8) provides an excellent fit of LD across several orders of magnitude in the vast majority of species.

Having estimated $l_r$, we next move to the main goal of inferring $l_{DD}$, the distance over which LD fully decays genomewide. In our model of homologous recombination, the effective rate of recombination (and by extension, the level of LD) saturates in the $l \gg l_r$ limit of Equation S6—that is, as the distance between variants becomes large relative to the size of a tract that can be horizontally transferred. To infer this distance, we took the value of $l_r$ estimated above as the mean of an exponential distribution of tract lengths for each species. We then took the $99^{\text{th}}$ percentile of this distribution to be our estimate of $l_{DD}$.

As expected, the decay of LD appears to saturate to a constant value at large distances in all species examined (Figure **S26**). Moreover, by visual inspection, our estimated values of $l_{DD}$ furnish a good approximation for this distance. The estimates of $l_r$ and $l_{DD}$ can be found in Table S3.

Next, to determine the window size $n$ given $l_{DD}$, we calculated the median number of intermediate frequency non-synonymous SNPs spanning this distance. In general, this window size depends on both $l_{DD}$ and the density of common non-synonymous SNPs, with larger $l_{DD}$ and higher densities leading to greater $n$.

Finally, once the window size has been assessed, we determined the number of bins $b$ used to bin LD within each window. We set $b$ such that each bin would contain approximately 50 LD measurements—that is, $b = \binom{n}{2}/50$. To avoid creating windows with either very few bins (which obscures the distance dependence of LD within a window) or very many bins (which reduces statistical power), we set a minimum bin number of 3 and a maximum bin number of 10. Window sizes and bin numbers for each species are summarized in Table S3.

###### *5.1.1 Comparison of tract length estimates with previous work*

To assess the performance of our method for estimating the mean tract length ($l_r$) and the $99^{\text{th}}$ percentile of the tract length distribution ($l_{DD}$), we compared our inferences of these quantities with those of Liu & Good (2024) [13]. The authors of this prior work explored the dynamics of homologous recombination in the same cohort of metagenomic samples examined here, employed QPing to obtain sample-specific haplotypes, and clade-controlled in an identical fashion. While the method developed by Liu & Good (2024) detects recent transfers using pairwise comparisons between closely related strains, we infer statistical properties of the recombination process using the decay of LD among all non-closely related strains. Our approaches are complementary, and below we show a broad agreement between our estimates.

Liu & Good (2024) identified $\approx$250000 unique instances of recombination—that is, the horizontal exchange of a genomic tract—across 29 species. The tract lengths for each these instances were provided in their supplemental Table S3. For each species analyzed in both datasets (28/29), we then computed the mean tract length ($l_r$) and the $99^{\text{th}}$ percentile of the tract length distribution (corresponding to our $l_{DD}$) from the Liu & Good (2024) empirical observations, and compared these quantities with those we obtained using our LD based method.

Overall, we found a correspondence between the two methods (Figure **S28**). Across all shared species, we found a correlation coefficient $r^2$ of 0.58 for $l_r$ and 0.6 for $l_{DD}$ between our approaches. The species for which our differing methodologies conflict most strongly is *Bacteroides caccae*, which Liu & Good (2024) [13] noted appears to have an anomalously low rate of recombination and large number of closely related lineages. Removing this single species increases the correlation coefficient $r^2$ for $l_r$ to 0.7 and $l_{DD}$ to 0.65.

#### 5.2 Identification of analysis windows with significant iLDS values

iLDS was applied to the data in analysis windows as defined above. As stated above, iLDS was calculated in sliding windows across the genome, with each window centered around a single intermediate frequency non-synonymous SNVs. iLDS was considered to be significant in a given analysis window if both AUC($r^2_N$) significantly exceeded AUC($r^2_S$) within the window, and overall AUC($r^2_{\text{local}}$) within the window significantly exceeded the genome-wide expectation for identically sized regions AUC($r^2_{\text{genome-wide}}$). Significance was determined by constructing confidence intervals (as described in the LD Confidence Interval section, above) around $r^2_N$ and $r^2_{\text{local}}$ and checking if their lower CIs exceeded $r^2_S$ and $r^2_{\text{genome-wide}}$, respectively, at the $\alpha = 95\%$ level.

#### 5.3 Evaluating the performance of iLDS in simulations

We assessed the ability of iLDS to correctly identify selective sweeps, as well as the rate at which it misclassified variants not under positive selection as selected. To assess the potential for misclassifying non-sweeps as sweeps, we simulated constant Ne populations and two demographic contractions. To assess the ability of iLDS to correctly identify sweeps in a range of evolutionary parameters, we simulated sweeps with varied selection strengths as well as the number of generations since selection ceased to capture the decay of a sweep via mutation and recombination events over time.

We used the forward simulation engine SLiM to simulate populations at mutation-selection balance and selective sweeps [17]. These simulations are similar to those described in Supplementary Section 2, but are 500Kb rather than 10Kb, allowing us to assess iLDS's ability to detect sweeps of various strengths that have had the opportunity to sufficiently decay to roughly neutral levels at the edges. We simulated populations with an initial size of $10^4$ individuals. As in the simulations outlined in Supplementary Section 2, we assumed a ratio of 2.31 non-synonymous sites per synonymous site, with a per-basepair mutation rate $\mu = 10^{-6}$ and recombination rate $\rho = 10^{-6}$. For each evolutionary scenario described below, we simulated 1000 replicate populations. From each replicate population, we sampled 100 individuals, and calculated the value of iLDS based on this sample.

Based on our analyses in Supplementary Section 2, we performed these simulations in a parameter regime where we expect iLDS to be effective in detecting selective sweeps: that is, when non-synonymous variants are under weak purifying selection such that $s_d < s_b$. If purifying selection is very strong, we expect that few deleterious variants to hitchhike along with the focal adaptive variant during a sweep. Conversely, if purifying selection is weaker than drift, we do not expect differential patterns of LD between synonymous and non-synonymous variants to emerge during selective sweeps (Figure **S5**). In testing iLDS, we therefore modeled all non-synonymous variants (with the exception of the sweeping variant) as deleterious, with a single selection coefficient $s_D = -10^{-3}$ (i.e. $N_e|sD|$= 10).

Before simulating selective sweeps or demographic contractions, we allowed each population to equilibrate to mutation-selection balance for $10N_e$ generations. Below we describe selective sweep and demographic contraction simulations.

###### *5.3.1 Selective sweeps*

We simulated selective sweeps of strength $s_B = 0.01, 0.05$, and $0.1$, beginning after the population had reached mutation-selection balance. The adaptive variant was seeded at the midpoint of the genome. Simulations were restarted if the variant was lost from the population prior to reaching 50% frequency. After the beneficial variant reached a frequency of 50% in the population, the selection coefficient of the focal variant changed from beneficial to neutral. To assess the impact of aging on the ability of iLDS to detect sweeps, the sweep was allowed to decay for 0, 10, 100, 500, and 1000 generations.

###### *5.3.2 Demographic contractions*

As in Supplementary Section 2, we simulated two kinds of contractions: short, sharp contractions (90% reduction in population size for 20 generations) and long, shallow contractions (50% reduction for 5000 generations). These population size changes were introduced to populations equilibrated to mutation-selection balance.

##### *5.3.3 Calculating iLDS*

iLDS is applied in windows of a fixed number of intermediate frequency ($0.2 < f < 0.8$), non-synonymous SNPs. To identify the appropriate number of SNPs to use for each window, we computed the distance at which LD decays ($l_{dd}$) in simulations under mutation-selection balance. This corresponded to ∼7.5Kb, or 10 intermediate frequency non-synonymous SNPs. While sweeps may elevate LD locally in the region of the adaptive variant, we observed in our simulations of sweeps that on average across the whole genome, LD decayed at roughly the same rate as mutation-selection balance.

As expected, LD decays considerably more slowly in the contraction scenarios (∼50Kb). We therefore set the window size to be 40 intermediate frequency non-synonymous SNPs for the populations undergoing contractions, corresponding to this distance.

For each simulation, we first calculated AUC($r_N^2 - r_S^2$) and AUC($r^2_{\text{local}}$–$r^2_{\text{genome-wide}}$) in each window as described in the Main Text:

$$r^2_{\Delta NS} = \text{AUC}(r_N^2 - r_S^2) \quad \text{and} \quad r^2_{\Delta LG} = \text{AUC}(r^2_{\text{local}} - r^2_{\text{genome-wide}})$$

and then normalized these quantities:

$$\bar{r}^2_{\Delta NS} = \frac{r^2_{\Delta NS} - E[r^2_{\Delta NS}]}{\text{std}(r^2_{\Delta NS})} \quad \text{and} \quad \bar{r}^2_{\Delta LG} = \frac{r^2_{\Delta LG} - E[r^2_{\Delta LG}]}{\text{std}(r^2_{\Delta LG})}$$

where $E[r^2_{\Delta NS}]$, $E[r^2_{\Delta LG}]$, std$[r^2_{\Delta NS}]$, and std$]r^2_{\Delta LG}]$ were computed from 1000 simulations at mutation-selection balance, where one value of AUC($r_N^2$–$r_S^2$) and AUC($r^2_{\text{local}}$–$r^2_{\text{genome-wide}}$) was drawn from each simulation at random.

However, for the populations experiencing demographic contractions, $E[r^2_{\Delta NS}]$, $E[r^2_{\Delta LG}]$, std$[r^2_{\Delta NS}]$, and std$[r^2_{\Delta LG}]$ were calculated from the simulations of contractions of the same magnitude.

To assess significance, we applied the criteria as described in Supplementary Section 5.2.

##### *5.3.4 Simulations results*

We plotted receiving operator characteristic (ROC) curves, which compare the true positive rate (TPR) of correctly identifying a selective sweep versus the false positive rate (FPR) of incorrectly classifying a neutral simulation as a selective sweep (Figures **S7** - **S9**). Overall, iLDS is able to identify recent and strong selective sweeps. Specifically, when selection coefficients are sufficiently strong (e.g. 0.05 or 0.1), iLDS has the ability to detect sweeps that have decayed for up to 500 generations. However, when selection coefficients are weak (.e.g. 0.01), iLDS can only reliably detect sweeps that have decayed up to 100 generations in the past. By 1000 generations in our simulations, iLDS typically struggles to distinguish sweeps. The performance of iLDS is marginally worse for populations experiencing bottlenecks, particularly sharp, short bottlenecks—however, this decrement in performance is mainly apparent for weak selection ($s_b = 0.01$, Figure **S8**).

Additionally, we computed false discovery rates (Figures **S10** - **S12**). To do so, we set the critical value to match that of the value applied to data ($\alpha = 0.05$) for the $\chi^2$ test of significance. We also required significant windows to have AUC($r_N^2 - r_S^2$) $> 0$ and AUC($r^2_{\text{local}} - r^2_{\text{genome-wide}}$) $> 0$. We computed the number of true positives identified in a set of 1000 simulations of selection and we computed the number of false positives in a set of 1000 simulations without selection. We found that in scenarios when iLDS has good power to detect selective sweeps, FDR rarely surpasses 10%.

#### 5.4 Peak calling

After significant windows were identified, we next sought to localize individual gene-specific selective sweeps by identifying "peaks" of elevated values of iLDS. To do so, we identified groups of physically proximate and tightly linked intermediate-frequency non-synonymous SNPs in the center of significant windows—that is, groups of non-synonymous SNPs consistent with having been carried together to intermediate frequency on shared recombinant tracts. Each SNP contained within a peak may be involved in the putative sweep as either a driver or passenger mutation.

We defined tight linkage as pairwise $r^2 \geq 0.5$ between SNPs. Among all pairwise LD comparisons between common, non-synonymous variants within distance $l_{DD}$, only 1.3% in R. bromii exceeded $r^2 = 0.5$, with similar proportions observed in other species. Thus, our linkage threshold ensures that variants within a peak are among the most tightly linked over this length scale.

To call peaks, we employed a greedy clustering algorithm which grouped tightly linked ($r^2 \geq 0.5$) and physically proximate (distance $\leq l_{DD}$) central SNPs in significant windows. First, we identified all pairwise comparisons between central SNPs in significant windows which met the linkage and proximity criteria described above. Next, we constructed a graph where nodes were individual central SNPs, and edges were drawn between tightly linked, proximate central SNPs. We then extracted the connected component containing the window with the largest iLDS value from this graph. Only connected components with more than one significant window were considered, otherwise solo significant windows were discarded from further peak-calling analysis. By filtering out clusters made up of only a single significant window, we conservatively ensure that all peaks are supported by multiple windows. While not all windows within a peak need be consecutive to one another, and central SNPs need not be tightly linked to all other variants in the peak, each is connected by an edge to at least one other significant window within the peak.

After identifying the peak with the highest iLDS value, we next removed from further peak calling analysis any significant windows within 25Kb of any variant in the cluster. By doing so, we attempt to identify peaks that comprise independent selective events.

We then repeated this procedure—i.e. drawing our proximity/linkage graph, identifying the connected component containing the window with the highest iLDS value as a peak, and removing windows within 25Kb of subsequent peaks—until all significant windows are either placed within a peak or removed from analysis.

To assign putative functions associated with each potential sweep, we then inspected the gene annotations for all significant SNPs contained in a single peak. We report all such annotations in Table S4.

### 6 Gene enrichment analysis

To assess if particular categories of genes were enriched for signatures of selection, we used gene annotations from the PATRIC database and performed several gene enrichment analyses on different classes of annotations [30, 31]. First, we collated a list of all genes which lay within a peak in any of the 31 species for which we ran a scan (Table S4). Next, we performed separate enrichment analyses on three types of annotations, (i) predicted gene products [30], (ii) enzyme commission numbers [8], and (iii) COG categories [7]. The MIDAS reference database already contained predicted gene product and enzyme commission number annotations from PATRIC [30]. To obtain COG categories for each gene, we annotated the reference genomes in MIDAS using eggnog-mapper [32].

To assess whether a functional category was significantly enriched among selected genes, we performed a Fisher exact test comparing the number of these classes of genes (i.e. (1), (2), & (3), above) detected to be under selection with the number of such genes found among all core genes, and corrected for false discovery using the Benjamini-Hochberg procedure. We assessed significance at the $\alpha = 95\%$ level.

Observed versus expected counts of functional categories among genes under selection, as well as statistical significance, is summarized in Figure **S22**.

### 7 Analyses with *Drosophila melanogaster*

Though the primary focus of this work is detecting selective sweeps in bacteria, hitchhiking of linked deleterious variants with selected sites is a general evolutionary phenomenon that many species experience [33, 34, 35, 36, 37]. If hitchhiking is an important factor elevating $r_N^2$ above $r_S^2$, we expect this signal to manifest in a very broad range of adapting populations. Therefore, in addition to our scans in gut bacteria, we further assessed the capacity of iLDS to recapitulate known sweeps in natural populations of *Drosophila melanogaster*, focusing in particular on three loci implicated in pesticide and virus resistance on chromosomes 2R (*Cyp6g1*) and 3R (*Ace*, *CHKov1*) [15, 38, 39, 40].

We downloaded alignments of 100 *D. melanogaster* genomes from the Drosophila Genetics Reference Panel [41] belonging to the Raleigh population, as well as annotations for all sites. While the method for obtaining a window size for iLDS used for bacteria is not appropriate for diploid species (in which recombination is governed by crossing-over, rather than exchange of tracts), we visually estimated that linkage among common synonymous variants fully decayed by $\approx$300Kb on both chromosomes, and used this distances to determine the window sizes on these chromosomes (Figure **S32**). We found that our scans were sharply peaked around the three known instances of selective sweeps at *Cyp6g1*, *Ace*, and *CHKov1* (Figure **S33**). While the sweeps at *Cyp6g1* and *CHKov1* are both driven by transposable element insertions, the sweep at *Ace* is a result of three single nucleotide mutations in that gene [15, 38]. When we applied our peak-calling procedure, in which groups of significant windows centered around variants which are both tightly linked and physically close to one another are considered a peak (Section 5.4), we found that the peak in chromosome 3R near *Ace* (Figure **S33**) was in fact made up of two variants lying in *Ace* and one other nearby variant. However, the peaks lying near *Cyp6g1* in chromosome 2R and *CHKov1* in chromosome 3R did not contain any SNPs lying in these genes, consistent with these sweeps being driven by transposable element insertions which are not directly visible to iLDS, which computes LD and calls peaks using SNPs alone.

While we only tested iLDS's ability to identify simulated single mutational origin selective sweeps (hard sweeps), the sweeps at *Cyp6g1*, *Ace*, and *CHKov1* are all thought to involve multiple, independent mutational origins (soft sweeps) [15, 38]. The ability of iLDS to successfully identify these soft sweeps may indicate that the statistic has power to identify both hard and soft selective sweeps. Future work testing the capacity of iLDS to uncover selective sweeps under a range of adaptive scenarios—including hard versus soft sweeps—is needed.
